# CCQM-P199: Interlaboratory comparability study of HIV-1 RNA copy number quantification

**DOI:** 10.1101/2024.04.12.589043

**Authors:** Alison S. Devonshire, Eloise J. Busby, Gerwyn M. Jones, Denise M. O’Sullivan, Ana Fernandez-Gonzalez, Sachie Shibayama, Shin-ichiro Fujii, Megumi Kato, John Emerson Leguizamon Guerrero, Claudia Patricia Tere Peña, María Mercedes Arias Cortes, Roberto Becht Flatschart, Marcelo Neves de Medeiros, Antonio Marcos Saraiva, Young-Kyung Bae, Inchul Yang, Hee-Bong Yoo, Alexandra Bogožalec Košir, Mojca Milavec, Lianhua Dong, Chunyan Niu, Xia Wang, Phattaraporn Morris, Sasithon Temisak, Megan H. Cleveland, Peter M. Vallone, Daniel Burke, Michael Forbes-Smith, Jacob McLaughlin, Samreen Falak, Martin Hussels, Rainer Macdonald, Andreas Kummrow, Burhanettin Yalçinkaya, Sema Akyurek, Muslum Akgoz, Maxim Vonsky, Andrey Runov, Clare Morris, Neil Almond, Jim F. Huggett

## Abstract

Infection with human immunodeficiency virus (HIV)-1 leads to acquired immunodeficiency syndrome (AIDS) if left untreated. According to UN figures, approximately 39 million people globally were living with HIV in 2022, with 76% of those individuals accessing antiretroviral therapy. Measurement of plasma viral RNA load using calibrated nucleic acid amplification tests (like reverse transcription quantitative PCR, RT-qPCR) is routinely performed to monitor response to treatment and ultimately prevent viral transmission. RNA quantities measured by commercial tests can vary over many orders of magnitude, from trace single copy levels to, in cases, over 10^9^ /mL of plasma, presenting an analytical challenge for calibrating across a broad measurement range. Interlaboratory study CCQM-P199 “HIV-1 RNA copy number quantification” (April to September 2019) was conducted under the auspices of the Consultative Committee for Amount of Substance (CCQM) Nucleic Acid analysis Working Group (NAWG), with the aims of supporting national metrology institutes (NMIs) and designated institutes (DIs) development of the capacity and evaluating candidate reference measurement procedures for applied viral nucleic acid measurements.

Thirteen laboratories participated in CCQM-P199 and were requested to report the RNA copy number concentration, expressed in copies per microliter, of the HIV-1 group specific antigen (*gag*) gene of *in vitro* transcribed RNA molecules at low (≈ 10^3^ /μL) and high concentration (≈ 10^9^ /μL) (Study Materials 1 and 2, linked by gravimetric dilution) and purified genomic RNA from cultured virus (Study Material 3). Study Materials 1 and 3 were measured by participants using one-step reverse transcription digital PCR (RT-dPCR) (Bio-Rad reagents) and/or two-step RT-dPCR with alternative cDNA synthesis reagents. Study Material 2 was measured by both RT-dPCR (one-step) (*n* = 4) and orthogonal methods: single molecule flow cytometric counting (*n* = 2), high performance liquid chromatograph (HPLC) (*n* = 1) and isotype dilution-mass spectrometry (ID-MS) (*n* = 1).

Interlaboratory reproducibilities (expressed as %CV) were 21.4 %, 15.3 % and 22.0 % for Study Materials 1, 2 and 3 respectively. Analysis of overdispersion showed that the interlaboratory variation for all three Study Materials was not accounted for in their reported uncertainties, indicating uncharacterized sources of variation remain. Although the mean values of RT-dPCR and orthogonal method results were not statistically significantly different (*p* = 0.46), the extrapolated mean Study Material 2 results were higher than mean Study Material 1 results (1196 *vs*. 808 /μL; *p* < 0.05). Follow-up analysis of Study Material 2 purity by ultra-performance liquid chromatography (UPLC) indicated higher molecular weight (MW) impurities constituted 16.6 % of the molecules, which are hypothesised to be the cause of the HPLC and ID-MS results being higher than the majority of Study Material 1 and 2 results.

This study demonstrates that reproducible measurement of RNA templates was achieved by metrology laboratories, illustrating the potential of RT-dPCR combined with complimentary orthogonal approaches to support traceability and precision of contemporary methods for RNA quantification. This study also highlighted that detailed characterization of RNA materials and sources of bias affecting measurements such as RT efficiency is needed to further establish RT-dPCR as a primary reference measurement procedure for RNA copy number quantification.

## INTRODUCTION

Plasma viral RNA load is routinely quantified for management of human immunodeficiency virus type 1 (HIV-1) infection. Quantitative estimates of viral load in copies per mL (of plasma) are used to monitor patient response to antiretroviral therapy and predict the progression of infection [1, 2]. Clinical assays target HIV-1 sequences including the long terminal repeat (LTR), polymerase (*pol*) and *gag* regions within the HIV-1 RNA viral genome (Figure 1) [3, 4]. The diagnostic range of tests for RNA copy number concentration of HIV-1 ranges from a limit of detection of about 20 /mL to >10^7^ /mL of sample [4] (approximately <2 /µL to 10^4^ /µL in the sample eluate, dependent on the extraction approach used), with sample matrices including human RNA background, which originates from blood cells and cell-free RNA in plasma.

**Figure 1:**
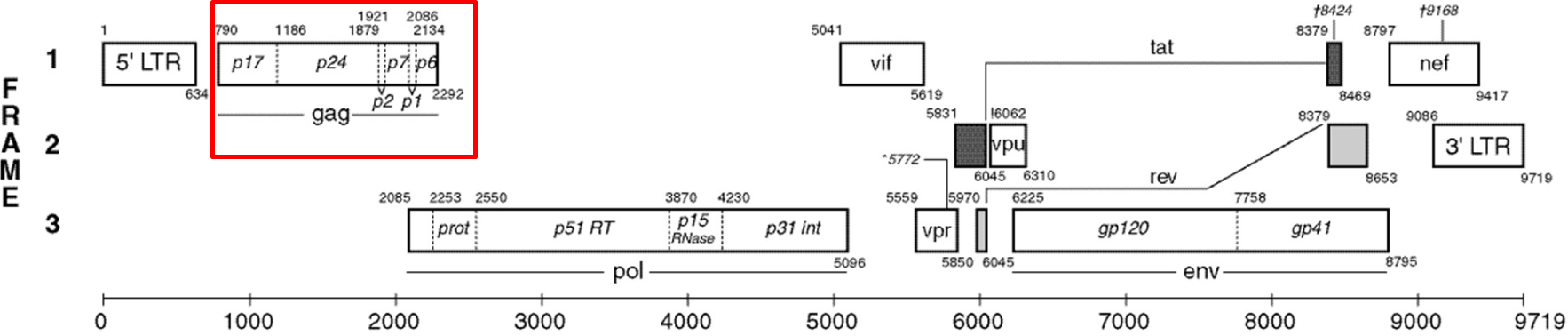
Schematic of the K03455.1 human immunodeficiency virus type 1 (HXB2) HIV1/HTLV-III/LAV reference genome [8]. The gag gene is highlighted by red box.

Clinical platforms used to quantify HIV-1 viral load often utilise secondary calibrators that are traceable to the HIV-1 RNA World Health Organization (WHO) International Standard (IS) with quantity values defined in International Units (IU). This is currently provided through use of the WHO 4^th^ IS (NIBSC code: 16/194), as an international conventional calibrator [5]. Assigned IU values are provided through consensus agreement following analysis by multiple laboratories using different approaches for nucleic acid amplification, including qPCR and transcription mediated amplification (TMA), and numerous target genes [6]. However, HIV-1 viral load continues to be reported clinically in copies per mL [7].

Establishing reference measurement procedures (RMPs) for value assignment of representative materials, such as RNA in buffered solution, in copy number concentration or IU could complement established material-based strategies for the standardization of measurements between laboratories. Reproducibility between quantitative results may facilitate continuity between different batches of IS used to value assign secondary reference materials and/or instrument calibrators [9]. Reverse transcription digital PCR (RT-dPCR) has been proposed as a candidate RMP for HIV-1 RNA copy number quantification [9].

CCQM-P199 sought to address the accuracy and interlaboratory comparability of RT-dPCR as a general candidate RMP methodology for quantification of targeted RNA copy number concentration in buffered solution, as well as to test specific candidate RMPs for HIV-1 RNA gene quantification. The *gag* gene was chosen as it is a common diagnostic target [10], alongside a concentration range relevant to that of purified clinical HIV-1 RNA extracts (≈ 10^1^/μL to ≈ 10^4^ /μL). This study differs from earlier NAWG RNA pilot studies as the participants were not provided with a recommended assay. Participants were provided with the sequence only and consequently the study also assesses participants’ ability to select appropriate assays. The diversity of assays further assessed the impact of different assays on RT-dPCR efficiency and specificity.

To test the trueness of RT-dPCR and to inform routes for SI-traceability (such as the requirement for calibration), RT-dPCR measurements were compared with “*orthogonal methods”:* high performance liquid chromatography (HPLC), isotype dilution mass spectrometry (ID-MS) and single molecule flow cytometry (FC) [11–13] which do not require reverse transcription or amplification, or enzymatic steps which may lead to biases in RT-dPCR [14]. To address the metrological state of the art and support analytical standardisation of diagnostic methods, *in vitro* synthesised and purified genomic RNA templates were chosen as the Study Materials for analysis. Therefore, the study aimed to provide evidence for capability of NMIs/DIs for value assignment of synthetic calibration materials and purified RNA materials. This study does not explore biological matrix-based materials requiring RNA extraction.

The following sections of this report document the timeline of CCQM-P199, the measurands, Study Materials, participants, results, discussion of factoring influencing comparability and trueness of measurements and consensus RVs. The Appendices (Supplementary Materials) reproduce the official communication materials and summarises information about the results provided by the participants.

## MEASURAND

The measurand (quantity intended to be measured [15]) of CCQM-P199 is RNA copy number concentration of the HIV-1 *gag* gene (Figure 1). The sample template type is *in vitro* transcribed RNA or purified viral genomic RNA, in a matrix of buffered solution (Study Material 2), or in a complex (human total RNA) background (Study Materials 1 and 3).

- **Measurand:** RNA copy number concentration expressed in copies per µL of the HIV-1 *gag* gene (K03455.1: 790 – 2257, KJ019215.1: 220 -1728)

Sequence information is provided in Appendix A (Supplementary Material).

## STUDY MATERIALS

### Background

Three Study Materials were prepared by NML (Study Materials 1 and 2) and jointly with NIBSC (Study Material 3). Coordinating laboratory methodology is provided in Appendix B (Supplemental material). All materials were synthetic or purified RNA, non-infectious requiring bio-safety level 1 containment. Sequence information is summarised in Table 1 and RNA sequence information for each material is provided in Appendix A (Supplemental material). It was expected that all study participants analyse Study Material 1 whereas analysis of Study Materials 2 and 3 was optional. Study participants were provided with four units of each Study Material. Details regarding the preparation, characterization and coordinators’ value assignment of each study material are included in the supplementary information.

**Table 1:**
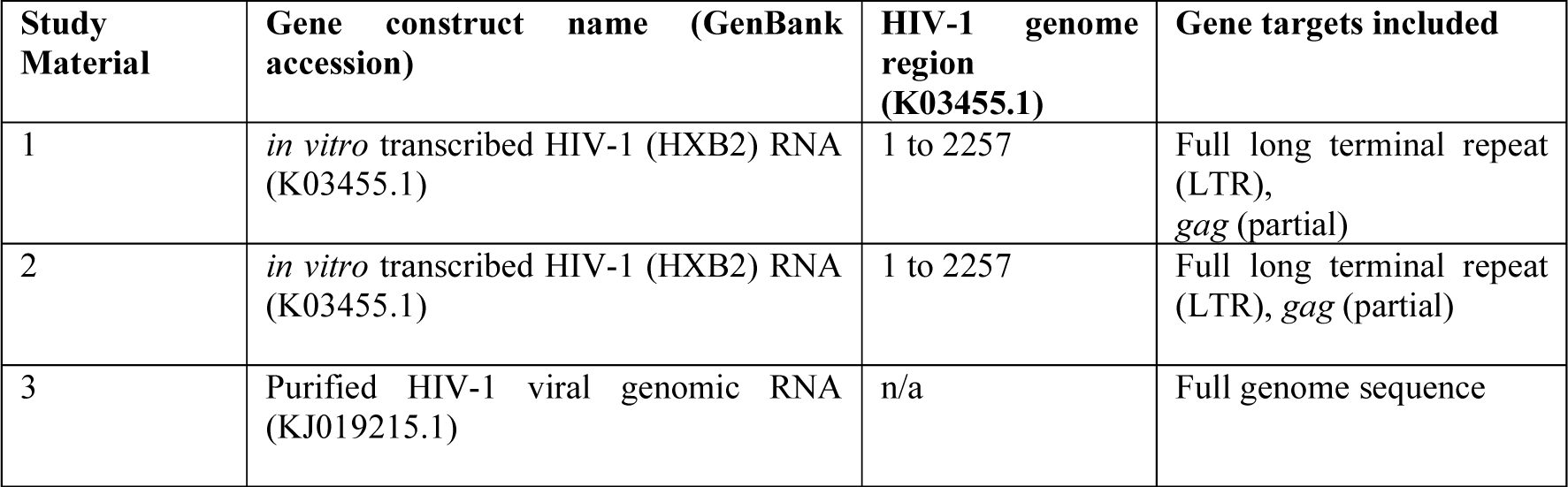
Summary of Study Material genomic information.

| Study Material | Gene construct name (GenBank accession) | HIV-1 genome region (K03455.1) | Gene targets included |
| --- | --- | --- | --- |
| 1 | *in vitro* transcribed HIV-1 (HXB2) RNA (K03455.1) | 1 to 2257 | Full long terminal repeat (LTR), *gag* (partial) |
| 2 | *in vitro* transcribed HIV-1 (HXB2) RNA (K03455.1) | 1 to 2257 | Full long terminal repeat (LTR), *gag* (partial) |
| 3 | Purified HIV-1 viral genomic RNA (KJ019215.1) | n/a | Full genome sequence |

### Preparation of Study Materials

**Study Material 1** was composed of an *in vitro* transcribed HIV-1 (HXB2) RNA molecule at an approximate copy number concentration ≈ 10^3^ /μL in ≈ 5 ng/μL human Jurkat cell line total RNA (Ambion “FirstChoice Human T-Cell Leukemia (Jurkat) Total RNA”, P/N AM7858) in buffered solution (1 mM^1^ sodium citrate, pH 6.5 (RNA Storage Solution Thermo Fisher Scientific P/N AM7001)). Initial measurement of a 20 -fold volumetric dilution of Human T-Cell Leukaemia RNA concentration was performed using a Qubit RNA BR kit, and the solution was subsequently gravimetrically diluted 10 -fold in RNA Storage Solution. Study Material 1 was prepared by gravimetric dilution of Study Material 2 using a Mettler Toledo XP205 balance to 5 decimal places. Following cleaning of the balance, linearity was tested using a set of laboratory standard weights covering the range 0.1 g to 200 g. Standard uncertainty of measurement for the balance was ± 0.000159 g (based on the calibration certificate). Further details can be found in Appendix B (Table B-5). A total of 327 units, each containing 100 µL, were prepared.

**Study Material 2** contained the same *in vitro* transcribed HIV-1 (HXB2) RNA molecule as Study Material 1 at an approximate copy number concentration of 10^9^ /µL to 10^10^ /μL (≈ 1 ng/μL to 20 ng/μL) in RNA Storage Solution (as above) in a volume of 200 μL per unit. A total of 120 units were prepared. No additional RNA molecules were added to this material as it was designed to be suitable for analysis using chemical methods and single molecule flow cytometry, where background nucleic acids will or may interfere with measurement.

**Study Material 3** was prepared from purified HIV-1 viral genomic RNA from viral stocks used to prepare the WHO 3^rd^ and 4^th^ IS for HIV-1 [16]. Prior to RNA purification, the HIV viral stock was heat inactivated for 1 h and then tested via tissue culture passage for one month with no viral growth detected. This is in ‘Appendix A to the Protocol’ (letter from NIBSC dated Jan 18, 2019) in Appendix C of this report (included as circulated to participants). RNA was purified from 4 vials using the QIAamp UltraSens Virus (Cat no 53704, Qiagen) and each was eluted in 60 µL of Buffer AVE (Qiagen). The four 60 µL eluates were pooled and subsequently diluted in ≈ 5 ng/μL human Jurkat cell line total RNA in RNA Storage Solution (as above) to an approximate RNA copy number concentration of ≈ 10^2^ /µL. Each unit of material contains 100 μL sample. A total of 325 units were prepared.

### Homogeneity Assessment of Study Materials

The homogeneity of all Study Materials was assessed by performing eight replicate measurements (sub-samplings) of 10 units. The detailed methods used by coordinator laboratories is described in Appendix B. The homogeneity of Study Materials 1 and 3 was evaluated by RT-dPCR analysis (Bio-Rad QX200). The homogeneity of Study Material 2 was evaluated by fluorimetric assay (QuantiFluor RNA System using the Quantus fluorimeter (Promega)).

### Analysis and results of homogeneity studies

Data was analysed using linear mixed effects models. The magnitude and statistical significance of between unit variation (standard deviation s_b_) are given in Table 3.

**Table 2:** Summary of gravimetric dilutions linking the production of Study Materials 1-2.

| Dilution | Mass RNA solution (g) | Mass RNA + Diluent (g) | Dilution factor (DF) | Relative $u$ (weighing RNA) | Relative $u$ (weighing RNA + Diluent) | Combined rel ($u$) | Combined $u$ (DF) |
| --- | --- | --- | --- | --- | --- | --- | --- |
| D1 | 0.0487 | 0.4997 | 10.2608 | 0.46 % | 0.045 % | 0.46 % |  |
| D2 | 0.0483 | 0.9906 | 20.5093 | 0.47 % | 0.023 % | 0.47 % |  |
| D3 | 0.0499 | 1.0015 | 20.0701 | 0.45 % | 0.022 % | 0.45 % |  |
| D4 | 0.0736 | 1.4991 | 20.3682 | 0.31 % | 0.015 % | 0.31 % |  |
| D5 | 0.5879 | 1.7427 | 2.9643 | 0.038 % | 0.013 % | 0.040 % |  |
| SM1 | 1.6343 | 32.941 | 20.1560 | 0.014 % | 0.00068 % | 0.014 % |  |
| Sum of variances | | | | | | $7.33 \times 10^{-5}$ | |
| Sum of covariances ($r = 1$) | | | | | | $2.31 \times 10^{-4}$ | |
| **Combined DF** | | | 5.14E+06 | | | 1.74 % | $8.96 \times 10^4$ |
| **Expanded uncertainty ($U$)** | | | | | | 3.49 % | $1.79 \times 10^5$ |

Relative uncertainty of gravimetric steps was calculated using a standard uncertainty of 0.00022486 g for each of the series of two weighing steps being performed (mass of tube and mass of liquid being added to the tube). Dilution D1 was prepared from Study Material 2. To account for the possibility that uncertainties in each dilution step may be correlated, the sum of pairwise covariances between the uncertainty for each step was calculated based on a maximum correlation coefficient ( $r$ ) of 1.0. A coverage factor ( $k$ ) = 2 was applied to the combined uncertainties for all dilution steps and expanded uncertainty rounded up to 2 significant digits. The combined dilution factor is rounded to the same order of magnitude as the expanded uncertainty.

**Table 3:** Study Material homogeneity results.

| Study Material | Relative $s_b$ (%)<sup>*</sup> | Significance ($p$) |
| --- | --- | --- |
| 1 | 2.1 % | < 0.001 |
| 2 | 0.36 % | *N.S.* |
| 3 | 1.5 % | *N.S.* |

Key: *N.S.*, not significant ( $p > 0.05$ ). <sup>\*</sup>Rounded up and shown to 2 significant digits.

### Stability Assessment of Study Materials

#### Design of short-term stability studies

A short-term stability study (STS) was performed by incubation of study materials on dry ice or at raised ambient temperature (27 °C) for 1 day and 7 days and compared to reference temperature (-80 °C) (*n* = 3 units per condition). Stability for Study Materials 1 and 3 was assessed by RT-dPCR using the HIV LTR-*gag* assay (*n* = 3) and for Study Material 2 by fluorometric assay (as Homogeneity study) and Agilent 2100 Bioanalyzer (fragment size).

#### Results of short-term stability studies

The effects of incubation time and temperature were evaluated using mixed effect models and no significant effect of incubation time was evident. Therefore, the effect of the two shipment temperatures was compared with the reference temperature using a random effects model with temperature as a fixed effect and sample (unit) as a random effect. The results of this analysis are shown in Table 4. At raised ambient temperature, a decrease in the measured concentration of Study Materials 1 and 3 was observed. No impact of incubation at 27 °C was observed on the concentration of Study Material 2. No significant differences were observed compared to the reference temperature on any of the Study Materials.

**Table 4:**
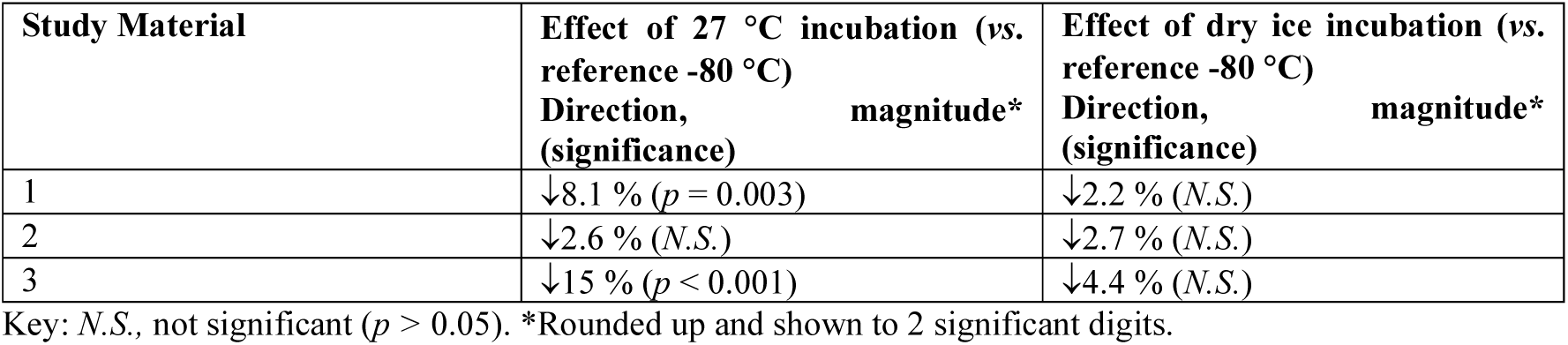
Results of Short-Term Stability studies.

| Study Material | Effect of 27 °C incubation (vs. reference -80 °C)<br>Direction, magnitude*<br>(significance) | Effect of dry ice incubation (vs. reference -80 °C)<br>Direction, magnitude*<br>(significance) |
| --- | --- | --- |
| 1 | ↓8.1 % ($p = 0.003$) | ↓2.2 % (*N.S.*) |
| 2 | ↓2.6 % (*N.S.*) | ↓2.7 % (*N.S.*) |
| 3 | ↓15 % ($p < 0.001$) | ↓4.4 % (*N.S.*) |

Key: *N.S.*, not significant ( $p > 0.05$ ). \*Rounded up and shown to 2 significant digits.

Analysis of Study Material 2 using the Agilent Bioanalyzer (Figure 2) showed a peak corresponding to the expected size of 2,266 bp which was observed under all simulated shipment conditions, with no degradation products visible. Therefore, it was concluded that dry ice shipment was suitable for all Study Materials but extended periods at room temperature should be avoided.

**Figure 2:**
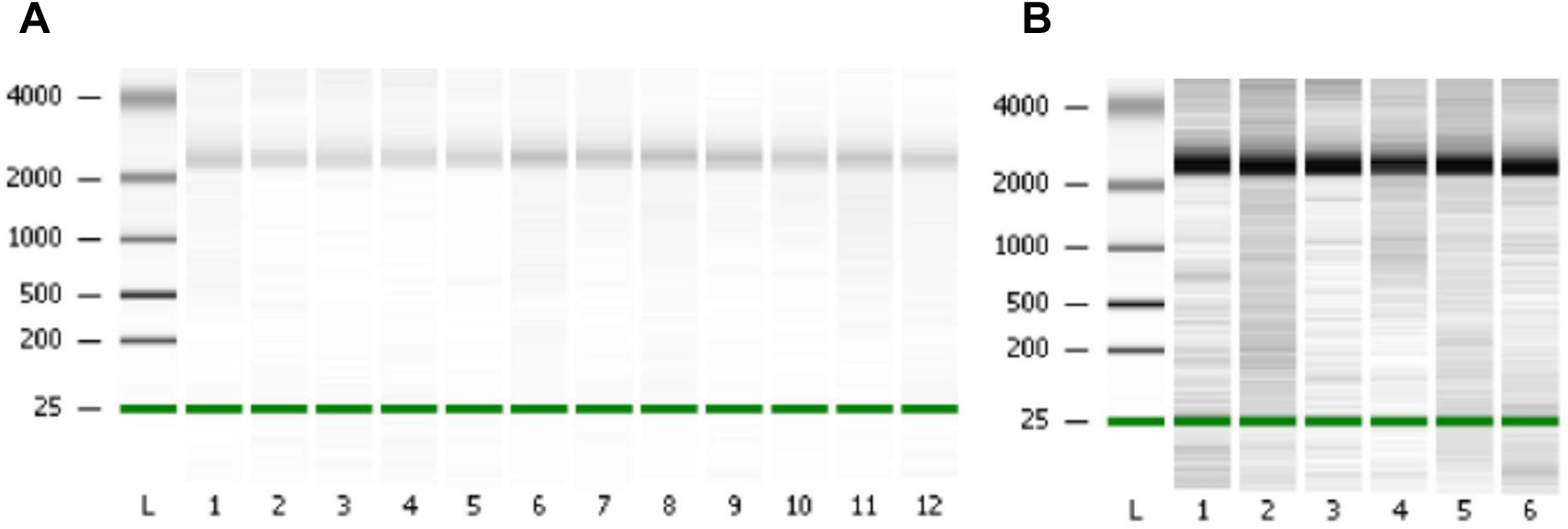
Study Material 2 short term stability: Fragment size analysis. Study Material 2 was analysed with the RNA 6000 Nano kit for the 2100 Bioanalyzer (Agilent). (A) Lane: L, RNA ladder; 1 to 3, Dry ice 1 day; 4 to 6, Dry ice 7 days; 7 to 9, 27 °C 1 day; 10 to 12, 27 °C 7 days. (B) Lane: L, RNA ladder; 1 to 3, -80 °C 1 day; 4 to 6, -80 °C 7 days.

#### Design of long-term stability studies

Long-term stability of the Study Materials was assessed at two time points; 4 months to 5 months and 7 months to 9 months after the homogeneity and short-term stability studies (Figure 3), and changes in copy number concentration were evaluated by RT-dPCR using the HIV-1 *gag* assay. Prior to LTS analysis, four units of each of Study Materials 1, 2 and 3 were incubated on dry ice for 7 days to simulate conditions during shipping. These were compared to four units of each material that had been stored at the reference temperature of -80 °C without dry ice incubation. A volumetric dilution series was prepared for each of the eight units (four with and four without dry ice incubation) of Study Material 2 to a concentration of 5,000 /µL based on RT-dPCR analysis. Study Material 2 was also measured by fluorimetry.

**Figure 3:**
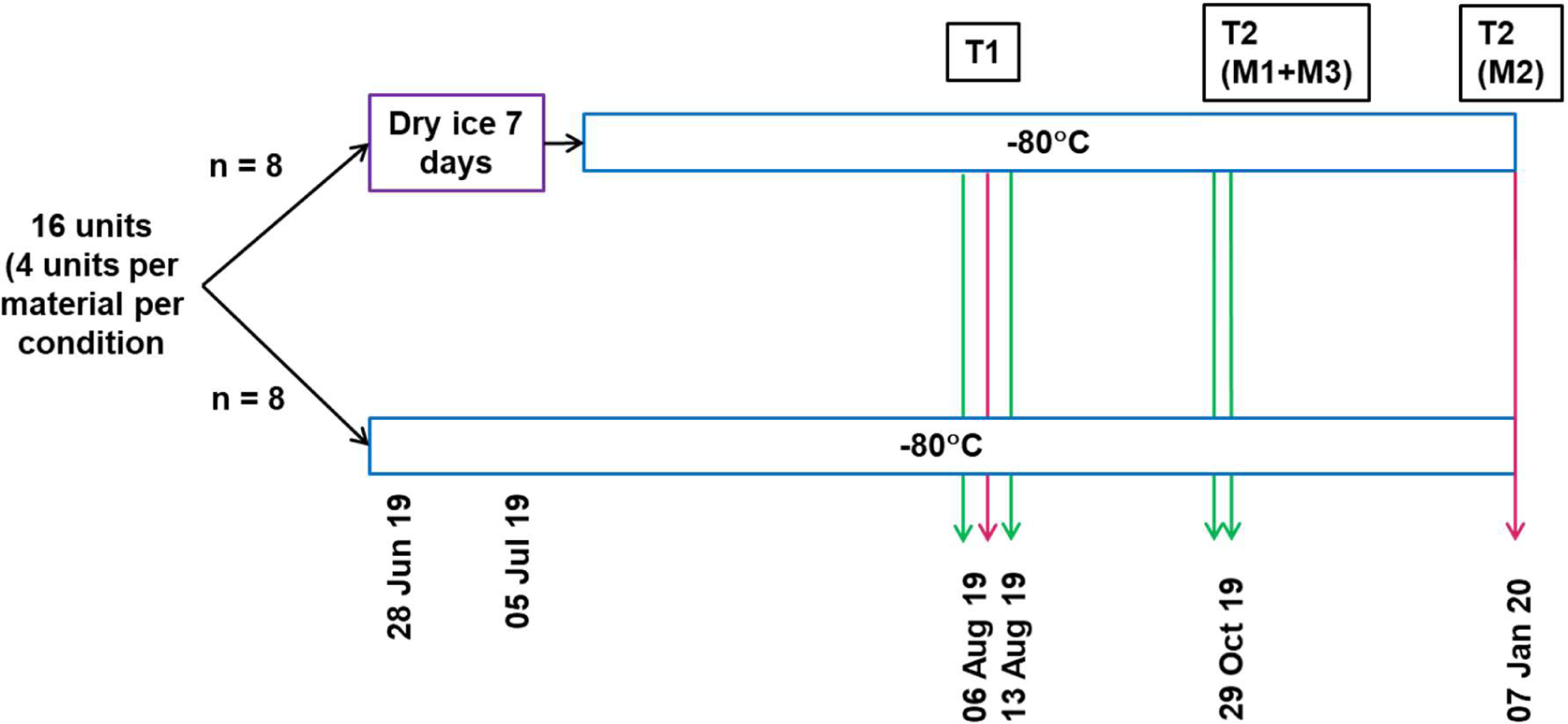
Design of Long-term stability study.

#### Results of long-term stability studies

The results of long-term stability studies are shown in Figure 4. No significant effect of storage time or temperature was observed for Study Materials 1 or 3, however RT-dPCR analysis indicated a significant decrease in concentration (-8.4 % relative to the first timepoint) was observed for Study Material 2 over time (*p* < 10^-4^, Figure 4B). This could indicate instability of the material however technical factors such as inaccuracy in the volumetric dilution series or variation in performance of RT-dPCR (due to reagent batch or ambient conditions) may have caused the observed differences. Fluorimetric measurements of Study Material 2 were similar between the STS and LTS (Figure 4D) and may also have been affected by variability in the calibration and performance of this approach. Moreover, the fluorimetric measurements are not expected to be significantly affected by subtle changes in RNA stability associated with fragmentation. Therefore it is unclear whether the results of LTS for Study Material 2 reflect an accurate estimation of material stability.

**Figure 4:**
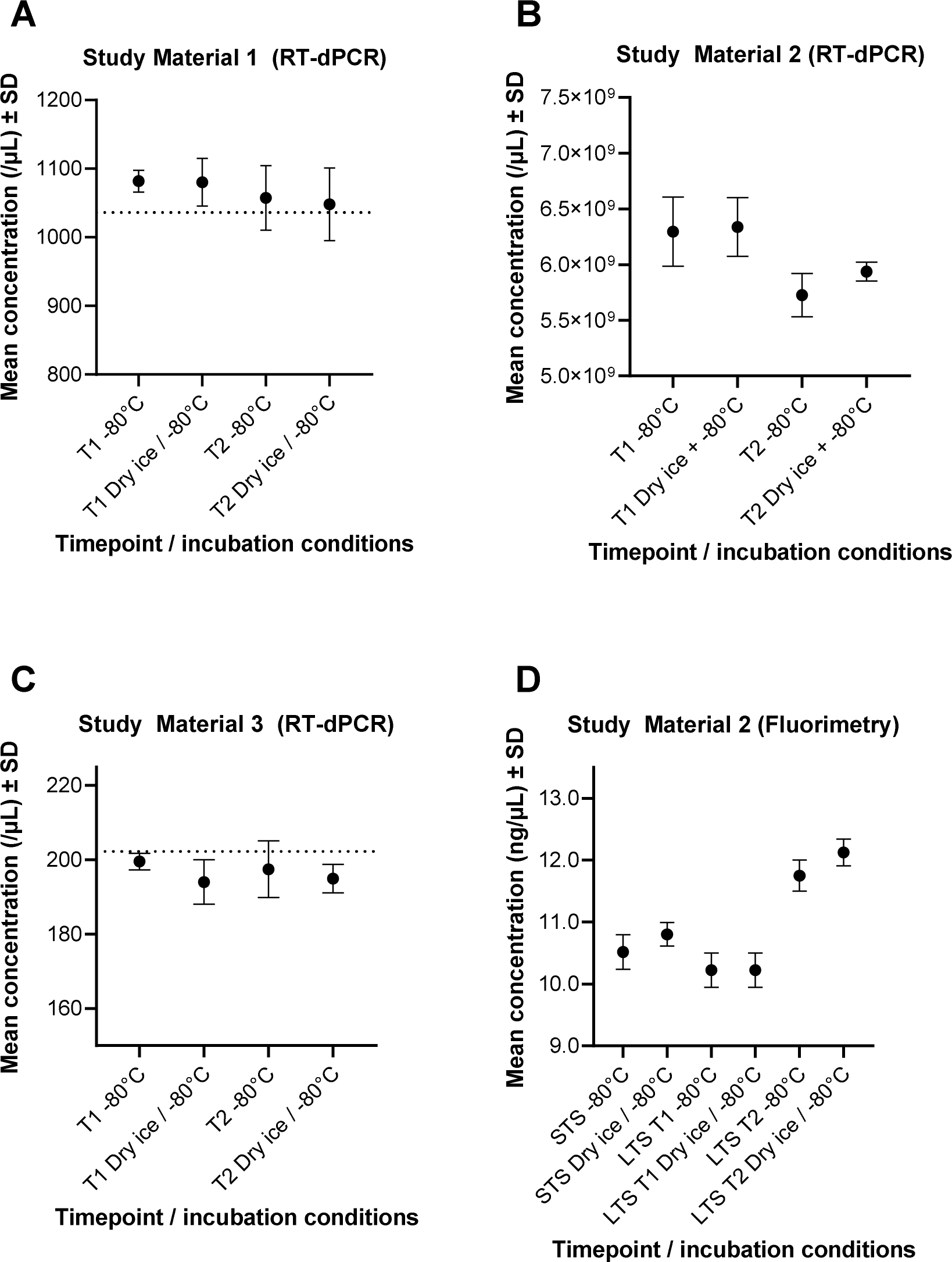
Results of the long-term stability assessment studies. (A) Study Material 1 (B) Study Material 2 and (C) Study Material 3 LTS results based on RT-dPCR and (D) Study Material 2 STS and LTS results based on fluorimetric measurements. Dots represent mean value per unit, error bars represent SD. Study Materials 1 and 3 are plotted compared to the homogeneity data (dashed line) which was assessed by RT-dPCR. (Study Material 2 homogeneity was not measured by RT-dPCR). Four units were analysed for each Study Material.

### Coordinators’ value assignment of Study Materials

The coordinator’s assigned values for Study Materials 1, 2 and 3 are shown in Table 5 based on measurements performed with the HIV-1 *gag* assay. Contributions to the uncertainty in the assigned values are shown in Table 6.

**Table 5:** Coordinator’s assigned values and uncertainties.

| **MATERIAL** | **MEASURAND (UNIT)** |
| --- | --- |
| **Study Material 1** | **Measurand 1 (*gag* gene)** |
| Value (*x*) | 1040 /μL (1035.6 /μL as calculated) |
| Standard uncertainty (*u*) | 86.4 /μL |
| Coverage factor (*k*) | 2.447 |
| Relative expanded uncertainty (Rel *U*) | 20.4 % (as calculated) |
| Expanded uncertainty (*U*) | 220 /μL |
| **Study Material 2** | **Mass concentration** |
| Value (*x*) | 10.93 ng/μL |
| Equivalent copy number concentration (copies/μL) | $9.0 \times 10^9$ /μL |
| Standard uncertainty (*u*) | 0.51 ng/μL |
| Coverage factor (*k*) | 4.3 |
| Relative expanded uncertainty (Rel *U*) | 20.1 % |
| Expanded uncertainty (*U*) | 2.2 ng/μL |
| **Study Material 3** | **Measurand 1 (*gag* gene)** |
| Value (*x*) | 203 /μL (202.58 /μL as calculated) |
| Standard uncertainty (*u*) | 17.4 /μL |
| Coverage factor (*k*) | 2.447 |
| Relative expanded uncertainty (Rel *U*) | 21.0 % (as calculated) |
| Expanded uncertainty (*U*) | 43 /μL |

**Table 6:** Uncertainty contributions to coordinator’s assigned values.

| **Factor** | **Study Material 1** | **Study Material 2** | **Study Material 3** |
| --- | --- | --- | --- |
| Method precision (%) | 2.45 % | 4.67 % | 3.27 % |
| Homogeneity (%) | 2.03 % | 0.36 % | 1.89 % |
| RT-dPCR assay* | 5.70 % | Not applicable | 5.70 % |
| Partition volume (%) | 5.19 % | Not applicable | 5.19 % |
| Stability (%) | No allowance | No allowance | No allowance |

Contributions are shown as relative standard uncertainties (RT-dPCR for SM1 and SM3, Qubit fluorimetry for SM2). \*Variation between alternative primers/probes (Type B).

### SAMPLE DISTRIBUTION

All Study Materials were shipped on dry ice. Dates below reflect the shipment dates for units that were analysed and does not include dates relating to failed shipments (see comments for more information).

### TIMELINE

Table 8 lists the timeline for CCQM P199.

**Table 7:**
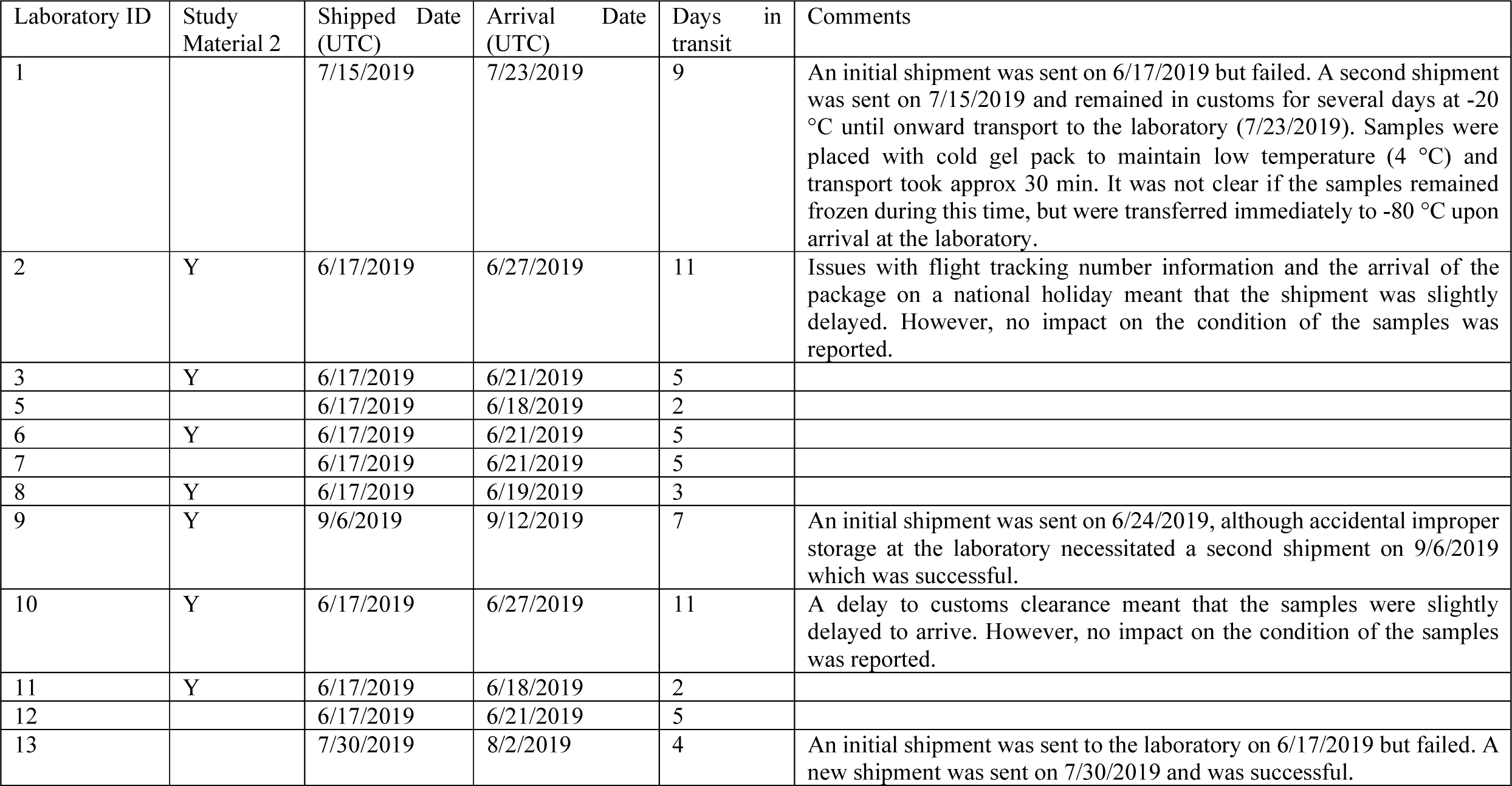
Distribution of Study Materials for CCQM-P199.

**Table 8:** Timeline for CCQM P199.

| Date | Timeline | Comments |
| --- | --- | --- |
| CCQM NAWG Apr 2018 | Viral (HIV-1) RNA quantification Pilot Study proposed to the NAWG | Aligned to EMPIR 15HLT07 AntiMicroResist project |
| June 2018 | Pilot study number assigned |  |
| CCQM NAWG Oct 2018 | Update to proposed P199 study given | Presentation of revised approach to production of *in vitro* transcribed (IVT) materials (Study Material 1, 2) with alternative plasmid restriction digestion towards 3' end of *gag* gene |
| January 14, 2019 | "CCQM P199 HIV-1 target sequences v1.0_14 Jan 2019" document circulated | Email from coordinator Jan. 15, 2019 |
| April 2019 CCQM | Call for participation |  |
| May 17, 2019 | Final protocol circulated |  |
| May 31, 2019 | Registration for participation | Participants reply with Form 1 |
| June-July 2019 | Study Material distribution | Participants reply with Form 2 |
| September 23, 2019 | Submission of results | Participants reporting with Forms 3, 4, 5 |
| October 03 to 04, 2019 | Initial report at NAWG (Turin) | Presentation of Study Material 1 and 3 results |
| May 12, 2020 | Results discussion NAWG (Webex) | Presentation of Study Material 2 results<br>Follow-up to Study Material 1 and 3 results (Laboratories 6 and 12) |
| November 18, 2020 | Results discussion NAWG (Webex) | Study Material 2 purity analysis (Laboratory 10)<br>Minor revision to Laboratory 9 Study Material 2 result based on amended MW |
| May 11, 2021 | Coordinator update NAWG (Webex) | Study Material 1 and 2 results comparison<br>Candidate consensus reference values presented |
| July 16, 2021 | Draft A report distributed |  |
| March 2022 | Draft B report distributed |  |
| November 2022 | Final report circulated to NAWG |  |

## RESULTS

Participants were requested to report the RNA copy number concentration of the *gag* gene in Study Material 1. Measurement of Study Materials 2 and 3 were optional.

In addition to the quantitative results, participants were instructed to describe their experimental design and analytical methods using Study Reply Forms 4 and 5 (dMIQE) (Appendices F and G (Supplementary information)). Participants reported a summary of their methods, results and approach to measurement uncertainty in presentations given at CCQM meetings October 2019 (Study Materials 1 and 3) and May 2020 (Study Material 2).

CCQM-P199 results were reported by all 13 institutions that received samples.

### Methods Used by Participants

Study Materials 1 and 3 were analysed by participants using RT-dPCR (Table H-1, Supplementary information). Assays targeting different regions of the *gag* gene were used (Figure 5; Table H-3 (Supplementary information)).

**Figure 5:**
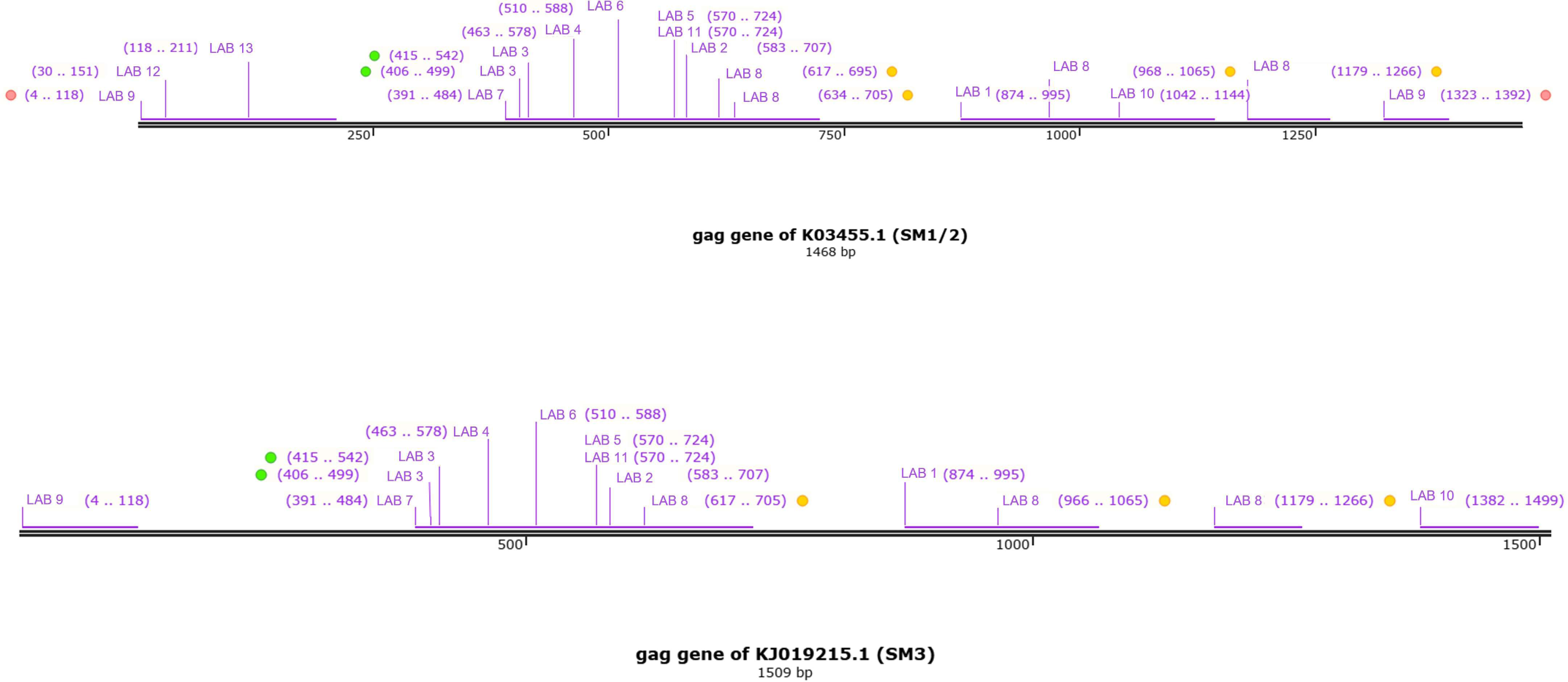
Alignment of CCQM-P199 participants’ RT-dPCR assays to *gag* gene sequence for Study Material 1/2 and Study Material 3. Institution number and assay designation shown with position in *gag* gene shown in brackets. Assays are colour coded in cases where institutes analysed more than one target per Study Material.

The majority of results were based on one-step RT-dPCR measurements using the QX100/200 system (Bio-Rad) (13/18 and 13/15 for Study Materials 1 and 3 respectively). Three laboratories reported results for a two-step RT-dPCR approach combined with the QX100/200 platform (Laboratory 12 and Laboratory 13 (Study Material 1 only) and Laboratory 3 (both Study Materials 1 and 3)). Laboratory 10 and Laboratory 12 also submitted results using a two-step approach with the Quantstudio 3D (QS3D) (Thermo Fisher Scientific) dPCR system. For one-step RT-dPCR with the QX100/200 platform, the reagent (One-step RT-ddPCR Advanced kit for Probes, Bio-Rad) is restricted by compatibility with the dPCR system. For two-step RT-dPCR, NMIs used alternative enzymes/kits for RT: Superscript III (Thermo Fisher Scientific) was used by Laboratory 3, Superscript IV (Thermo Fisher Scientific) was used by Laboratory 10 and RevertAid H Minus First Strand cDNA Synthesis Kit (Thermo Fisher Scientific) was used by Laboratory 13. Following submission and presentation of the study data, Laboratory 12 identified an error in the process implemented for two-step RT-dPCR consisting of an RT-PCR reagent being used for the RT step. This affected results for both dPCR platforms (QX200 and QS3D).

Three laboratories applied correction for RT efficiency to results: Laboratory 1, Laboratory 2 and Laboratory 6 (Table H-6 (Supplementary information)). Laboratory 1 used a 74 nucleotide RNA oligo as an internal amplification control for RT efficiency correction. Laboratory 2 (main results) and Laboratory 6 (supplementary results) applied RT efficiency correction based on UV spectrometric measurements of Study Material 2.

Study Material 2 was analysed using a range of techniques (Table H-2 (Supplementary information)): RT-dPCR (4 results), single molecule flow cytometric counting (2 results), HPLC with UV detection (1 result) and isotope dilution mass spectrometry (ID-MS) (1 result). The HPLC measurements were calibrated to an NMIJ RNA CRM panel (Table 10).

**Table 9:**
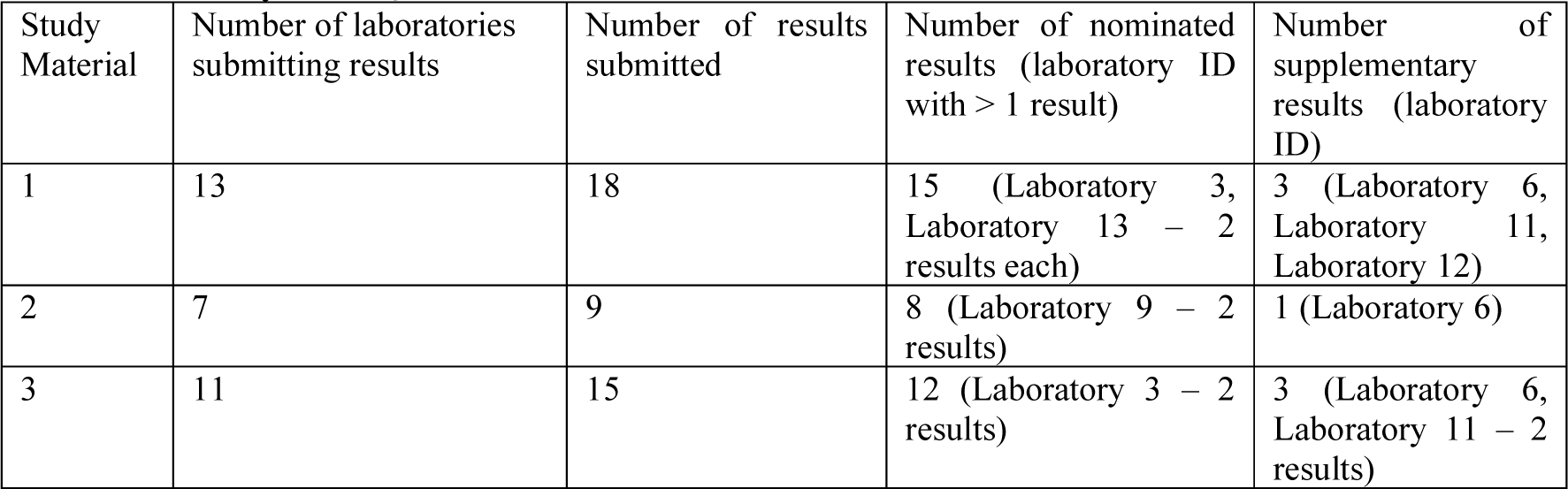
Summary of CCQM-P199 submitted results.

| Study Material | Number of laboratories submitting results | Number of results submitted | Number of nominated results (laboratory ID with > 1 result) | Number of supplementary results (laboratory ID) |
| --- | --- | --- | --- | --- |
| 1 | 13 | 18 | 15 (Laboratory 3, Laboratory 13 – 2 results each) | 3 (Laboratory 6, Laboratory 11, Laboratory 12) |
| 2 | 7 | 9 | 8 (Laboratory 9 – 2 results) | 1 (Laboratory 6) |
| 3 | 11 | 15 | 12 (Laboratory 3 – 2 results) | 3 (Laboratory 6, Laboratory 11 – 2 results) |

**Table 10:** Certified Reference Materials Used.

| Laboratory ID | CRM (batch no.) | Provider | Analyte | Certified value and uncertainty<sup>a</sup> | Method used to value assign CRM [CCQM study or CMC demonstrating competency] | Fragment size/Purity evaluation |
| --- | --- | --- | --- | --- | --- | --- |
| 9 | CRM 6204-b RNA standards (1000A, 1000B) (Batch 080) | NMIJ | 1000 nucleotide sequence specified in certificate | RNA1000A:<br>$x = 68.2 \text{ ng}/\mu\text{L}$<br>$U = 5.8 \text{ ng}/\mu\text{L}$<br>RNA1000B:<br>$x = 64.1 \text{ ng}/\mu\text{L}$<br>$U = 5.5 \text{ ng}/\mu\text{L}$ | ID-MS (ribonucleotides)<br>ICP-MS (phosphorus) | Microchip gel electrophoresis (NMIJ) |
| 10 | In-house NMP standard solution | Lab. 10 | Concentration of nucleotide monophosphate | AMP: 1083.5 nmol/g<br>$\pm 31.4 \text{ nmol/g}$<br>CMP: 1179.2 nmol/g<br>$\pm 31.9 \text{ nmol/g}$<br>GMP: 1052.8 nmol/g<br>$\pm 24.2 \text{ nmol/g}$<br>UMP: 1155.7 nmol/g<br>$\pm 25.7 \text{ nmol/g}$ | <sup>1</sup>H NMR using potassium hydrogen phthalate CRM (NMIJ CRM 3001-b) as internal standard [appropriate CCQM study of qNMR was K55.d.] | N/A |

<sup>a</sup> Stated as Value ± expanded uncertainty ( $x \pm U_{95}(x)$ )

Following presentation and discussion of the Study Material 3 results, it was identified that Laboratory 6 had not received the sequence information for Study Material 3 (see Timeline). It was subsequently established (May 2020 NAWG) that the reverse primer of the assay used by Laboratory 6 (designated Assay 1) has partial homology to a region 3’ in the Study Material 3 template sequence (Figure 6).

**Figure 6:**
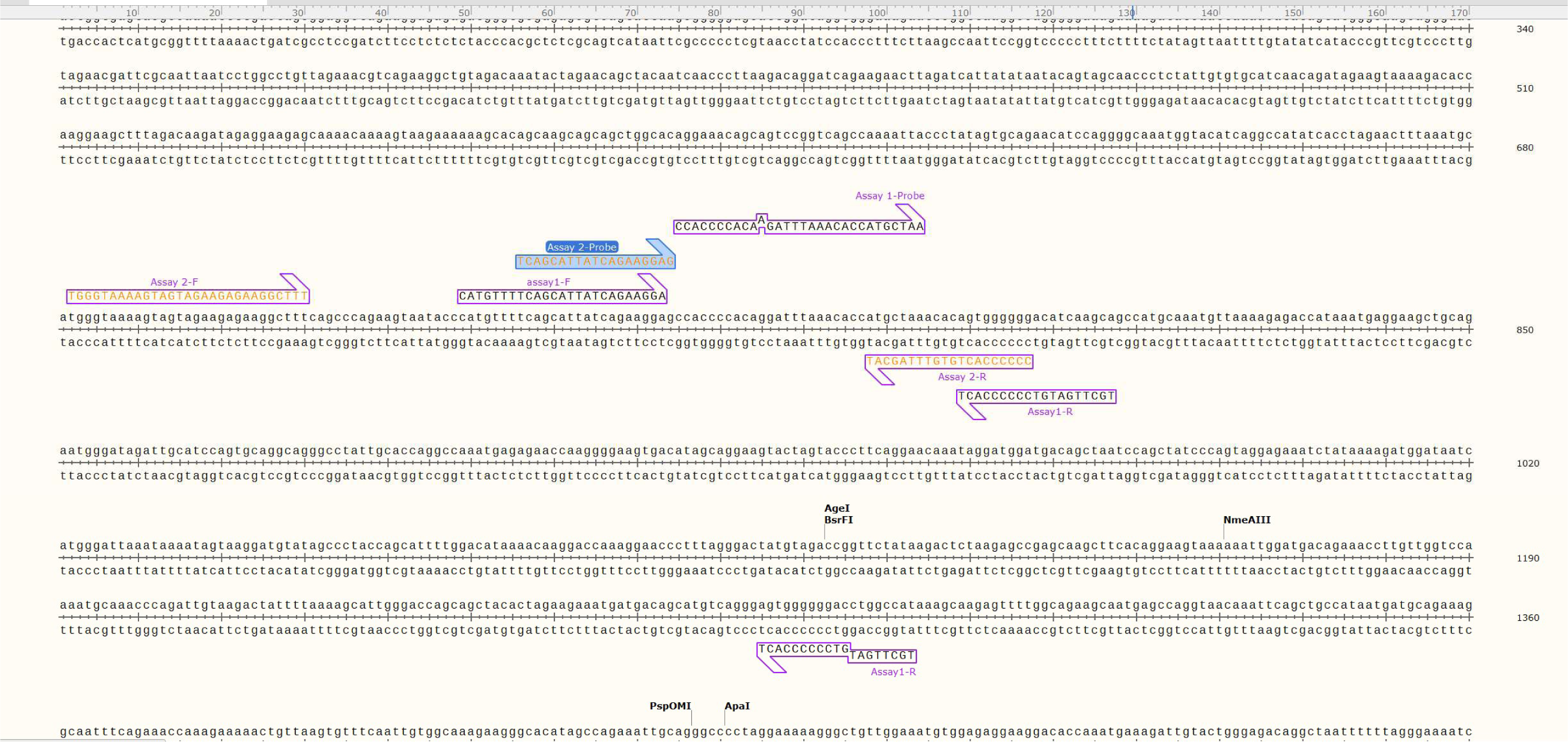
Alignment of Laboratory 6 Assays to Study Material 3. Two assays (Assay 1 and 2) were designed by laboratory 6to the *gag* sequence in Study Material 1/2 (Appendix A) and Assay 1 used for measurement of all three study materials (Table H-3 (Supplementary information)). Subsequent alignment of the assays to the *gag* sequence of Study Material 3 revealed potential binding of the Assay 1 reverse primer (Assay 1-R) to two positions.

**Figure 7:**
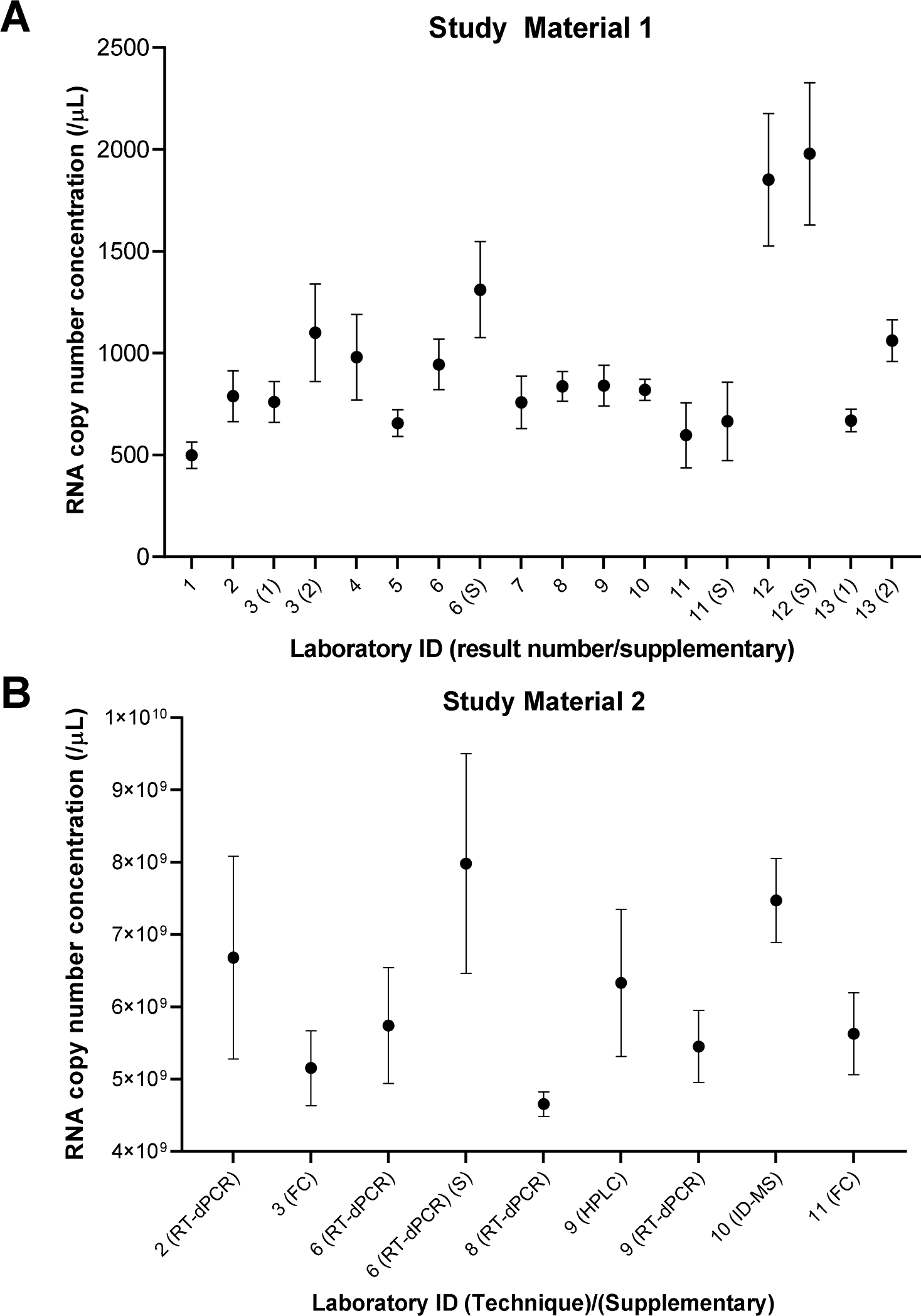
P199 participants reported results (A) Study Material 1 and (B) Study Material 2. Dots represent the reported values, *x*; bars their 95 % expanded uncertainties (99 % for Laboratory 8), *U*(*x*). For >1 nominated result, result number shown in parenthesis. Supplementary result indicated by (S).

**Figure 8:**
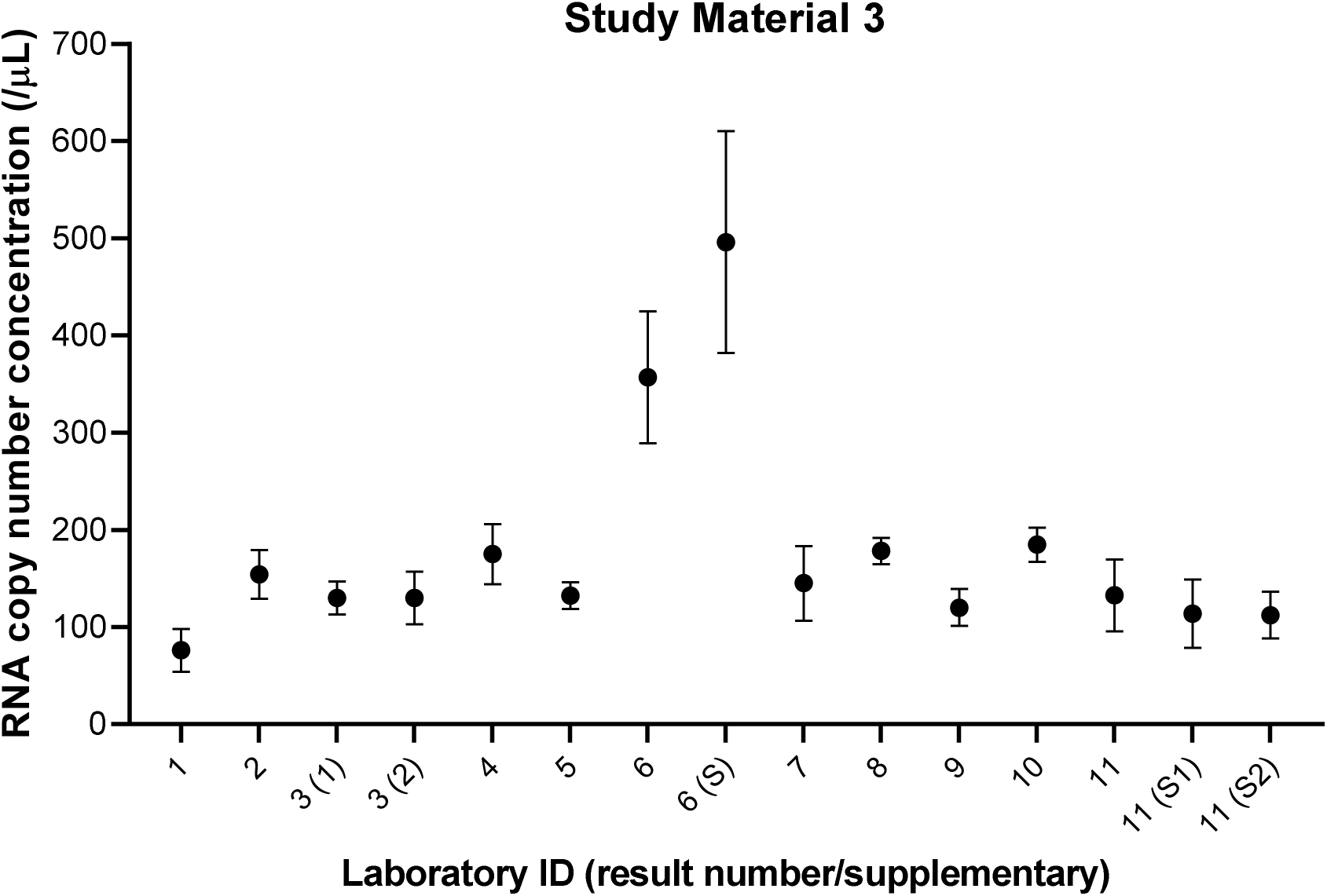
P199 participants reported results: Study Material 3. Dots represent the reported values, *x*; bars their 95 % expanded uncertainties, *U*(*x*). For >1 nominated result, result number shown in parenthesis. Supplementary result indicated by (S) and number if >1 supplementary result.

Further details of the analytical methods used by participants are provided in Appendix H (Supplementary Information). The participants’ approaches to estimating uncertainty are provided in Appendix I (Supplementary Information).

### Calibration Materials Used by Participants (Study Material 2)

For HPLC (Laboratory 9) and ID-MS (Laboatory 10) measurements of Study Material 2, participants established the metrological traceability of their results using certified reference materials (CRMs) with stated traceability and/or commercially available high purity materials for which they determined the purity. Table 10 lists the CRMs that were used and how participants established traceability. If in-house value-assigned calibration materials were used, Table 10 lists the material, its assigned purity, the method used, and how the participant had demonstrated their competence in the use of the method(s).

### Participants’ Results

Tables 11 to 13 show the final participants’ reported results. In the case of an amendment to data following the first version of the dataset, a footnote to the table describes the amendment. The original results are shown in Appendix J (Supplementary information) where applicable.

**Table 11:** CCQM-P199 participants’ measurement results for Study Material 1.

| Lab ID | | Result number | Nominated* | Methodology** | $x$ (/μL) | $u$ (/μL) | $k$ | $U$ (/μL) | Rel $U$ (%) |
| --- | --- | --- | --- | --- | --- | --- | --- | --- | --- |
| 1 |  | 1 | Y | one-step RT-dPCR | 498 | 29 | 2.26 | 65 | 13 % |
| 2 |  | 1 | Y | one-step RT-dPCR | 788 | 62 | 2 | 124 | 15.8 % |
| 3 |  | 1 | Y | one-step RT-dPCR | 760 | 52 | 2.00 | 100 | 14 % |
| 3 |  | 2 | Y | two-step RT-dPCR | 1100 | 120 | 2.10 | 240 | 21 % |
| 4 |  | 1 | Y | one-step RT-dPCR | 980 | 79.3 | 2.57 | 210 | 21 % |
| 5 |  | 1 | Y | one-step RT-dPCR | 656.04 | 32.44 | 2 | 64.89 | 9.89 % |
| 6 |  | 1 | Y | one-step RT-dPCR | 944 | 62 | 2 | 124 | 13 % |
| 6 |  | 2 | S | one-step RT-dPCR (RT efficiency corrected) | 1311 | 118 | 2 | 236 | 18 % |
| 7 |  | 1 | Y | one-step RT-dPCR | 757.33 | 64.50 | 2 | 129.00 | 17 % |
| 8 |  | 1 | Y | one-step RT-dPCR | 837 | 28 | 2.58† | 73 | 8.8 % |
| 9 |  | 1 | Y | one-step RT-dPCR | 840 | 46 | 2.2 | 100 | 12 % |
| 10 |  | 1 | Y | two-step RT-dPCR | 818.9 | 26.1 | 2 | 52.2 | 6.37 % |
| 11 |  | 1 | Y | one-step RT-dPCR (Assay Ref. [17]) | 596.3 | 79.3 | 2.01 | 159.4 | 26.7 % |
| 11 |  | 2 | S | one-step RT-dPCR (Assay Ref. [18]) | 664.7 | 88.5 | 2.18 | 192.7 | 29 % |
| 12 |  | 1 | Y | two-step RT-dPCR (QX200) | 1851 | 163 | 2 | 325 | 17.6 % |
| 12 |  | 2 | S | two-step RT-dPCR (QS3D) | 1978 | 174 | 2 | 349 | 17.6 % |
| 13 |  | 1 | Y | one-step RT-dPCR | 669 | 28 | 2 | 56 | 8.4 % |
| 13 |  | 2 | Y | two-step RT-dPCR | 1061 | 51 | 2 | 102 | 9.6 % |

**Key:** $x$ , RNA copy number concentration (*gag* gene); $u$ , standard uncertainty; $k$ , coverage factor; $U$ , expanded uncertainty; *Rel U*, relative expanded uncertainty. \*Nominated results indicated by Y (Yes) or supplementary (S). \*\*For laboratories submitting >1 result, discriminating methodological information is provided in brackets. † Indicates a 99 % confidence interval (CI). *Note 1:* Minor amendments to reported standard uncertainty (Laboratory 5) or both combined and expanded uncertainties (Laboratory 12 nominated result). Original submitted values presented in Table J-1.

**Table 12:** CCQM-P199 participants’ measurement results for Study Material 2.

| Lab ID | | Result | Nominated* | Methodology | $x$ (/μL) | $u$ (/μL) | $k$ | $U$ (/μL) | Rel U (%) |
| --- | --- | --- | --- | --- | --- | --- | --- | --- | --- |
| 2 |  | 1 | Y | RT-dPCR | 6.68E+09 | 6.99E+08 | 2 | 1.40E+09 | 20.9 % |
| 3 |  | 1 | Y | Flow cytometry | 5.15E+09 | 1.1E+08 | 2.3 | 5.2E+08 | 10 % |
| 6 |  | 1 | Y | RT-dPCR | 5.74E+09 | 4.00E+08 | 2 | 8.00E+08 | 14 % |
| 6 |  | 2 | S | RT-dPCR (RT efficiency corrected) | 7.98E+09 | 7.60E+08 | 2 | 1.52E+09 | 19 % |
| 8 |  | 1 | Y | RT-dPCR | 4.652E+09 | 6.60E+07 | 2.57 | 1.700E+08 | 3.6 % |
| 9 |  | 1 | Y | HPLC | 6.33E+09 | 5.10E+08 | 2 | 1.02E+09 | 16 % |
| 9 |  | 2 | Y | RT-dPCR | 5.45E+09 | 2.1E+08 | 2.31 | 5.0E+08 | 9.1 % |
| 10 |  | 1 | Y | LC-MS/MS (ID-MS) | 7.47E+09 | 2.90E+08 | 2 | 5.80E+08 | 7.76 % |
| 11 |  | 1 | Y | Flow cytometry | 5.626E+09 | 1.78E+08 | 3.18 | 5.66E+08 | 10 % |

**Key:** $x$ , RNA copy number concentration (*gag* gene); $u$ , standard uncertainty; $k$ , coverage factor; $U$ , expanded uncertainty; *Rel U*, relative expanded uncertainty. \*Nominated results indicated by Y (Yes) or supplementary (S). *Note 1:* Laboratory 2 changed the number of significant figures reported in their standard uncertainty. *Note 2:* Laboratory 9 slightly amended their submitted value and uncertainty following application of an amended MW in their calculations. Original values shown in Appendix J Table J-2 (Supplementary information).

**Table 13:**
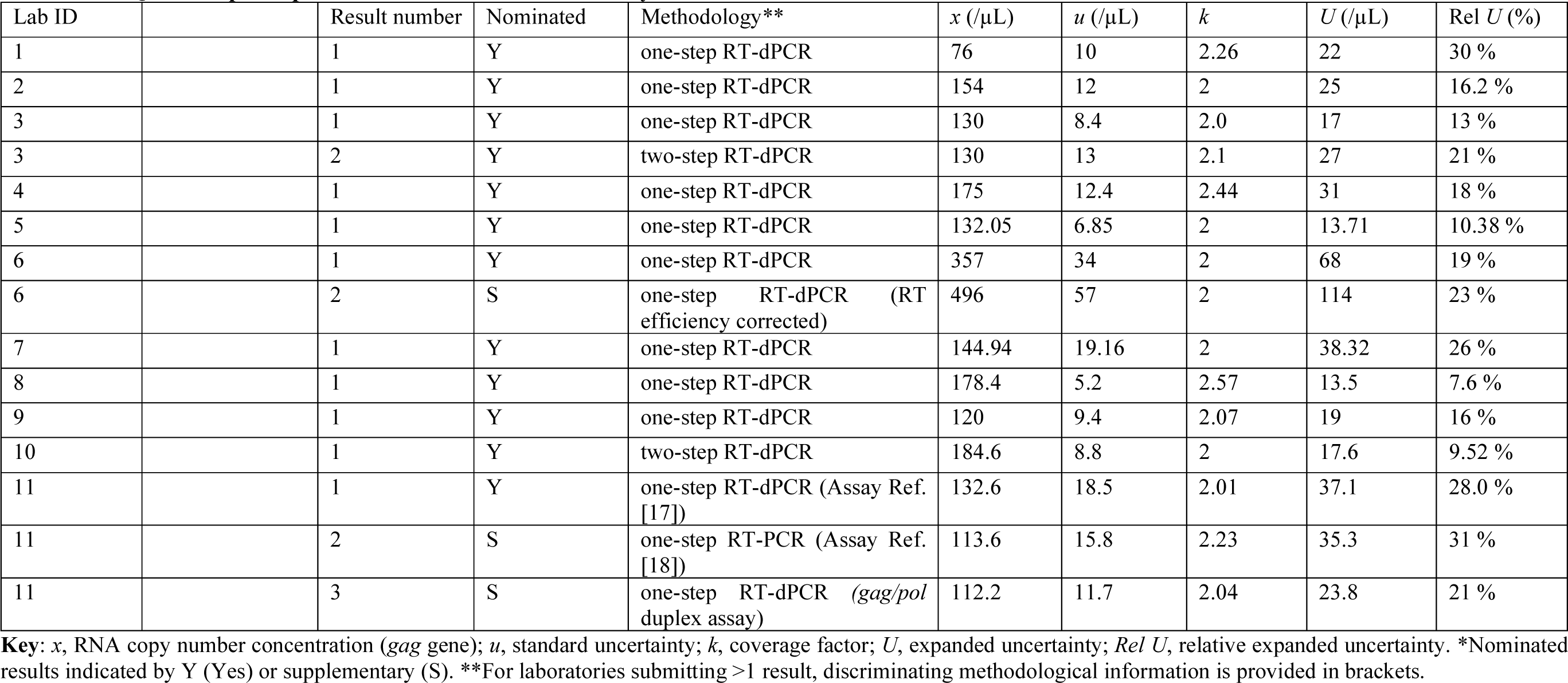
CCQM-P199 participants’ measurement results for Study Material 3.

| Lab ID | | Result number | Nominated | Methodology** | $x$ (/μL) | $u$ (/μL) | $k$ | $U$ (/μL) | Rel $U$ (%) |
| --- | --- | --- | --- | --- | --- | --- | --- | --- | --- |
| 1 |  | 1 | Y | one-step RT-dPCR | 76 | 10 | 2.26 | 22 | 30 % |
| 2 |  | 1 | Y | one-step RT-dPCR | 154 | 12 | 2 | 25 | 16.2 % |
| 3 |  | 1 | Y | one-step RT-dPCR | 130 | 8.4 | 2.0 | 17 | 13 % |
| 3 |  | 2 | Y | two-step RT-dPCR | 130 | 13 | 2.1 | 27 | 21 % |
| 4 |  | 1 | Y | one-step RT-dPCR | 175 | 12.4 | 2.44 | 31 | 18 % |
| 5 |  | 1 | Y | one-step RT-dPCR | 132.05 | 6.85 | 2 | 13.71 | 10.38 % |
| 6 |  | 1 | Y | one-step RT-dPCR | 357 | 34 | 2 | 68 | 19 % |
| 6 |  | 2 | S | one-step RT-dPCR (RT efficiency corrected) | 496 | 57 | 2 | 114 | 23 % |
| 7 |  | 1 | Y | one-step RT-dPCR | 144.94 | 19.16 | 2 | 38.32 | 26 % |
| 8 |  | 1 | Y | one-step RT-dPCR | 178.4 | 5.2 | 2.57 | 13.5 | 7.6 % |
| 9 |  | 1 | Y | one-step RT-dPCR | 120 | 9.4 | 2.07 | 19 | 16 % |
| 10 |  | 1 | Y | two-step RT-dPCR | 184.6 | 8.8 | 2 | 17.6 | 9.52 % |
| 11 |  | 1 | Y | one-step RT-dPCR (Assay Ref. [17]) | 132.6 | 18.5 | 2.01 | 37.1 | 28.0 % |
| 11 |  | 2 | S | one-step RT-PCR (Assay Ref. [18]) | 113.6 | 15.8 | 2.23 | 35.3 | 31 % |
| 11 |  | 3 | S | one-step RT-dPCR (*gag/pol* duplex assay) | 112.2 | 11.7 | 2.04 | 23.8 | 21 % |

**Key:** $x$ , RNA copy number concentration (*gag* gene); $u$ , standard uncertainty; $k$ , coverage factor; $U$ , expanded uncertainty; *Rel U*, relative expanded uncertainty. \*Nominated results indicated by Y (Yes) or supplementary (S). \*\*For laboratories submitting >1 result, discriminating methodological information is provided in brackets.

### Interlaboratory reproducibility and consistency

Initial review of the results identified Laboratory 12’s results for Study Material 1 and Laboratory 6’s results for Study Material 3 as outliers and technical grounds for this were established (see **Methods Used by Participants**). With the exclusion of the outlying results, the remaining nominated results were found to be consistent with a normal distribution, with interlaboratory reproducibility (s_R_) expressed as % coefficient of variation (%CV) of between 15.3 % and 22.0 % (Table 14).

**Table 14:** Inter-laboratory reproducibility of nominated results.

| Study Material | Number of results | Mean (copies/μL) | SD (copies/μL) | %CV | Normally distributed* |
| --- | --- | --- | --- | --- | --- |
| 1 | 14 | 808 | 173 | 21.4 % | Y |
| 2 | 8 | $5.89 \times 10^9$ | $0.901 \times 10^9$ | 15.3 % | Y |
| 3 | 11 | 142 | 31.1 | 22.0 % | Y |

Values rounded to 3 significant digits. \*Shapiro Wilk test.

A check for overdispersion (Table 15) showed that the interlaboratory variation in results for all three Study Materials was not accounted for in their reported uncertainties.

**Table 15:**
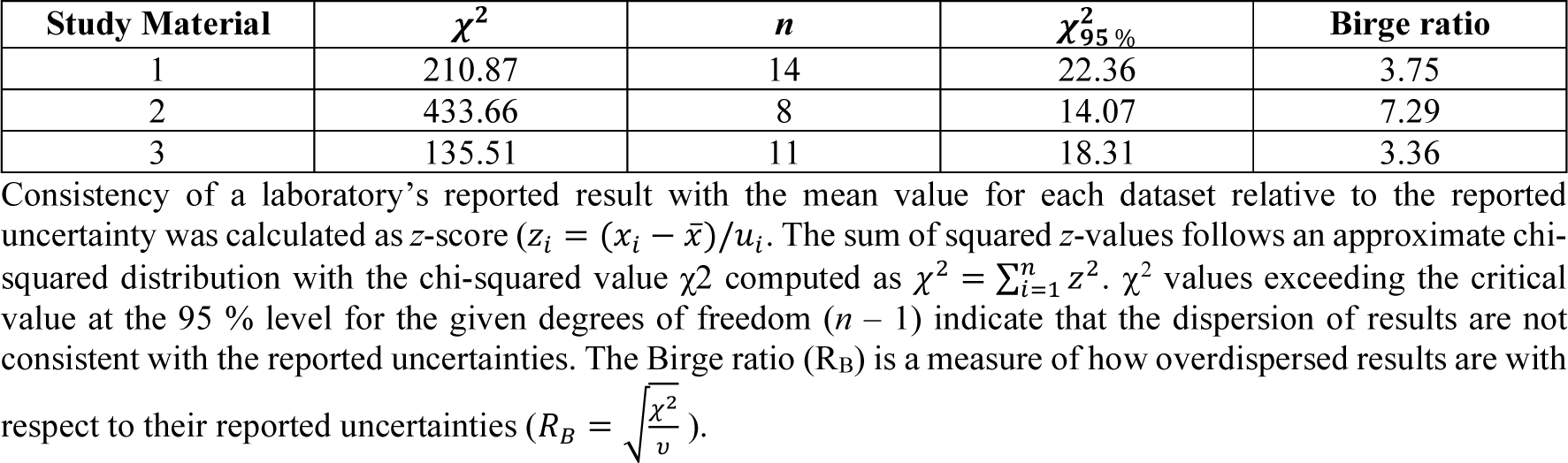
Analysis of Results for overdispersion (chi-squared χ^2^ values).

### Evaluation of assay position and amplicon size on results

To evaluate whether the alternative assays used by participants (Figure 5) influenced the reported RNA concentrations, Study Material 1 and 3 results were plotted against 5’ position in the ≈ 1.5 kb *gag* target sequences for respective materials (Appendix A) and against amplicon size (Figure 9). Results indicated that assay position and amplicon size did not have a systematic effect on reported RNA copy number concentration. Laboratories 4 and 11(S) used the same published assay by Bosman *et al*. [18] (amplicon size 116 bp), however laboratory 4’s values for both Study Materials were higher than Laboratory 11’s supplementary results. This may be due to Laboratory 4 including a heat denaturation step prior to RT-dPCR. In contrast, results for both Study Materials 1 and 3 were very similar for the Kondo *et al*. [17] assay, which had the largest amplicon size of 155 bp, applied by laboratories 5 and 11 (nominated result).

**Figure 9:**
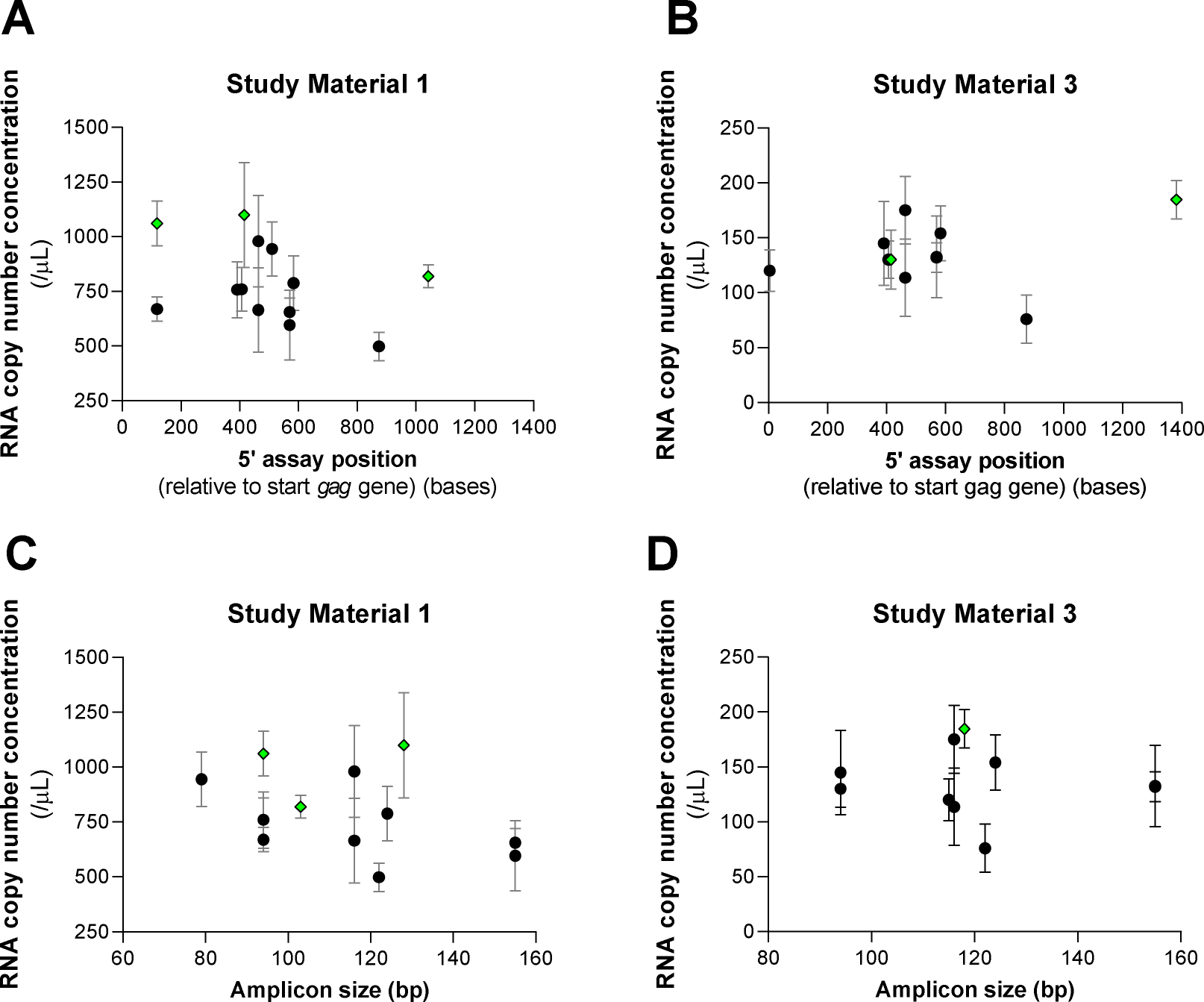
Influence of assay position and amplicon size on Study Material 1 and 3 RNA copy number concentration measurements. One-step RT-dPCR results are shown as black circles; two-step RT-dPCR results are shown as green diamond. Error bars are expanded uncertainties as reported by particpants. Results of laboratories 8 and 9 that utilised a combination multiple RT-dPCR assays (with different 5’ assay positions and amplicon sizes) are not shown for either Study Materials 1 or 3 (Laboratory 8) or Study Material 1 (Laboratory 9).

### Evaluation of trueness: Comparison between RT-dPCR and orthgonal methods

The trueness of RT-dPCR measurements of Study Materials 1 and 2 was evaluated by comparison with results from orthogonal methods applied to Study Material 2.

For the comparison <u>within</u> the Study Material 2 dataset, there were four nominated RT-dPCR results, conducted by diluting Study Material 2 to be measured by dPCR, and four nominated results using orthogonal approaches (Figure 10A). Comparison of the mean values for each group by unpaired *t*-test (equal variances assumed) confirmed that there was no significant difference between groups (Figure 10A) (*p* = 0.46). A similar result was obtained when equal variances were not assumed (Welch’s *t*-test (*p* = 0.46)).

**Figure 10:**
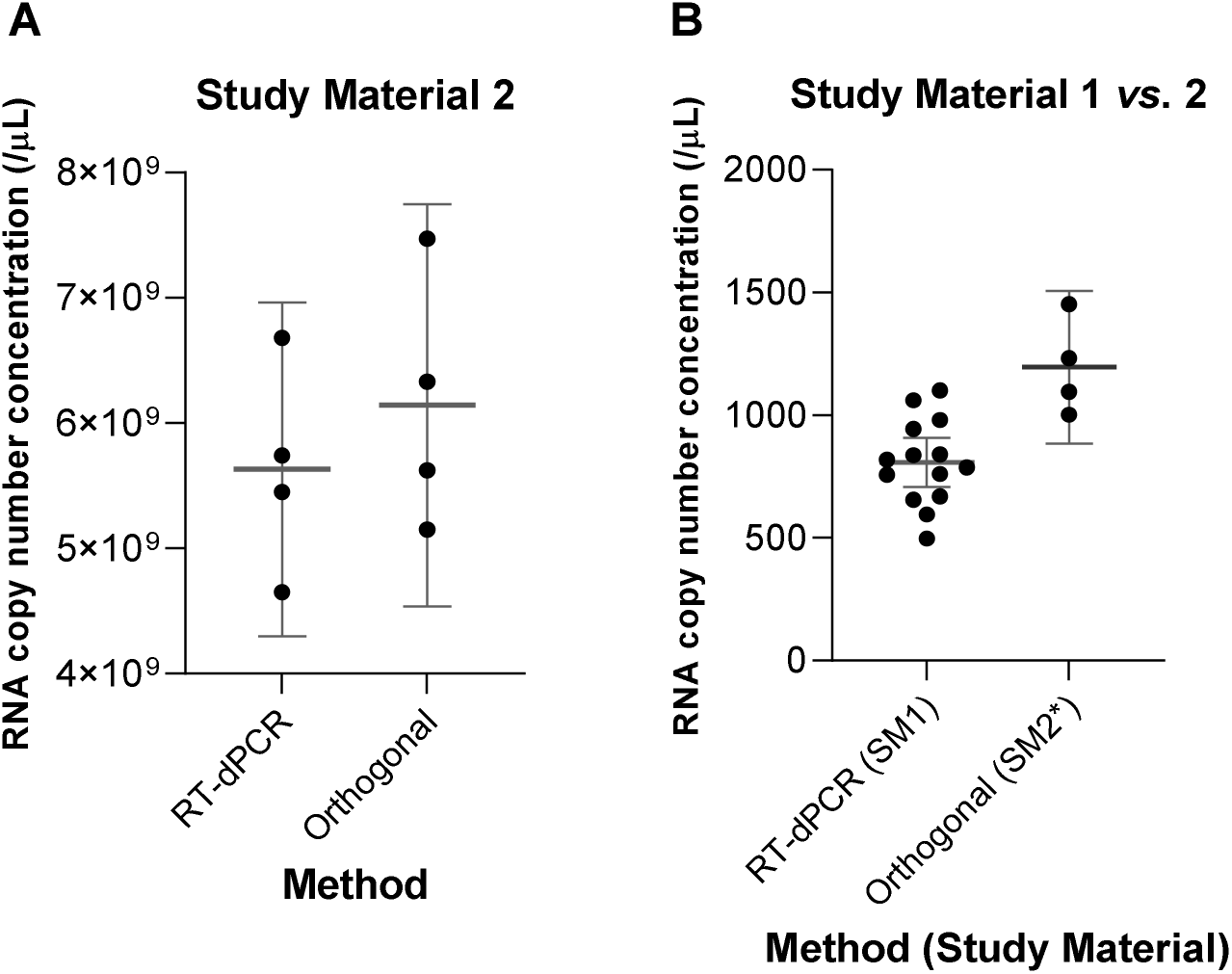
Comparison of RT-dPCR and orthogonal method-based measurements of Study Material 1 and 2 (mean values). Comparison (A) within Study Material 2 results and (B) between Study Material 1 and Study Material 2* results (orthogonal methods only). Individual results are shown by black dots. Mean and 95 % confidence intervals are shown by grey line and error bars respectively. SM2* results extrapolated to Study Material 1 range.

For the comparison <u>between</u> Study Material 1 and 2 results, the Study Material 2 results were scaled according to the dilution factor used in the preparation of Study Material 1 (Table 2) and an additional uncertainty in the gravimetric preparation (relative standard uncertainty of 1.74 %) combined with the laboratory’s reported standard uncertainty (converted to a relative uncertainty) (Table 12). The same coverage factor as reported was applied to calculate a scaled expanded uncertainty. The extrapolated results are shown in Table 16. The Study Material 1 nominated results (*n* = 14) were compared with the extrapolated Study Material 2 results based on orthogonal methods (*n* = 4) and a difference was observed in the group mean values (808 /μL and 1196 /μL respectively) (Figure 10B), which was significant using Welch’s *t*-test (*p* = 0.0195).

**Table 16:** Study Material 2 results extrapolated to Study Material 1 range.

| Lab ID | Result | Nominated | $x^*$ (/μL) | Combined relative $u$ | Combined relative $U$ | $U$ (/μL) |
| --- | --- | --- | --- | --- | --- | --- |
| 2 | 1 | Y | 1300 | 10.61 % | 21.22 % | 276 |
| 3 | 1 | Y | 1002 | 2.76 % | 6.34 % | 64 |
| 6 | 1 | Y | 1117 | 7.18 % | 14.37 % | 160 |
| 6 | 2 | S | 1553 | 9.68 % | 19.36 % | 301 |
| 8 | 1 | Y | 905 | 2.25 % | 5.78 % | 52 |
| 9 | 1 | Y | 1232 | 8.24 % | 16.49 % | 203 |
| 9 | 2 | Y | 1060 | 4.23 % | 9.77 % | 104 |
| 10 | 1 | Y | 1453 | 4.26 % | 8.51 % | 124 |
| 11 | 1 | Y | 1095 | 3.61 % | 11.49 % | 126 |

\*Reported value divided by $5.16 \times 10^6$ .

Further comparison of the Study Material 1 and 2 results in order of Laboratory ID is shown in Figure 11A and pairwise comparison of results in Appendix L (Supplementary information).

**Figure 11:**
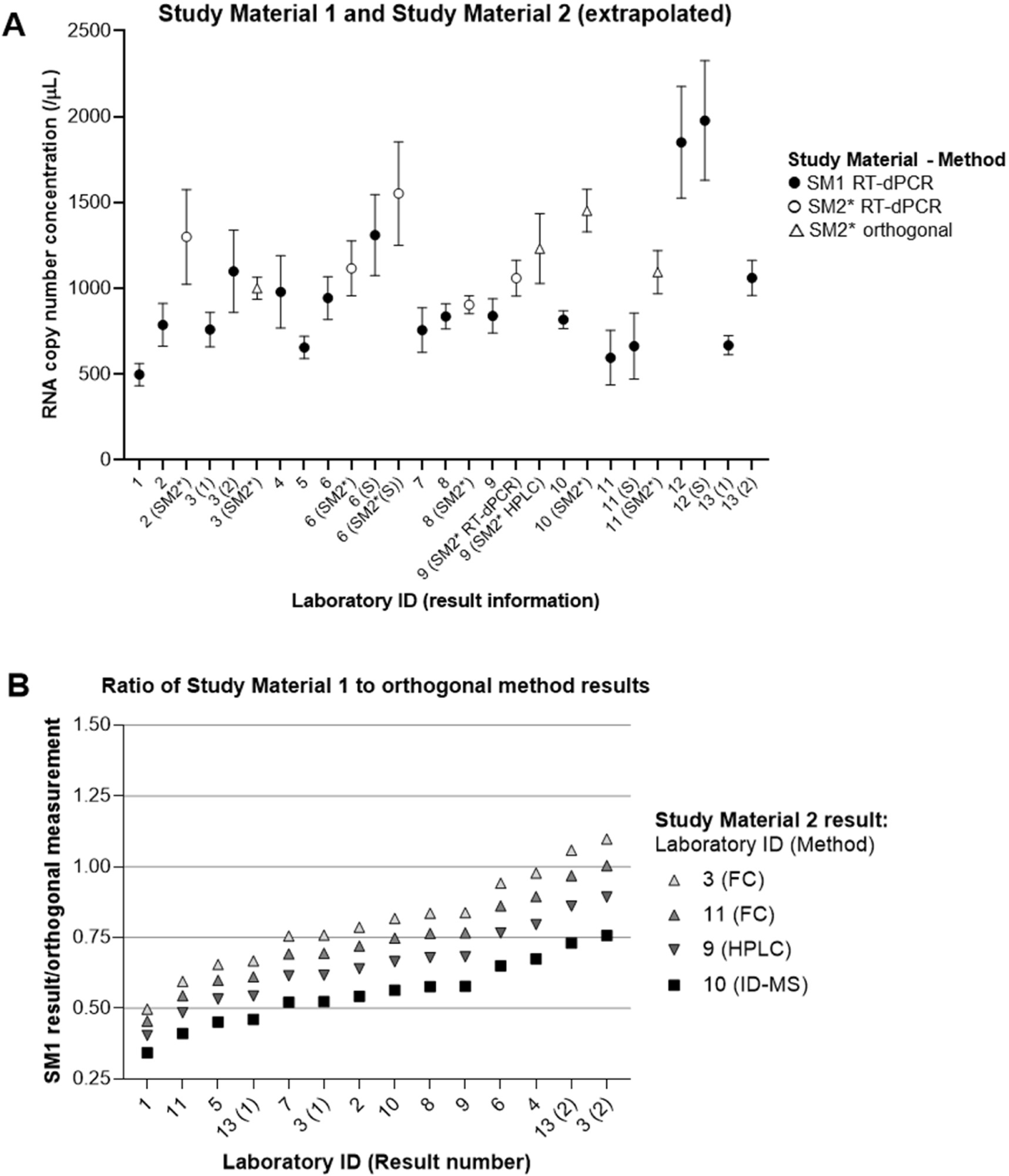
Comparison of Individual Laboratories’ Study Material 1 and 2 results. (A) All results for both Study Materials in order of laboratory ID. Results for Study Material 1 are shown as solid black circles; for Study Material 2 extrapolated results (SM2*), RT-dPCR results for Study Material 2 as open circles and results using orthogonal methods as open triangles. Error bars show expanded uncertainty. (B) Nominated Study Material 1 reported values (*x*) expressed a ratio to each extrapolated Study Material 2* result based on an orthogonal method (legend). Laboratories with multiple nominated or supplementary (S) results indicated by number in parenthesis.

For the seven laboratories who measured both Study Material 1 and Study Material 2, partial consistency between Study Material 1 and Study Material 2 results was observed. Laboratory 3’s Study Material 2 result using single molecule counting was consistent with their two-step RT-dPCR result for Study Material 1 but not their one-step RT-PCR result (*p* < 0.01, Appendix L (Supplementary information)). Laboratory 6’s Study Material 1 and 2 results were consistent for each approach taken (with / without RT efficiency correction; supplementary and nominated results respectively). Laboratory 8’s Study Material 1 and 2 results based on RT-dPCR were mutually consistent. Laboratory 9’s two Study Material 2 results using RT-dPCR and HPLC were mutually consistent, whereas the Study Material 1 result was slightly lower (≈ 1.3 fold to ≈ 1.5 -fold respectively). Laboratory 10’s ID-MS result for Study Material 2 was ≈1.7 -fold higher than their result for Study Material 2 (*p* < 0.001, Appendix L). Laboratory 11’s single molecule counting-based result for Study Material 2 was ≈ 1.8 -fold higher than their RT-dPCR result for Study Material 1 (*p* < 0.001, Appendix L (Supplementary information)).

All Study Material 1 nominated results were compared to each of the four Study Material 2 results based on orthogonal methods to investigate the possible extent of bias affecting RT-dPCR (Figure 11B). The ratio of the Study Material 1 results compared to single molecule flow cytometric counting results ranged from 0.46 to 1.1 (mean 0.81 and 0.74 for laboratory 3 and 11 results respectively). Using HPLC- and ID-MS-based Study Material 2 results as the reference point, Material 1 result ratios were on average 0.66 (range 0.40 to 0.89) and 0.56 (range 0.34 to 0.76).

### Follow-up Analysis (Laboratory 12)

Laboratory 12 undertook additional experiments to investigate the outlying high reported results for Study Material 1 (Table 11 / Figure 7). Two-step RT-dPCR experiments were performed using alternative RT reagents (High capacity RNA to cDNA and Superscript IV RT kits (both Thermo Fisher Scientific)), primers (random octamers, and/or Laboratory 12 ’s assay reverse primer or Laboratory 4’s assay reverse primer) and/or Laboratory 12’s assays or Laboratory 4’s assays (Table H-3, (Supplementary information)). The results (Table 17) were consistent with values reported by other laboratories for Study Material 1 (within 1 SD of the mean, 808 /µL ± 173 /μL (Table 14)).

**Table 17:** Follow-up of Study Material 1 results (Laboratory 12)

| RT reagent | RT priming | dPCR assay (Lab ID) | Result [*gag* copies] (mean) (/μL) | SD [*gag* copies] (/μL) |
| --- | --- | --- | --- | --- |
| High Capacity | Random octamers | 12 | 882 | 10 |
| High Capacity | Random octamers | 4 | 745 | 23 |
| High Capacity | Random octamers and UME's reverse primer | 12 | 846 | 52 |
| SuperScript IV | UME's reverse primer | 12 | 662 | 40 |
| SuperScript IV | NML's reverse primer | 4 | 766 | 125 |

Each result corresponds to a single RT reaction with triplicate dPCR assays (QX200) ( $n = 3$ ).

Further replicate experiments were performed using the High capacity RNA to cDNA kit with random octamer RT priming which were sent to the coordinator in May 2020 (Appendix K Table K-1, (Supplementary information)). These results (*x* = 945, *U* = 286 /μL) were also consistent with the main results for Study Material 1.

### Follow-up Analysis of Study Material 2 purity (Laboratory 10)

It was hypothesised that relative differences in RNA copy number concentration of Study Material 2 measured by chemical analysis methods (ID-MS and, to a lesser extent, HPLC) compared to RT-dPCR of the same material and Study Material 1 may be due to the sensitivity of mass spectrometry and LC approaches to nucleic acid impurities which lack the target sequence. It was also noted that any variation between the expected MW of the *in vitro* transcribed RNA (Appendix A) and the true value will also lead to a discrepancy in approaches which utilise MW in calculations to convert mass concentration to copy number concentration.

To follow up these issues, Laboratory 10 received an additional five units of Study Material 2 in June 2020 and investigated size impurities using Ultra Performance Liquid Chromatography (UPLC) (Waters H-class system)-based size exclusion chromatography (SEC) (experimental information recorded in Table H-7). The NMIJ CRM panel 6204-b (1000-A and 500-A standards) and Nucleotide monomers (AMP) were used to establish the linearity of detection over the range 0 ng/g to 40 ng/g. Figure 12 shows chromatograms of Study Material 2 analysed in isolation and mixed with a single-stranded RNA (ssRNA) ladder of different sizes).

**Figure 12:**
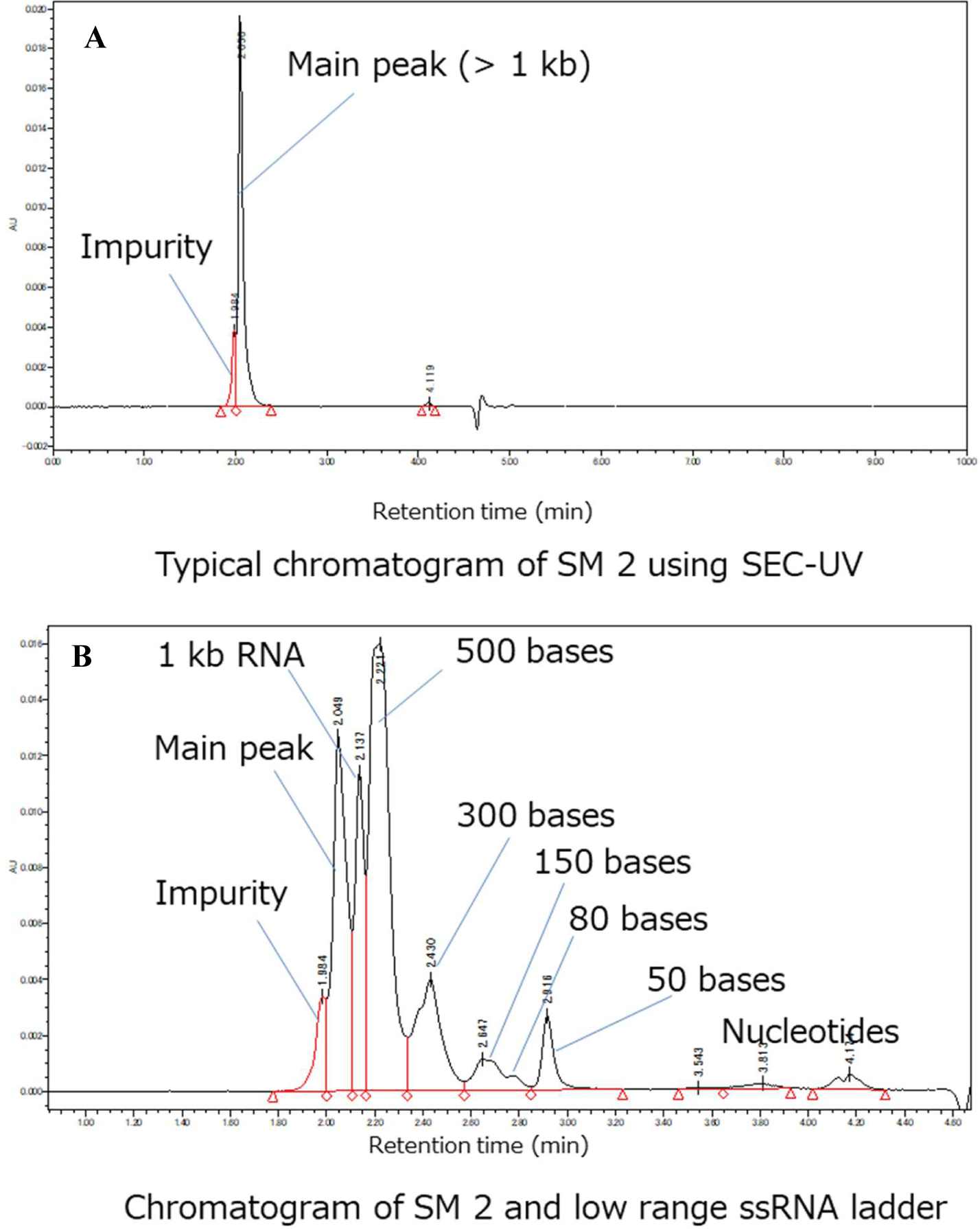
UPLC analysis of Study Material 2 fragment size impurities. Chromatograms are shown for (A) Study Material 2 analysed on its own and (B) mixed with a RNA ladder (1 kb, 500 bases, 300 bases, 150 bases, 80 bases and 50 bases) and nucleotides

No small molecular impurities were detected in Study Material 2, confirming the absence of fragmented or truncated IVT products or carryover of nucleotide triphosphates from the IVT reaction. A secondary peak of larger size than the main peak was observed (Figure 12). The relative concentration of the larger fragment compared to the main peak was quantified by area under the curve (AUC) (Table 18) and estimated to constitute on average 16.6 % ±5.8 % (k=2) of the sample *by mass*. Pre-treatment of Study Material 2 with S1 nuclease which is specific to single-stranded nucleic acids lead to the appearance of monomers in the UPLC chromatogram (Appendix K, (Supplementary information)). This indicated that the impurity consisted of RNA (rather than plasmid DNA contamination).

**Table 18:** Results of UPLC analysis of Study Material 2 fragment size impurities.

| **Unit Number** | **59** | **69** | **71** | **87** | **105** |
| --- | --- | --- | --- | --- | --- |
| Main peak area (n = 5) | 65089 | 73481 | 75552 | 71138 | 76707 |
| SD | 2443 | 1059 | 755 | 585 | 664 |
| RSD | 3.8 % | 1.4 % | 1.0 % | 0.8 % | 0.9 % |
| Impurity peak area (n= 5) | 16506 | 12379 | 11667 | 17333 | 14018 |
| SD | 583 | 116 | 270 | 104 | 88 |
| RSD | 3.5 % | 0.9 % | 2.3 % | 0.6 % | 0.6 % |
| Ratio Impurity / Main peak | 25.4 % | 16.8 % | 15.4 % | 24.4 % | 18.3 % |
| % Impurity peak area / total AUC | 20.2 % | 14.4 % | 13.4 % | 19.6 % | 15.5 % |

The UPLC chromatogram profile observed by Laboratory 10 had some similarity to that observed by Laboratory 9 in their participant HPLC-UV analysis; with a small “shoulder” observed to the left of the main peak, which was not as distinct as the UPLC chromatogram. This area was not included in the AUC calculations for Study Material 2 result reporting (Appendix K, (Supplementary information)), which may explain why the Laboratory 9 value for Study Material 2 was lower than Laboratory 10’s result. Review of capillary electrophoresis data by the coordinator also indicated the the presence of peaks with sizes of approximately ≈ 2300 (expected size) and ≈ 2500 nucleotides (Appendix B, Table B-1 (Supplementary information)). However the proportion of the sample in each peak was the inverse of the UPLC results, with the AUC data indicating that the larger peak constituted 81 % of the sample compared to 19 % for the smaller peak. Standard denaturing agarose gel electrophoresis did not show multiple bands (Appendix B, (Supplementary information)). Laboratory 3 analysed Study Material 2 using PAGE and also observed multiple peaks (Appendix K, (Supplementary information)). Residual secondary structure may be a factor in the discrepancy between the Bioanalyzer and the UPLC/HPLC results.

### Discussion of Results

CCQM-P199 evaluated RT-dPCR as a candidate RMP methodology for quantification of RNA copy number concentration, as well as testing candidate RMPs for the HIV-1 target gene which constitutes the model system and analyte (*gag* gene). As well as measuring interlaboratory reproducibility and consistency, trueness was investigated by means of comparison with alternative non-PCR based techniques.

Both one-step and two-step RT-dPCR approaches were used for analysis of Study Materials 1 and 3. Two-step RT-dPCR results tended to be higher than those using one-step RT-dPCR methodology. This was evident in Laboratory 3’s and Laboratory 13’s parallel one- and two-step results for Study Material 1, however Laboratory 3’s one- and two-step values were the same for Study Material 3 (Laboratory 13 did not measure this material). Laboratory 10’s two-step result using an alternative dPCR platform (QS3D) was close to the mean value for Study Material 1, and, although their Study Material 3 result was ranked highest (excluding Laboratory 12’s results), the value was similar to and consistent with a number of results applying the more common one-step RT-dPCR format with the QX100/200 dPCR platform. Moreover, the higher results for Study Material 1 were found to be more consistent with the scaled results for Study Material 2 based on orthogonal approaches. These comparisons suggested that sources of negative bias may affect some of the RT-dPCR results reported for Study Material 1 results. Comparing RT-dPCR results to the results of single molecule counting which directly measures the measurand of the study (i.e. concentration of molecular entities as opposed to HPLC and ID-MS which indirectly calculate molar concentration and copy number), the estimate of bias was between -19 % and -26 % (ratios of 0.81 and 0.74).

The causes of negative bias affecting RT-dPCR measurements may include RT efficiency and partition volume. In a one-step RT-dPCR format, it is possible that buffer compatibility for RT and *Taq* polymerase enzymes is less than ideal and this could lead to a negative bias in RT efficiency (i.e. <100 % RNA template being converted to cDNA) or dPCR efficiency (i.e. <100 % dPCR partitions containing cDNA producing detectable amplification). In addition, application of dPCR partition volume values which are higher than the true values (in the case of laboratories not directly measuring partition volume) may contribute to an underestimation of RNA copy number concentration. Some laboratories applied the default value of 0.85 nL for the QX100/200 system (three institutes) which was higher than the majority of partition volumes of the One-step RT-ddPCR Advanced kit for Probes measured directly by other participants (Appendix H, (Supplementary information)). It should be noted that the lowest report results for P199 Study Materials 1 and 3 (1.6-fold and 1.9 -fold lower than interlaboratory mean values respectively) were from Laboratory 1 whose shipment experienced significant delays (Table 7), therefore the low values may be in part be attributable to degradation of the RNA rather than technical factors.

Considering the possibility of positive bias affecting orthogonal methods, it is of note that the orthogonal method values for Study Material 2 are all lower than the Laboratory 12 Study Material 1 results, where a multiple cDNA template per RNA copies may have occurred, leading to results which are ≈ 2 -fold higher than the mean value for this material. Therefore range of systematic error is expected to be <2 -fold between RT-dPCR and orthogonal approaches. Follow-up investigation of Study Material 2 purity by Laboratory 10 suggested possible causes of the Laboratory 9 HPLC and Laboratory 10 ID-MS results being higher than some RT-dPCR measurements: discrepancy of the actual size/MW of the IVT material and/or presence of impurities which do not contain the target sequence. The latter hypothesis is yet to be tested, but could be established by applying NGS approaches similar to those used by Gholamalipour *et al*. for small IVT molecules [19]. The MW discrepancy alone could account for 10 % difference and reflects an important consideration when deploying established SI-traceable methods to macromolecules like DNA.

Inter-laboratory reproducibility for RT-dPCR measurements was approximately ≈ 20 % (%CV) which is similar to that observed for previous studies evaluating dPCR as a candidate RMP for DNA measurements [20–22].

Inter-laboratory consistency analyses suggest that reported measurement uncertainties for RT-dPCR were not large enough to account for all sources of uncertainty causing inter-laboratory dispersion to be greater than predicted based on within-laboratory estimation alone. Factors which were included in measurement uncertainty budgets by the majority of laboratories included precision and partition volume, whilst between assay variation was not evaluated by all laboratories (Table I-1, (Supplementary information)). As noted in the preceding paragraphs, lack of an accepted approach for testing and correction for RT efficiency also affected inclusion of this factor in RT-dPCR uncertainty estimation.

### PILOT STUDY CONSENSUS REFERENCE VALUE

Nominated results were included in the calculation of consensus reference values (RV) for all Study Materials. Multiple nominated results from the same institution were considered independently in the consensus RV calculations in the case of two alternative methods being applied: one-step and two-step RT-dPCR results (Laboratory 3 and Laboratory 13) for Study Materials 1 (both institutes) and 3 (Laboratory 3 only; Laboratory 13 did not analyse this material), and HPLC and RT-dPCR results for Study Material 2 (Laboratory 9). Results were excluded from the consensus RV calculations where a technical error or issue was considered to be the cause of outlying results. For Study Material 1, Laboratory 12’s nominated result was not included due to the substitution of an RT-PCR kit for an RT kit in the two-step RT-dPCR process, which was considered to be the cause of their result being approximately 2 -fold higher than the mean. For Study Material 3, Laboratory 6’s nominated result was not included due to the identification of multiple priming sites for the reverse primer in the assay present in the target sequence in this material, which was hypothesised to be the reason for their nominated result being approximately 2 -fold higher than the average.

Candidate RVs and uncertainties (Table 19, Figure 13) were calculated using approaches based only on variation between laboratories’ reported values (arithmetric mean, median and Huber Proposal 2) and estimators weighted inversely to the laboratories’ reported uncertainties with an excess variance component related to inter-laboratory dispersion (weighted mean with Birge ratio; DerSimonian-Laird, Mandel Paule). Consistency analysis of laboratories’ results (see Discussion of Results) indicate significant over-dispersion relative to laboratories’ reported uncertainties and smaller uncertainty budgets which may not be due to higher accuracy in the underlying methods; consequently weighted approaches are not considered appropriate. For example, the weighted mean for Study Material 1 is skewed by the lower uncertainties reported by Laboratory 1, Laboratory 5 and Laboratory 13 (1) and likewise, the low uncertainty for Laboratory 8’s Study Material 2 result leads to the weighted mean being lower than the other RV estimators. Although the DSL and Mandel-Paule estimators do not show this problem, the lack of consistency in factors included in reported uncertainties suggest that estimators which include these are not appropriate conceptually.

**Figure 13:**
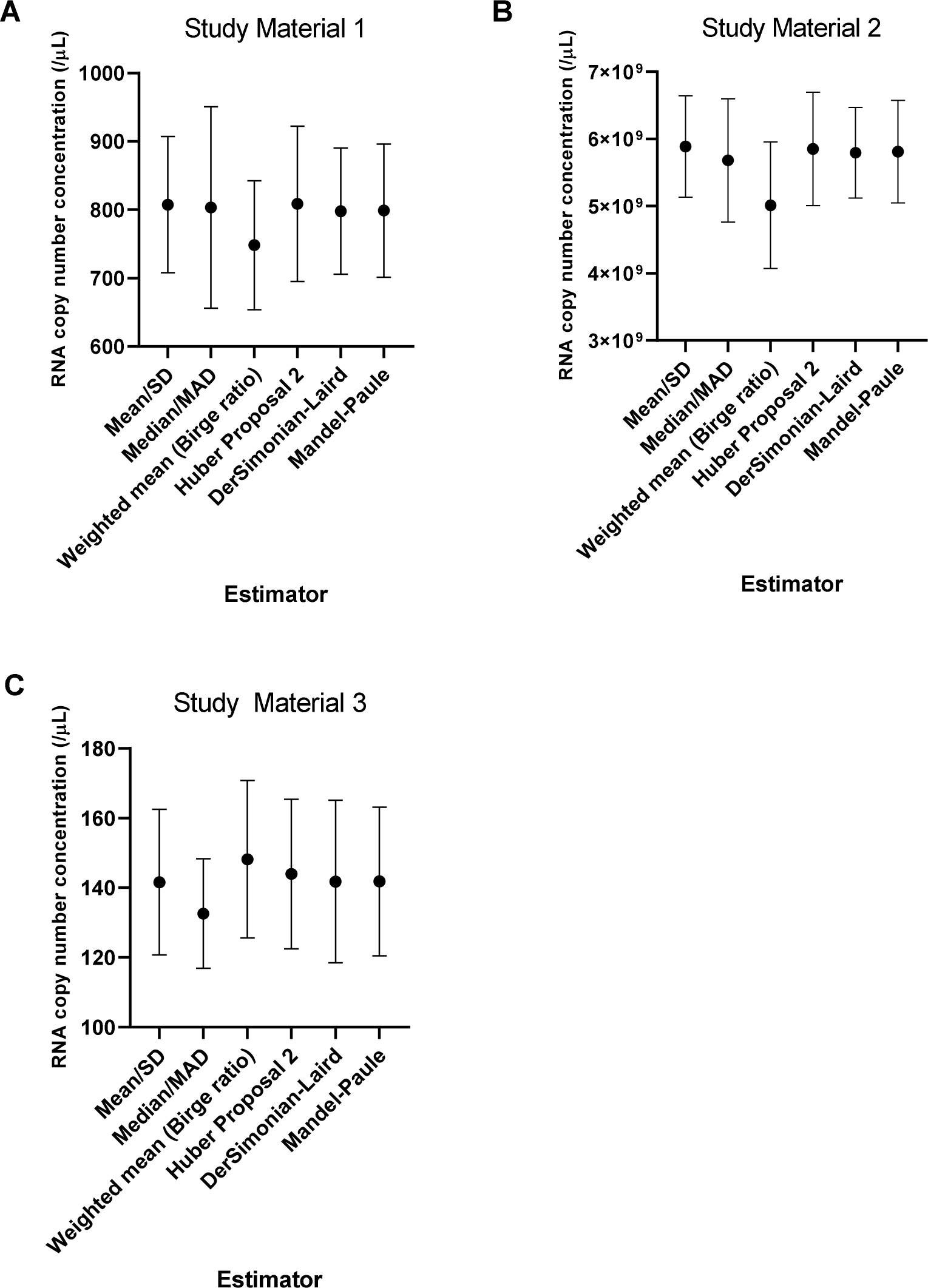
Comparison of alternative estimators for CCQM-P199 consensus Reference Values. Dot shows consensus RV. Error bars show expanded uncertainty (95 % confidence interval).

**Table 19:** Candidate consensus Reference Values estimators: *gag* RNA copy number concentration.

| **Study Material 1** |  |  |  |  |  |
| --- | --- | --- | --- | --- | --- |
| **Estimator** | **RV (/μL)** | ***u* (/μL) Note 2** | ***m*** | ***k*** | ***U* (/μL) Note 1** |
| Arithmetic mean* | 808 | 46.168 | 14 | 2.1604 | 100 |
| Median/MADe | 803 | 68.285 | 14 | 2.1604 | 148 |
| Weighted mean with Birge ratio | 748 | 43.599 | 14 | 2.1604 | 95 |
| Huber Proposal 2 | 809 | 52.597 | 14 | 2.1604 | 114 |
| DerSimonian-Laird (DSL) | 798 | 42.646 | 14 | 2.1604 | 93 |
| Mandel-Paule | 799 | 45.046 | 14 | 2.1604 | 98 |
| **Study Material 2** |  |  |  |  |  |
| **Estimator** | **RV (×10<sup>9</sup>/μL)** | ***u* (×10<sup>9</sup>/μL)** | ***m*** | ***k*** | ***U* (×10<sup>9</sup>/μL)** |
| Arithmetic mean* | 5.89 | 0.318 | 8 | 2.3646 | 0.76 |
| Median/MADe | 5.68 | 0.388 | 8 | 2.3646 | 0.92 |
| Weighted mean with Birge ratio | 5.01 | 0.398 | 8 | 2.3646 | 0.95 |
| Huber Proposal 2 | 5.85 | 0.357 | 8 | 2.3646 | 0.85 |
| DerSimonian-Laird (DSL) | 5.79 | 0.286 | 8 | 2.3646 | 0.68 |
| Mandel-Paule | 5.81 | 0.322 | 8 | 2.3646 | 0.77 |
| **Study Material 3** |  |  |  |  |  |
| **Estimator** | **RV (/μL)** | ***u* (/μL)** | ***m*** | ***k*** | ***U* (/μL)** |
| Arithmetic mean* | 142 | 9.388 | 11 | 2.2281 | 21 |
| Median/MADe | 133 | 7.059 | 11 | 2.2281 | 16 |
| Weighted mean with Birge ratio | 148 | 10.143 | 11 | 2.2281 | 23 |
| Huber Proposal 2 | 144 | 9.637 | 11 | 2.2281 | 22 |
| DerSimonian-Laird (DSL) | 142 | 10.456 | 11 | 2.2281 | 24 |
| Mandel-Paule | 142 | 9.587 | 11 | 2.2281 | 22 |

**Note 1:** RV and expanded uncertainties for Study Materials 1 and 3 rounded to the nearest whole copy (outwards for expanded uncertainty). Standard uncertainties shown to 3 d.p.
**Note 2:** No allowance for material homogeneity as between unit variation was low (Table 3) combined with 4 units of each material being provided to participants for analysis (effective standard uncertainty of $\leq 1.1$ %), therefore the contribution to the consensus RV uncertainty due to inhomogeneity is negligible ( $<4$ % of total variance).
**Note 3:** *m*, number of nominated results used to calculate RV. Effective degrees of freedom (*m*-1) was used to calculate *k*. Coverage factor for 95 % confidence level calculated.
**Note 4:** Huber Proposal 2 is a robust estimator taking no account of reported uncertainty. Typically behaves between median and mean.
\*Recommended estimator for P199 consensus RV.

As inter-laboratory variation dominates, together with the study result datasets used for consensus RV calculation being normally distributed in all cases, without any statistical outliers, the mean/SEM estimator was recommended by the coordinators and agreed by study participants to form the RV for all three Study Materials. The agreed RVs and uncertainties are compared to the study results for each Study Material in Figure 14.

**Figure 14:**
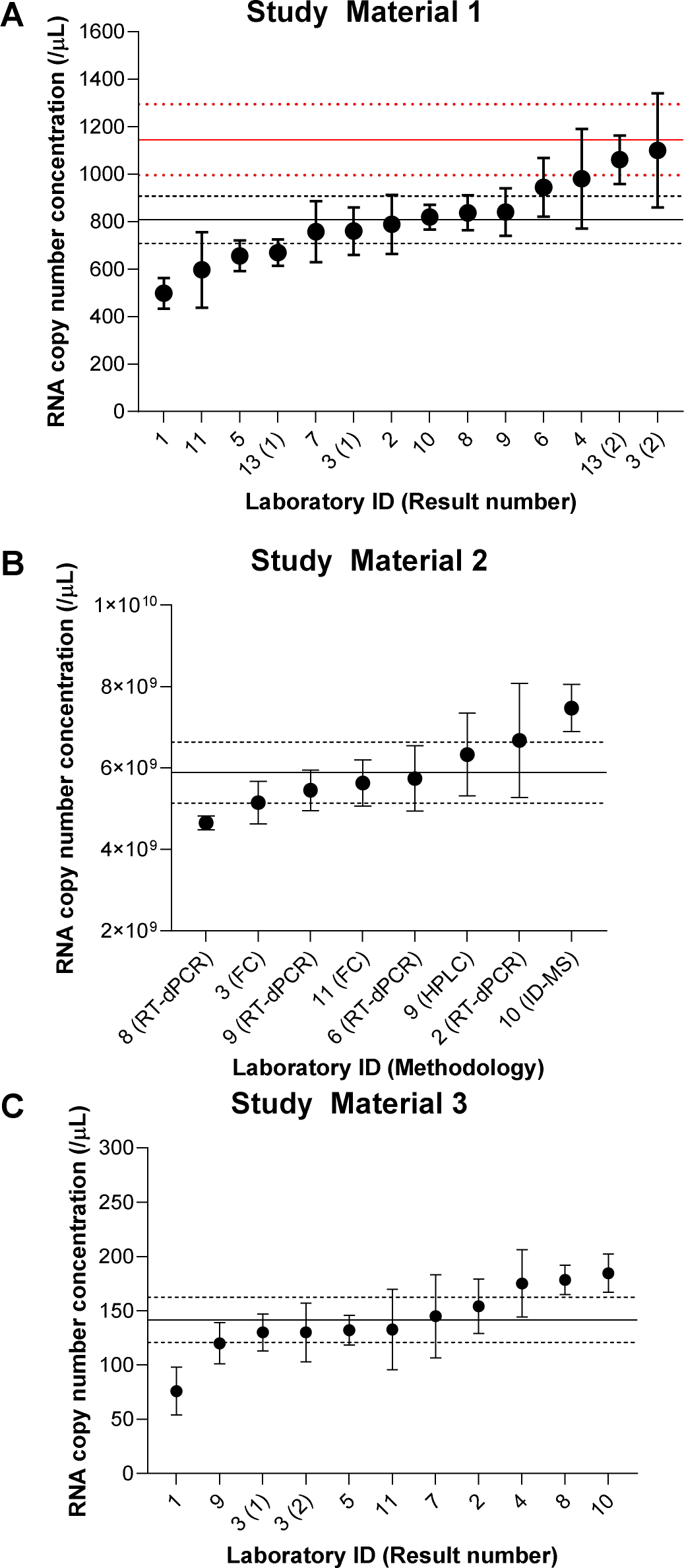
CCQM-P199 study results compared to agreed consensus RVs. Consensus RVs based on arithmetic mean (solid line; expanded uncertainty, dotted lines) are compared to reported results in ascending order. Error bars show laboratories’ reported expanded uncertainties. Consensus mean and expanded uncertainty (solid and dotted lines, respectively) for Study Material 2 extrapolated to Study Material 1 is shown in red in A.

## CONCLUSIONS

CCQM-P199 assessed participants’ capabilities for targeted RNA copy number concentration measurements and viral gene quantification using candidate higher order methodologies including those based on enumeration: RT-dPCR and single molecule flow cytometry; and chemical analysis approaches: HPLC (traceable to RNA CRMs defined in mass and molar concentration) and ID-MS (traceable to nucleotide standards of defined purity). Enzymatically synthesized RNA molecules (at two concentrations approximately 6 orders of magnitude apart) and purified RNA from whole virus were analysed, and RNA copy number concentration reported in copies per µL. RT-dPCR analysis demonstrated comparable quantification of HIV- 1 *gag* between participants, for both the *in vitro* transcribed RNA and the whole viral RNA material, with results within 22 % coefficient of variation (CV) or less.

The majority of measurements were performed using a one-step RT-dPCR approach which was common throughout the study, with some participants performing two-step RT-dPCR. Results obtained by two-step RT-dPCR tended to be higher than one-step results, possibly associated with a negative bias for the latter approach. RT-dPCR performance may be influenced by numerous factors including assay choice, reverse transcriptase efficiency and template type [14, 23] as well as partition volume. In terms of assay choice, participants had to select or design their own assays based on the HIV-1 *gag* gene sequences provided for the materials (spanning ≈ 1.5 kb). The HIV-1 *gag* gene was chosen in this study to represent a commonly used target for clinical viral load measurement. Although assay choice was found to have no systematic impact on result in this study, sequence-specific effects should be taken into consideration when comparing RNA copy numbers for highly divergent HIV-1 genomes, as different PCR assays can generate significantly different values [24].

Evaluation of trueness with orthogonal methods estimates that differences between techniques are less than 2 -fold. The counting-based methods, which were not affected by study material purity issues, are in 1.3 -fold to 1.4 -fold agreement. This is consistent with measurement uncertainty for RT-dPCR based on inter-laboratory reproducibility (expanded uncertainty of 40 %). Therefore, determining the most appropriate measurement uncertainty for RT-dPCR results should take into account not only between-laboratory agreement for RT-dPCR measurements but also their agreement with orthogonal techniques (Figure 14A). This performance is fit for purpose in supporting standardisation and harmonisation of clinical viral load measurements which can vary by several orders of magnitude. However, to reduce measurement uncertainty and support viral load quantification in whole virus biological standards and materials, further evaluation and the development and testing of appropriate controls for RT efficiency is needed along with studies that will also explore extraction of the viral genome from biological specimens.

## Supporting information

CCQM P199 supplementary file Appendix A

CCQM P199 supplementary file Appendix B

CCQM P199 supplementary file Appendix C

CCQM P199 supplementary file Appendix D

CCQM P199 supplementary file Appendix E

CCQM P199 supplementary file Appendix F

CCQM P199 supplementary file Appendix G

CCQM P199 supplementary file Appendix H

CCQM P199 supplementary file Appendix I

CCQM P199 supplementary file Appendix J

CCQM P199 supplementary file Appendix K

CCQM P199 supplementary file Appendix L

## ACKNOWLEDGEMENTS

The study coordinators thank the participating laboratories for providing the requested information used in this study. NML would like to thank Stephen Ellison, Simon Cowen and Stephen Nyangoma for their input with statistical analysis of the study results. Copyright notice for Figure 1: Copyright Triad National Security, LLC. All Rights Reserved.

Points of view in this document are those of the authors and do not necessarily represent the official position or policies of the U.S. Deparent of Commerce. Certain commercial software, instruments, and materials are identified in order to specify experimental procedures as completely as possible. In no case does such identification imply a recommendation or endorsement by NIST, nor does it imply that any of the materials, instruments, or equipment identified are necessarily the best available for the purpose.

## Footnotes

1 The molecular biology community express amount of substance concentration with units of molarity (M) (SI units: mmol/L).

## REFERENCES

[1] Mellors JW, Munoz A, Giorgi JV, Margolick JB, Tassoni CJ, Gupta P, et al. Plasma viral load and CD4+ lymphocytes as prognostic markers of HIV-1 infection. Annals of Internal Medicine (1997);126. 10.7326/0003-4819-126-12-199706150-00003

[2] Steinmetzer K, Seidel T, Stallmach A, Ermantraut E. HIV load testing with small samples of whole blood. J Clin Microbiol (2010);48(8):2786–92. 10.1128/JCM.02276-09

[3] Rouet F, Ménan H, Viljoen J, Ngo-Giang-Huong N, Mandaliya K, Valéa D, et al. In-house HIV-1 RNA real-time RT-PCR assays: principle, available tests and usefulness in developing countries. Expert Review of Molecular Diagnostics (2008);8(5):635–50. 10.1586/14737159.8.5.635

[4] Gullett JC, Nolte FS. Quantitative nucleic acid amplification methods for viral infections. Clinical Chemistry (2015);61(1):72–8. 10.1373/clinchem.2014.223289

[5] ISO 17511:2020. In vitro diagnostic medical devices — Requirements for establishing metrological traceability of values assigned to calibrators, trueness control materials and human samples. https://www.iso.org/standard/69984.html

[6] Prescott G, Hockley J, Atkinson E, Rigsby P, Morris C. International collaborative study to establish the 4th WHO international standard for HIV-1 NAT assays. (2017);WHO/BS/2017.2314. https://www.who.int/publications/m/item/WHO-BS-2017.2314

[7] Angus B, Brook G, Awosusi F, Barker G, Boffito M, Das S, et al. BHIVA guidelines for the routine investigation and monitoring of adult HIV-1-positive individuals (2019 interim update). https://www.bhiva.org/monitoring-guidelines

[8] HIV sequence database. HIV-1 Gene Map; (2017). Available from: https://www.hiv.lanl.gov/content/sequence/HIV/MAP/landmark.html. [Accessed 25th May 2021.

[9] Falak S, Macdonald R, Busby EJ, O’Sullivan DM, Milavec M, Plauth A, et al. An assessment of the reproducibility of reverse transcription digital PCR quantification of HIV-1. Methods (2021);201:34–40. 10.1016/j.ymeth.2021.03.006

[10] Amendola A, Sberna G, Forbici F, Abbate I, Lorenzini P, Pinnetti C, et al. The dual-target approach in viral HIV-1 viremia testing: An added value to virological monitoring? PloS one (2020);15(2):e0228192. 10.1371/journal.pone.0228192

[11] Yoo HB, Park SR, Dong L, Wang J, Sui Z, Pavsic J, et al. International Comparison of Enumeration-Based Quantification of DNA Copy-Concentration Using Flow Cytometric Counting and Digital Polymerase Chain Reaction. Anal Chem (2016);88(24):12169–76. 10.1021/acs.analchem.6b03076

[12] Ma F, Li Y, Tang B, Zhang CY. Fluorescent Biosensors Based on Single-Molecule Counting. Accounts of chemical research (2016);49(9):1722–30. 10.1021/acs.accounts.6b00237

[13] Yoo HB, Park SR, Hong KS, Yang I. Precise RNA Quantification by Counting Individual RNA Molecules Using High-Sensitivity Capillary Flow Cytometry. Anal Chem (2022). 10.1021/acs.analchem.1c04355

[14] Schwaber J, Andersen S, Nielsen L. Shedding light: The importance of reverse transcription efficiency standards in data interpretation. Biomol Detect Quantif (2019);17:100077. https://www.sciencedirect.com/science/article/pii/S2214753517302188

[15] Joint Committee for Guides in Metrology. International vocabulary of metrology – Basic and general concepts and associated terms (VIM). (2012). Available from: https://jcgm.bipm.org/vim/en/index.html. [Accessed 08/03/2024.

[16] Gall A, Morris C, Kellam P, Berry N. Complete Genome Sequence of the WHO International Standard for HIV-1 RNA Determined by Deep Sequencing. Genome Announcements (2014);2(1). 10.1128/genomeA.01254-13

[17] Kondo M, Sudo K, Tanaka R, Sano T, Sagara H, Iwamuro S, et al. Quantitation of HIV-1 group M proviral DNA using TaqMan MGB real-time PCR. Journal of Virological Methods (2009);157. 10.1016/j.jviromet.2008.12.006

[18] Bosman KJ, Nijhuis M, van Ham PM, Wensing AM, Vervisch K, Vandekerckhove L, De Spiegelaere W. Comparison of digital PCR platforms and semi-nested qPCR as a tool to determine the size of the HIV reservoir. Scientific Reports (2015);5:13811. 10.1038/srep13811

[19] Gholamalipour Y, Karunanayake Mudiyanselage A, Martin CT. 3’ end additions by T7 RNA polymerase are RNA self-templated, distributive and diverse in character-RNA-Seq analyses. Nucleic Acids Research (2018);46(18):9253–63. https://www.ncbi.nlm.nih.gov/pmc/articles/PMC6182178/

[20] Whale AS, Jones GM, Pavšič J, Dreo T, Redshaw N, Akyurek S, et al. Assessment of Digital PCR as a Primary Reference Measurement Procedure to Support Advances in Precision Medicine. Clinical Chemistry (2018);64(9):1296–307. 10.1373/clinchem.2017.285478

[21] Milavec M, Pavšič J, Bogožalec Košir A, Jones GM, O’Sullivan DM, Devonshire AS, et al. The performance of human cytomegalovirus digital PCR reference measurement procedure in seven external quality assessment schemes over four years. Methods (2021). https://www.sciencedirect.com/science/article/pii/S1046202321000906

[22] Burke D, Devonshire AS, Pinheiro LB, Jones GM, Griffiths KR, Gonzalez AF, et al. Standardisation of cell-free DNA measurements: An International Study on Comparability of Low Concentration DNA Measurements using cancer variants. bioRxiv (2023):2023.09.06.554514. https://www.biorxiv.org/content/biorxiv/early/2023/09/06/2023.09.06.554514.full.pdf

[23] Kiselinova M, Pasternak AO, De Spiegelaere W, Vogelaers D, Berkhout B, Vandekerckhove L. Comparison of Droplet Digital PCR and Seminested Real-Time PCR for Quantification of Cell-Associated HIV-1 RNA. PloS one (2014);9(1):e85999. 10.1371%2Fjournal.pone.0085999

[24] Carneiro J, Resende A, Pereira F. The HIV oligonucleotide database (HIVoligoDB). Database (Oxford) (2017)(1):bax005. https://www.ncbi.nlm.nih.gov/pmc/articles/PMC5502365/

