## Supplementary material for "CCQM-P199: Interlaboratory comparability study of HIV-1 RNA copy number quantification": CCQM P199 supplementary file Appendix A

#### APPENDIX A: Sequence information

#### 1. Description of Study Materials

##### Study Materials 1 and 2

Materials 1 and 2 are composed of *in vitro* transcribed (IVT) RNA in (i) human Jurkat cell line total RNA (Material 1) or (ii) buffered solution (Material 2). The IVT RNA sequence is derived from K03455.1 Human immunodeficiency virus type 1 (HXB2) complete HIV1/HTLV-III/LAV reference genome (positions 1 to 2,257). Nucleotides 1 to 9 of the below sequence represent the G-terminal of the T7 promoter used for *in vitro* transcription of RNA and a *Not*I restriction site. The sequence terminates at a *Bsu*36I restriction site on the anti-sense strand. Note: the anti-sense DNA strand serves as the template for *in vitro* transcription. The total sequence length is 2,266 nucleotides. Total molecular weight (MW) of the single stranded RNA transcript was estimated to be 734035.2 g/mol.

```
GGCGGCCGCUGGAAGGGCUAAUUCACUCCCAACGAAGACAAGAUAUCCUUGAUCUGUGGA
UCUACCACACACAAGGCUACUUCCCUGAUUAGCAGAACUACACACCAGGGCCAGGGAUCAG
AUAUCCACUGACCUUUGGAUGGUGCUACAAGCUAGUACCAGUUGAGCCAGAGAAGUUAGAA
GAAGCCAACAAAGGAGAGAACACCAGCUUGUUACACCCUGUGAGCCUGCAUGGAAUGGAUG
ACCCGGAGAGAGAAGUGUUAGAGUGGAGGUUUGACAGCCGCCUAGCAUUUCAUCACAUGG
CCCGAGAGCUGCAUCCGGAGUACUUCAAGAACUGCUGACAUCGAGCUUGCUACAAGGGAC
UUUCCGCUGGGGACUUUCCAGGGAGGCGUGGCCUGGGCGGGACUGGGGAGUGGCGAGCC
CUCAGAUCCUGCAUAUAAGCAGCUGCUUUUUGCCUGUACUGGGUCUCUCUGGUUAGACCA
GAUCUGAGCCUGGGAGCUCUCUGGCUAACUAGGGAACCCACUGCUUAAGCCUCAAUAAAG
CUUGCCUUGAGUGCUUCAAGUAGUGUGUGCCCGUCUGUUGUGUGACUCUGGUAACUAGAG
AUCCCUCAGACCCUUUUAGUCAGUGUGGAAAAUCUCUAGCAGUGGCGCCCGAACAGGGAC
CUGAAAGCGAAAGGGAAACCAGAGGAGCUCUCUCGACGCAGGACUCGGCUUGCUGAAGCG
CGCACGGCAAGAGGCGAGGGGCGGCGACUGGUGAGUACGCCAAAAAUUUUGACUAGCGGA
GGCUAGAAGGAGAGAGAUGGGUGCGAGAGCGUCAGUAUUAAGCGGGGGAGAAUUAGAUCG
AUGGGAAAAAAUUCGGUUAAGGCCAGGGGGAAAGAAAAAAUAUAAAUUAAAACAUAUAGUAU
GGGCAAGCAGGGAGCUAGAACGAUUCGCAGUUAAUCCUGGCCUGUUAGAAACAUCAGAAG
GCUGUAGACAAAUACUGGGACAGCUACAACCAUCCCUUCAGACAGGAUCAGAAGAACUUAG
AUCAUUAUAUAAUACAGUAGCAACCCUCUAUUGUGUGCAUCAAAGGAUAGAGAUAAAAGACA
CCAAGGAAGCUUUAGACAAGAUAGAGGAAGAGCAAAACAAAAGUAAGAAAAAAGCACAGCAA
GCAGCAGCUGACACAGGACACAGCAAUCAGGUCAGCCAAAAUUACCCUAUAGUGCAGAACA
UCCAGGGGCAAAUGGUACAUCAGGCCAUAUCACCUAGAACUUUAAAUGCAUGGGUAAAAGU
AGUAGAAGAGAAGGCUUUCAGCCCAGAAGUGAUACCCAUGUUUUCAGCAUUAUCAGAAGGA
GCCACCCCACAAGAUUUAAACACCAUGCUAAACACAGUGGGGGGACAUCAAGCAGCCAUGC
AAAUGUUAAAAGAGACCAUCAAUGAGGAAGCUGCAGAAUGGGAUAGAGUGCAUCCAGUGCA
UGCAGGGCCUAUUGCACCAGGCCAGAUGAGAGAACCAAGGGGAAGUGACAUAGCAGGAAC
UACUAGUACCCUUCAGGAACAAAUAGGAUGGAUGACAAAUAAUCCACCUAUCCCAGUAGGA
GAAAUUUAUAAAAGAUGGAUAAUCCUGGGAUUAAAUAAAAUAGUAAGAAUGUAUAGCCCUAC
CAGCAUUCUGGACAUAAGACAAGGACCAAAGGAACCCUUUAGAGACUAUGUAGACCGGUUC
UAUAAAACUCUAAGAGCCGAGCAAGCUUCACAGGAGGUAAAAAAUUGGAUGACAGAAACCU
UGUUGGUCCAAAAUGCGAACCCAGAUUGUAAGACUAUUUUAAAAGCAUUGGACCAGCGGC
```

```
UACACUAGAAGAAAUGAUGACAGCAUGUCAGGGAGUAGGAGGACCCGGCCAUAAGGCAAGA
GUUUUGGCUGAAGCAAUGAGCCAAGUAACAAAUUCAGCUACCAUAAUGAUGCAGAGAGGCA
AUUUUAGGAACCAAAGAAAGAUUGUUAAGUGUUUCAAUUGUGGCAAAGAAGGGCACACAGC
CAGAAAUUGCAGGGCCCCUAGGAAAAAGGGCUGUUGGAAAUGUGGAAAGGAAGGACACCAA
AUGAAAGAUUGUACUGAGAGACAGGCUAAUUUUUUAGGGAAGAUCUGGCCUUCCUACAAGG
GAAGGCCAGGGAAUUUUCUUCAGAGCAGACCAGAGCCAACAGCCCCACCAGAAGAGAGCUU
CAGGUCUGGGGUAGAGACAACAACUCCCCCUCAGAAGCAGGAGCCGAUAGACAAGGAACUG
UAUCCUUUAACUUCCCUCA
```

##### Study Material 3

Material 3 is composed of total RNA extracted from cultured HIV-1 and corresponds to the same strain used to prepare the WHO International Standard for HIV-1,

KJ019215.1 HIV-1 isolate NIBSC-1 from United Kingdom (1).

#### 2. HIV-1 *gag* gene Sequence information

Sequences are provided in fasta format obtained from https://www.ncbi.nlm.nih.gov/ for digital PCR assay design in relation to the analysis of Study Materials 1 and 3 (Study Material 2 is intended for analysis with orthogonal methods).

##### Study Material 1

HIV-1 *gag* gene sequence derived from K03455.1 Human immunodeficiency virus type 1 (HXB2) HIV1/HTLV-III/LAV reference genome (positions 790 to 2,257).

```
>K03455.1: 790-2257 Human immunodeficiency virus type 1 (HXB2), complete genome; HIV1/HTLV-III/LAV reference genome
AUGGGUGCGAGAGCGUCAGUAUUAAGCGGGGGAGAAUUAGAUCGAUGGGAAAAAAUUCGG
UUAAGGCCAGGGGGAAAGAAAAAAUAUAAAUUAAAACAUAUAGUAUGGGCAAGCAGGGAGC
UAGAACGAUUCGCAGUUAAUCCUGGCCUGUUAGAAACAUCAGAAGGCUGUAGACAAAUACU
GGGACAGCUACAACCAUCCCUUCAGACAGGAUCAGAAGAACUUAGAUCAUUAUAUAAUACA
GUAGCAACCCUCUAUUGUGUGCAUCAAAGGAUAGAGAUAAAAGACACCAAGGAAGCUUUAG
ACAAGAUAGAGGAAGAGCAAAACAAAAGUAAGAAAAAAGCACAGCAAGCAGCAGCUGACACA
GGACACAGCAAUCAGGUCAGCCAAAAUUACCCUAUAGUGCAGAACAUCCAGGGGCAAAUGG
UACAUCAGGCCAUAUCACCUAGAACUUUAAAUGCAUGGGUAAAAGUAGUAGAAGAGAAGGC
UUUCAGCCCAGAAGUGAUACCCAUGUUUUCAGCAUUAUCAGAAGGAGCCACCCCACAAGAU
UUAAACACCAUGCUAAACACAGUGGGGGGACAUCAAGCAGCCAUGCAAAUGUUAAAAGAGA
CCAUCAAUGAGGAAGCUGCAGAAUGGGAUAGAGUGCAUCCAGUGCAUGCAGGGCCUAUUG
CACCAGGCCAGAUGAGAGAACCAAGGGGAAGUGACAUAGCAGGAACUACUAGUACCCUUCA
GGAACAAAUAGGAUGGAUGACAAAUAAUCCACCUAUCCCAGUAGGAGAAAUUUAUAAAAGAU
GGAUAAUCCUGGGAUUAAAUAAAAUAGUAAGAAUGUAUAGCCCUACCAGCAUUCUGGACAU
AAGACAAGGACCAAAGGAACCCUUUAGAGACUAUGUAGACCGGUUCUAUAAAACUCUAAGA
GCCGAGCAAGCUUCACAGGAGGUAAAAAAUUGGAUGACAGAAACCUUGUUGGUCCAAAAUG
CGAACCCAGAUUGUAAGACUAUUUUAAAAGCAUUGGGACCAGCGGCUACACUAGAAGAAAU
GAUGACAGCAUGUCAGGGAGUAGGAGGACCCGGCCAUAAGGCAAGAGUUUUGGCUGAAGC
```

AAUGAGCCAAGUAACAAAUUCAGCUACCAUAAUGAUGCAGAGAGGCAAUUUUAGGAACCAAA
GAAAGAUUGUUAAGUGUUUCAAUUGUGGCAAAGAAGGGCACACAGCCAGAAAUUGCAGGGC
CCCUAGGAAAAAGGGCUGUUGGAAAUGUGGAAAGGAAGGACACCAAAUGAAAGAUUGUACU
GAGAGACAGGCUAAUUUUUUAGGGAAGAUCUGGCCUUCCUACAAGGGAAGGCCAGGGAAU
UUUCUUCAGAGCAGACCAGAGCCAACAGCCCCACCAGAAGAGAGCUUCAGGUCUGGGGUA
GAGACAACAACUCCCCCUCAGAAGCAGGAGCCGAUAGACAAGGAACUGUAUCCUUUAACUU
CCCUCA

**Study Material 3**

HIV-1 *gag* gene sequence derived from KJ019215.1 HIV-1 isolate NIBSC-1 from United Kingdom (positions 220 to 1,728).

**>KJ019215.1: 220-1728 HIV-1 isolate NIBSC-1 from United Kingdom gag protein (gag), pol protein (pol), vif protein (vif), vpr protein (vpr), tat protein (tat), rev protein (rev), vpu protein (vpu), envelope glycoprotein (env), and nef protein (nef) genes, complete cds**
AUGGGUGCGAGAGCGUCAGUAUUAAGCGGGGGAGCAUUGGAUAGGUGGGAAAGAAUUCGG
UUAAGGCCAGGGGGAAAGAAAAGAUAUCAAUUAAAACAUAUAGUAUGGGCAAGCAGGGAAC
UAGAACGAUUCGCAAUUAAUCCUGGCCUGUUAGAAACGUCAGAAGGCUGUAGACAAAUACU
AGAACAGCUACAAUCAACCCUUAAGACAGGAUCAGAAGAACUUAGAUCAUUAUAUAAUACAG
UAGCAACCCUCUAUUGUGUGCAUCAACAGAUAGAAGUAAAAGACACCAAGGAAGCUUUAGA
CAAGAUAGAGGAAGAGCAAAACAAAAGUAAGAAAAAAGCACAGCAAGCAGCAGCUGGCACAG
GAAACAGCAGUCCGGUCAGCCAAAAUUACCCUAUAGUGCAGAACAUCCAGGGGCAAAUGGU
ACAUCAGGCCAUAUCACCUAGAACUUUAAAUGCAUGGGUAAAAGUAGUAGAAGAGAAGGCU
UUCAGCCCAGAAGUAAUACCCAUGUUUUCAGCAUUAUCAGAAGGAGCCACCCCACAGGAUU
UAAACACCAUGCUAAACACAGUGGGGGGACAUCAAGCAGCCAUGCAAAUGUUAAAAGAGAC
CAUAAAUGAGGAAGCUGCAGAAUGGGAUAGAUUGCAUCCAGUGCAGGCAGGGCCUAUUGC
ACCAGGCCAAAUGAGAGAACCAAGGGGAAGUGACAUAGCAGGAAGUACUAGUACCCUUCAG
GAACAAAUAGGAUGGAUGACAGCUAAUCCAGCUAUCCCAGUAGGAGAAAUCUAUAAAAGAU
GGAUAAUCAUGGGAUUAAAUAAAAUAGUAAGGAUGUAUAGCCCUACCAGCAUUUUGGACAU
AAAACAAGGACCAAAGGAACCCUUUAGGGACUAUGUAGACCGGUUCUAUAAGACUCUAAGA
GCCGAGCAAGCUUCACAGGAAGUAAAAAAUUGGAUGACAGAAACCUUGUUGGUCCAAAAUG
CAAACCCAGAUUGUAAGACUAUUUUAAAAGCAUUGGGACCAGCAGCUACACUAGAAGAAAU
GAUGACAGCAUGUCAGGGAGUGGGGGGACCUGGCCAUAAAGCAAGAGUUUUGGCAGAAGC
AAUGAGCCAGGUAACAAAUUCAGCUGCCAUAAUGAUGCAGAAAGGCAAUUUCAGAAACCAAA
GAAAAACUGUUAAGUGUUUCAAUUGUGGCAAAGAAGGGCACAUAGCCAGAAAUUGCAGGGC
CCCUAGGAAAAAGGGCUGUUGGAAAUGUGGAGAGGAAGGACACCAAAUGAAAGAUUGUACU
GGGAGACAGGCUAAUUUUUUAGGGAAAAUCUGGCCUUCCCACAAGGGAAGGCCAGGGAAU
UUUCUUCAGAGCAGACCAGAGCCAACAGCCCCACCAGCCCCACCAGAAGAGAGCUUCAGGU
UUGGGGAGGAGACAACAACUCCCCCUCAAAAGCAGGAGAAGAUAGACAAGGAACUGUAUCC
UUUAGCUUCCCUCAGAUCACUCUUUGGCAACGACCCCUCGUCACAA

### REFERENCES

1. Gall A, Morris C, Kellam P, Berry N. Complete Genome Sequence of the WHO International Standard for HIV-1 RNA Determined by Deep Sequencing. Genome Announcements. 2014;2(1).
