## Supplementary material for "CCQM-P199: Interlaboratory comparability study of HIV-1 RNA copy number quantification": CCQM P199 supplementary file Appendix B

### APPENDIX B: Coordinating laboratory methodology

#### Coordinating laboratory methodology: NIBSC

##### RNA purification from NIBSC HIV-1 Standard (Study Material 3)

RNA was extracted from 4 units of viral material previously used to produce the WHO 4th HIV-1 International Standard (NIBSC code 16/194; GenBank Accession KJ019215.1 HIV-1 isolate NIBSC-1) using the QIAamp UltraSens Virus Kit (Cat no 53704, Qiagen). Four vials each containing approximately 60 µL of eluate were transported to the NML on dry ice and stored at -80 °C on arrival.

#### Coordinating laboratory methodology: NML

##### Study Materials 1-2: Synthetic construct design

A region of the human immunodeficiency virus type 1 (HXB2) complete HIV1/HTLV-III/LAV reference genome (K03455.1 positions 1 to 5,619) was identified for generation of a synthetic RNA test molecule. A 5,675 bp insert, which included an 18 bp T7 RNA polymerase promoter sequence TAATACGACTCACTATAG (1), was cloned into a standard pEX-A2 vector. Gene insertion in the correct orientation was confirmed by Sanger sequencing (Eurofins genomics).

##### *In vitro* transcription of RNA

One microgram of the HIV-1 HXB2 plasmid was linearised between positions 2,254 and 2,255 (K03455.1) with 0.2 units/µL *Bsu*36I (R0524S), 1× CutSmart buffer (B72045, both New England Biolabs) and nuclease-free water (AM9937), in each of six replicate digests containing a reaction volume of 50 µL, for 1 h at 37 °C. The reaction was terminated by heating at 80 °C for 20 min, and each digest was purified using the QIAquick PCR purification kit (28104, Qiagen) with elution into 50 µL elution buffer. Successful linearisation of an ≈ 8,125 bp molecule was confirmed using the 2100 Bioanalyzer with DNA 7500 series II kit (Agilent) (Figure B-1). A slightly smaller peak of ≈ 7,799 bp was observed which was within 95 % of the expected size. DNA concentration was estimated using the Qubit 2.0 fluorometer with the dsDNA HS Assay Kit (Q32851, Invitrogen). Following initial quality control (QC), the six replicate digests were pooled to give a volume of ≈ 270 µL and the concentration measured again using the dsDNA HS Qubit kit.

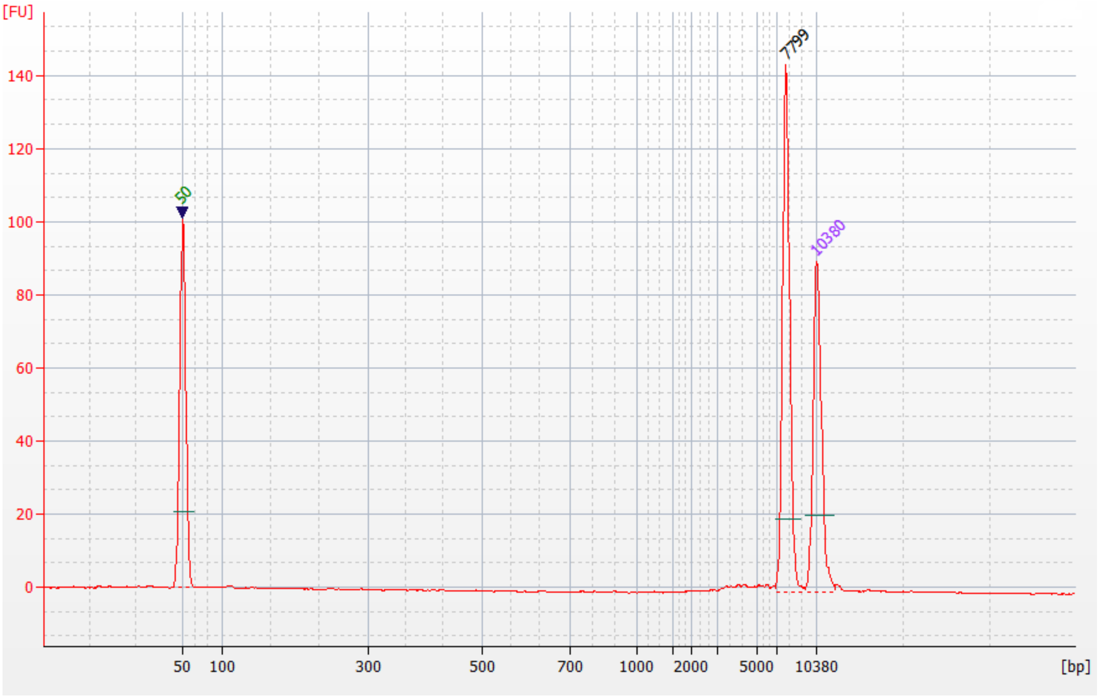


**Figure B-1: Linearised plasmid assessed using the 2100 Bioanalyzer with DNA 7500 series II kit (Agilent), confirming the presence of a ≈ 8,125 bp molecule.**

To generate positive sense strand RNA *in vitro* transcription (IVT) was performed using the MEGAscript T7 kit (AM1334, Life Technologies). Fifteen replicate reactions were included, each containing 7.5 mM of each of ATP, CTP, GTP and UTP, 1X Reaction Buffer, 4 µL T7 enzyme mix and 16 µL (approximately 174 ng) of plasmid. Incubation was performed at 37 °C for 4 h followed by TURBO DNase treatment. The resulting RNA was purified using the RNeasy Mini Kit for RNA clean up protocol (Cat no 74104, Qiagen), which included an additional on-column treatment with RNase-free DNase I (Cat no 79254, Qiagen). RNA transcripts were eluted in 50 µL RNase-free water. A 2.2 µL aliquot of RNA was taken and the nucleic acid concentration estimated using a NanoDrop 2000 spectrophotometer (Thermo Scientific). Successful *in vitro* transcription was confirmed by analysing 1 µL of the neat RNA stock with the 2100 Bioanalyzer RNA 6000 Nano kit (Agilent) and agarose gel electrophoresis (Lonza FlashGel) (Figure B-2). Transcripts were expected to be 2,266 nt in length. The IVT replicates were pooled and diluted to an approximate RNA concentration of 100 ng/µL based on NanoDrop. The concentration of the pooled, diluted RNA (approximately 8.0 mL total volume) was confirmed using a Qubit 2.0 fluorometer and a Qubit RNA BR Assay Kit (Q10210, Invitrogen). Total molecular weight (MW, g/mol) of the single stranded RNA transcript was estimated by multiplying the number of each nucleotide present (A, C, G, U) by the respective MW. Mass per RNA molecule (g) was calculated using the Avogadro number ($6.022 \times 10^{23}$ $mol^{-1}$). Copy number concentration in the stock RNA solution was calculated using the Qubit results and the mass per RNA molecule in g.

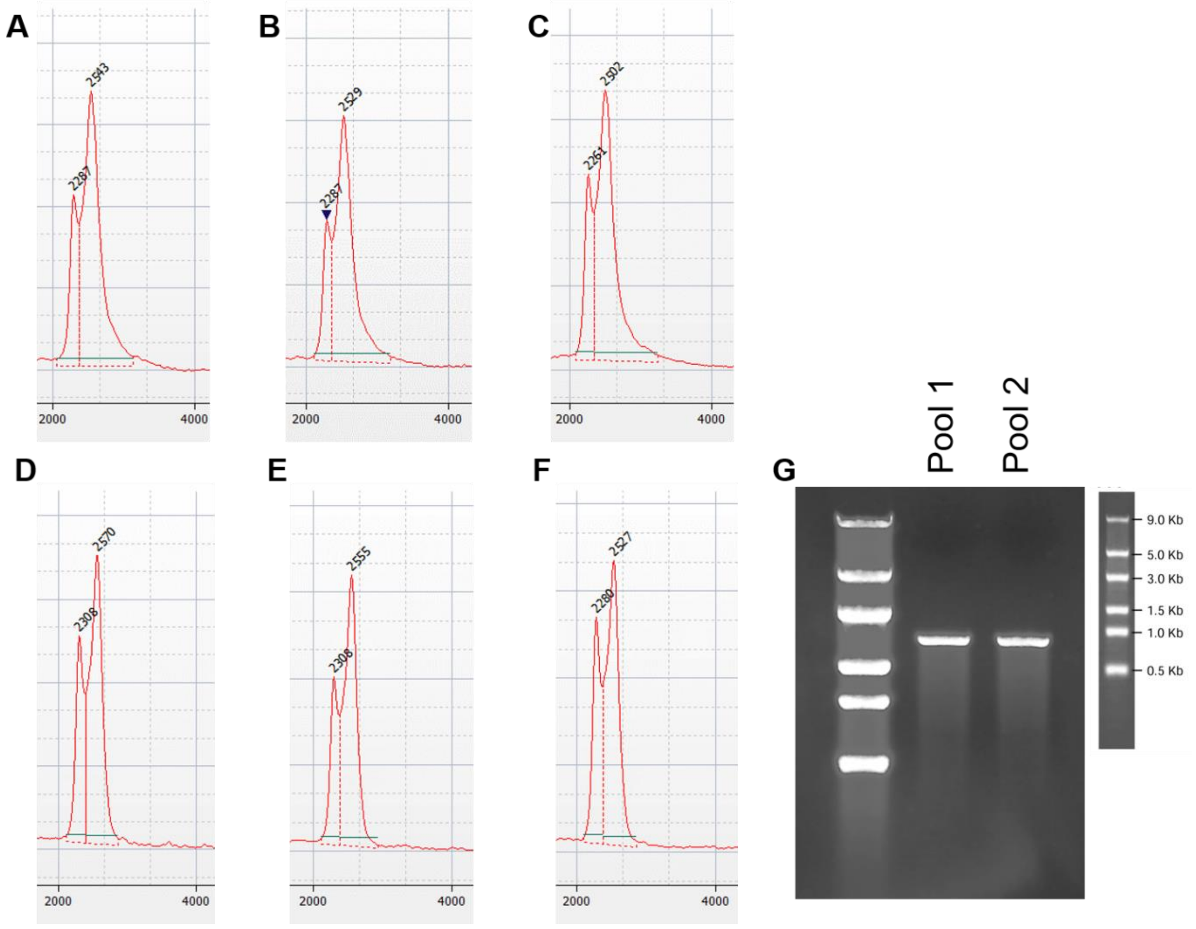


**Figure B-2: *In vitro* transcribed HIV-1 HXB2 RNA fragment size assessed using the 2100 Bioanalyzer RNA 6000 Nano kit (Agilent) and Flashgel (Lonza).**
Replicate Bioanalyzer measurements ($n$ = 3) of pooled IVT product used in the preparation of Study Materials 1 and 2 are shown: Pool 1 (A-C); Pool 2 (D-F) in the size range 2000 nt to 4000 nt, with fragment size (nt) indicated above each peak. (G) Flashgel image of RNA ladder (lane 1, insert) and Pools 1-2 (lanes 2-3).

Subsequent analysis of Bioanalyzer data (Figure B-2) describes the size distribution and AUC for the IVT stock material used for Study Material 1 and 2 production (Table B-1).

**Table B-1: Analysis of Bioanalyzer results for HIV-1 HXB2 *in vitro* transcribed RNA pools**

| | | **Peak 1** | | | | **Peak 2** | | | |
|---|---|---|---|---|---|---|---|---|---|
| IVT Pool | Replicate | **Start Size [nt]** | **End Size [nt]** | **Area (AU)** | **% Peak/total area** | **Start Size [nt]** | **End Size [nt]** | **Area (AU)** | **% Peak/total area** |
| 1 | 1 | 2019 | 2351 | 1.80 | 14 % | 2380 | 3100 | 11.20 | 86 % |
| | 2 | 2088 | 2351 | 2.00 | 16 % | 2380 | 2975 | 10.50 | 84 % |
| | 3 | 2076 | 2324 | 2.50 | 19 % | 2352 | 2959 | 10.90 | 81 % |
| 2 | 1 | 2169 | 2382 | 4.10 | 24 % | 2411 | 2780 | 13.30 | 76 % |
| | 2 | 2169 | 2351 | 2.00 | 13 % | 2380 | 2752 | 13.00 | 87 % |
| | 3 | 2142 | 2354 | 4.70 | 29 % | 2383 | 2730 | 11.50 | 71 % |
| | **Mean** | **2111** | **2352** | | **19 %** | **2381** | **2883** | | **81 %** |
| | SD | 60 | 18 | | 6 % | 19 | 150 | | 6 % |

### RT-dPCR: Oligonucleotide sequences

**Table B-2: Primer and probe sequences homologous to the HIV-1 HXB2 reference genome (K03455.1) and HIV-1 isolate NIBSC-1 (KJ019215.1) used for analysis and development of study materials**

| Assay designation | Genbank accession/ loci | Name | 5' to 3' * | Amplicon (bp) | Source |
|---|---|---|---|---|---|
| HIV-1 *gag* | K03455.1: 1252-1367<br><br>KJ019215.1: 682-797 | HIV-1 *gag* P | FAM-TCAGCATTATCAGAAGGAG-MGBEQ | 116 | (2) |
| | | HIV-1 *gag* F | TGGGTAAAAGTAGTAGAAGAGAAGGCTTT | | |
| | | HIV-1 *gag* R | CCCCCCACTGTGTTTAGCAT | | |
| Long Terminal Repeat – *gag* junction of HIV-1 (HIV-1 LTR-*gag*) | K03455.1: 522-642 | HIV_FAMv2 | FAM-TCTGAGGGA-Nova-TCTCTAGTTACCAGAGTCACA-BHQ | 121 | (3), probe this study |
| | | HIV1LTR_F | GCCTCAATAAAGCTTGCCTTGA | | |
| | | HIV1LTR_R | GGCGCCACTGCTAGAGATTTT | | |
| HIV-1 *pol* | KJ019215.1: 1975-2101 | *Pol* P | FAM-AAGCCAGGAATGGATGGCC-NFQMGB | 127 | (4) |
| | | *Pol* F | GCACTTTAAATTTTCCCATTAGTCCTA | | |
| | | *Pol* R | CAAATTTCTACTAATGCTTTTATTTTTTC | | |

### One-step RT-dPCR methodology

One-step RT-dPCR experiments were performed using the One-Step RT-ddPCR Advanced Kit for Probes (Cat no. 1864021, Bio-Rad). Reactions were prepared in a total volume of 22 µL containing 1X Supermix, 20 U/µL reverse transcriptase, 15 mM[1] DTT, 5.5 µL of RNA template, nuclease-free water and primers and probes at a concentration of 900 nM and 200 nM (50 nM of the HIV-1 *pol* probe), respectively. RNA templates were heat denatured at 65 °C for 5 min and immediately quenched on ice prior to addition into the reaction. dPCR was performed using the QX200™ Droplet Digital™ PCR System (Bio-Rad). 20 µL was pipetted into the sample well of a DG8 cartridge, and droplets generated as previously described (5). Thermocycling conditions were as follows: Reverse transcription at 47.5 °C for 60 min, 10 min at 95 °C, 40 cycles of 94 °C for 30 s, and 58 °C for 1 min, followed by 98 °C for 10 min and a 4 °C hold. The ramp rate for each step was 2 °C/s. Droplets were read using the QX200 Droplet Reader, and the data were analyzed using QuantaSoft version 1.7.4.0917. No Template Controls (NTCs) of nuclease-free water were employed as controls, along with RT negative controls (water in place of reverse transcriptase), and in all cases returned a negative result. A partition volume of 0.776 nL was used to calculate copy number concentration for preparation

[1] The molecular biology community express amount of substance concentration with units of molarity (M) (SI units: mmol/L).

of the study materials. Data from dPCR experiments were subject to threshold and baseline setting in QuantaSoft software (Bio-Rad), and were exported as .csv files to be analysed in Microsoft Excel 2010. The average number of copies per droplet (λ) was calculated as described previously (6).

### Gravimetry

Gravimetric preparation of the Study Materials was performed using a Mettler Toledo XP205 balance to 5 decimal places. Following cleaning of the balance, linearity was tested using a set of laboratory standard weights covering the range 0.1 g to 200 g. Standard uncertainty of measurement for the balance was ± 0.000159 g.

### Study Material 1 and 2 preparation

Study Material 2 (SM2) was prepared using the Qubit RNA BR Assay result as an initial indicator of copy number concentration of the prepared HXB2 RNA transcript stock. A target dilution factor (DF) of 3.25 was chosen to prepare SM2 from the stock. The pooled, diluted stock of HXB2 RNA transcript was thawed and diluted gravimetrically in RNA Storage Solution (Table B-3).

**Table B-3: Gravimetric preparation of Study Material 2.**

| Mass of tube (g) | Diluent vol. added (mL) | Mass of tube + diluent (g) | Mass of diluent (g) | Sample vol. added (mL) | Mass of tube + diluent + sample (g) | Mass of sample (g) | Volumetric DF | Gravimetric DF |
|---|---|---|---|---|---|---|---|---|
| 14.4768 | 17.1280 | 31.5671 | 17.0903 | 7.6220 | 39.1840 | 7.6169 | 3.2471 | 3.2437 |

The dilution factor to prepare SM2 from the stock solution was 3.24 (± 0.00010) based on gravimetry. The prepared solution of SM2 was placed on a roller mixer at 4 °C for 30 min to obtain a homogenous solution prior to aliquoting the units. A total of 120 units were prepared using a E3 multipipette (Eppendorf) to a volume of 200 µL. Filling accuracy of the pipette was checked prior to aliquoting of the units using RNA Storage Solution (Table B-4).

**Table B-4: Record of filling accuracy using 200 µL (0.2 g) of RNA Storage Solution (RSS).**

| Measurement | Weight of tube (g) | Weight after filling with 200 µL RSS (g) | Added (g) | Vol according to mass (µL) |
|---|---|---|---|---|
| 1 | 1.5016 | 1.7004 | 0.1988 | 198.8 |
| 2 | 1.5028 | 1.7030 | 0.2002 | 200.2 |
| 3 | 1.5130 | 1.7133 | 0.2003 | 200.3 |
| 4 | 1.5073 | 1.7073 | 0.2000 | 200.0 |
| 5 | 1.5109 | 1.7099 | 0.1990 | 199.0 |
| 6 | 1.5127 | 1.7134 | 0.2007 | 200.7 |
| 7 | 1.5069 | 1.7067 | 0.1998 | 199.8 |
| 8 | 1.5055 | 1.7058 | 0.2003 | 200.3 |
| 9 | 1.5069 | 1.7077 | 0.2008 | 200.8 |
| 10 | 1.5053 | 1.7052 | 0.1999 | 199.9 |
| | | Mean | 0.2000 | |
| | | SD | 0.0007 | |

Following gravimetric preparation of SM2, ten 55 µL aliquots were retained for RT-dPCR analysis, and for preparation of Study Material 1 (SM1). For three separate aliquots, volumetric dilutions were prepared to an input concentration of 1.09 x$10^4$ /µL (based on the Qubit result) and analysed by one-step RT-dPCR using the HIV-1 *gag* assay (Table B-2). SM1 was gravimetrically prepared based on the one-step RT-dPCR result using a thawed 55 µL aliquot of SM2 (termed 'Mat 2'). A target dilution factor (DF) of 4.81 x$10^6$ was planned to prepare SM1 from SM2, using Human Jurkat cell total RNA as carrier (Ambion AM7858). Carrier solution was gravimetrically prepared to a concentration of 4.8 ng/µL in RNA Storage Solution using the same 5-figure balance as used to prepare SM2 and SM1.

**Table B-5: Gravimetric preparation of Study Material 1.**

| Sample | Mass of tube (g) | Sample (µL) | Mass of tube + sample (g) | Mass of sample (g) | Diluent vol. (µL) | Mass of tube + sample + diluent (g) | Mass of diluent (g) | Volumetric DF | Gravimetric DF |
|---|---|---|---|---|---|---|---|---|---|
| D1 | 0.9966 | 50.0 | 1.0453 | 0.0487 | 450.0 | 1.4963 | 0.4510 | 10.0 | 10.3 |
| D2 | 0.9955 | 50.0 | 1.0438 | 0.0483 | 950.0 | 1.9861 | 0.9423 | 20.0 | 20.5 |
| D3 | 1.0080 | 50.0 | 1.0579 | 0.0499 | 950.0 | 2.0095 | 0.9516 | 20.0 | 20.1 |
| D4 | 1.0970 | 75.0 | 1.1706 | 0.0736 | 1425.0 | 2.5961 | 1.4255 | 20.0 | 20.4 |
| D5 | 1.0908 | 583.3 | 1.6787 | 0.5879 | 1166.7 | 2.8335 | 1.1548 | 3.0 | 3.0 |
| SM1 | 14.4770 | 1645.7 | 16.1113 | 1.6343 | 31354.3 | 47.4180 | 31.3067 | 20.1 | 20.2 |

Based on gravimetry, there was a dilution factor of $5.14 \times 10^6$ ($\pm 4.40 \times 10^4$) between SM2 and SM1 (Table B-5). The prepared solution of SM1 was placed on a roller mixer at 4 °C for a total of 30 min to obtain a homogenous solution prior to aliquoting the units. A total of 327 units were prepared using a E3 multipipette (Eppendorf) to a volume of 100 µL. Filling accuracy of the pipette was checked prior to aliquoting of the units using RNA Storage Solution (Table B-6).

**Table B-6: Record of filling accuracy using 100 µL (0.1 g) of RNA Storage Solution (RSS).**

| Measurement | Weight of tube (g) | Weight after filling with 100 µL RSS (g) | Added (g) | Vol according to mass (µL) |
|---|---|---|---|---|
| 1 | 1.50391 | 1.60327 | 0.0994 | 99.4 |
| 2 | 1.52252 | 1.62212 | 0.0996 | 99.6 |
| 3 | 1.51383 | 1.61412 | 0.1003 | 100.3 |
| 4 | 1.52253 | 1.62261 | 0.1001 | 100.1 |
| 5 | 1.52036 | 1.62015 | 0.0998 | 99.8 |
| 6 | 1.50612 | 1.60638 | 0.1003 | 100.3 |
| 7 | 1.50809 | 1.60808 | 0.1000 | 100.0 |
| 8 | 1.50227 | 1.60211 | 0.0998 | 99.8 |
| 9 | 1.51036 | 1.61099 | 0.1006 | 100.6 |
| 10 | 1.52196 | 1.62265 | 0.1007 | 100.7 |
| | | Mean | 0.1001 | |
| | | SD | 0.0004 | |

### Study Material 3 preparation

Purified RNA from HIV-1 (NIBSC code 16/194) received from NIBSC was thawed and the four vials were pooled (approximately 212.5 µL total volume). A 20 µL aliquot was removed for analysis, and the remaining ≈ 192.5 µL of RNA stock solution returned to storage at -80 °C.

An initial one-step RT-dPCR experiment was performed using the HIV-1 *gag* assay (Table B-2) to determine the copy number concentration of the purified RNA. The HIV-1 RNA stock was volumetrically diluted to approximately $2.0 \times 10^3$ /µL to provide enough volume for Study Material preparation, and the one-step RT-dPCR analysis was repeated. Based on the result, Study Material 3 (SM3) was gravimetrically prepared in Human Jurkat cell total RNA carrier (Ambion AM7858), which had been gravimetrically prepared to a concentration of 3.1 ng/µL in RNA Storage Solution. A target dilution factor (DF) of 10.6 was planned to prepare the material from the diluted RNA (≈ $2.0 \times 10^3$ /µL).

**Table B-7: Gravimetric preparation of Study Material 3.**

| Sample | Mass of tube (g) | Diluent vol. added (mL) | Mass of tube + diluent (g) | Mass of diluent (g) | Sample vol. added (mL) | Mass of tube + diluent + sample (g) | Mass of sample (g) | Volumetric DF | Gravimetric DF |
|---|---|---|---|---|---|---|---|---|---|
| SM3 | 14.46489 | 29.891 | 44.30500 | 29.840 | 3.109 | 47.40656 | 3.102 | 10.6 | 10.621 |

Based on gravimetry, a dilution factor of 10.62 (± 0.00077) was applied to prepare SM3 (Table B-7). The prepared solution of SM3 was placed on a roller mixer at 4 °C for a total of 30 min to obtain a homogenous solution prior to aliquoting the units. A total of 325 units were prepared using a E3 multipipette (Eppendorf) to a volume of 100 µL. All materials were stored at -80 °C.

### Homogeneity study

Homogeneity of Study Materials (SM) 1 and 3 was assessed by one-step RT-dPCR using the HIV-1 *gag* assay (Table B-2). Ten undiluted units of each of SM1 and SM3 were assessed, and 8 RT-dPCR replicates were analysed per unit.

For homogeneity testing of Study Material 2 (SM2), ten undiluted units were analysed using a Quantus Fluorometer (Cat: E6150, Promega) and QuantiFluor RNA Kit (Cat: E3310, Promega). All assay components, which included 20× TE Buffer (pH 7.5), Quantifluor RNA dye and RNA standard (100 µg/mL), were equilibrated to ambient temperature. A working solution was prepared following the 'high standard calibration' protocol by combining the RNA dye with the TE buffer diluted to 1X in nuclease-free water (AM9937). The 'High Standard Calibration' Quantus programme was selected and a standard curve calibration performed using a blank sample (working solution only) and the high RNA standard, which was prepared by adding 5 µL (500 ng) of RNA standard to 200 µL of working solution. Eight replicate assay tubes were prepared for each of the ten units of SM2 by adding 5 µL of sample to 200 µL of working solution, along with eight blank assay tubes containing RNA Storage Solution in working solution. The assay tubes (n =88) were analysed following a randomized order to determine RNA concentration of SM2 in ng/µL.

***Statistical methods:*** Data for one measurement was removed from the Study Material 1 homogeneity dataset (Unit 7, Replicate # 6, well F1) due to being a statistical outlier. Homogeneity data for Study Materials 1, 2 and 3 were analysed using R version 3.6.1 running inside RStudio version 1.2.5001, using mixed effects models with maximum likelihood estimation. Models with mean value and dPCR plate row and column as fixed effects, and between unit variation as (nested) random effect(s) were evaluated, with the most appropriate model chosen based on Akaike Information Criterion (AIC). For Material 2 the best model was with mean value as the only fixed effect. Replicate order was found to be significantly associated with concentration (p value <0.001).

### Short-term stability study

Short-term stability (STS) of Study Materials 1, 2 and 3 was assessed isochronously following incubation on dry ice and at 27 °C for 1 day and 7 days in comparison to a reference temperature of -80 °C. Three units of each material were included per condition. For SM1 and SM3 short-term stability was assessed by one-step RT-dPCR using the HIV-1 *gag* assay (Table B-2). Short-term stability of SM2 was assessed using a QuantiFluor RNA Kit (Cat: E3310, Promega) and the High Standard Calibration protocol as described for the homogeneity study. Three assay tubes were analysed for each of the 18 units of SM2 (3 units per storage condition). In addition to fluorometric analysis, the 18 undiluted STS units of SM2 were analysed on the Agilent Bioanalyser 2100 using an RNA 6000 Nano kit, according to the manufacturer's instructions.

***Statistical methods:*** One data point (Study Material 1, dry ice condition, unit "S1", RT-dPCR replicate #3) was excluded from the final data analysis due to be a statistical outlier.

### Long-term stability study

Long-term stability (LTS) of the materials was assessed at two time points; 4 to 5 months and 7 to 9 months after the homogeneity and short-term stability studies. Prior to LTS analysis, four units of each of SM1, SM2 and SM3 were incubated on dry ice for 7 days to simulate conditions during shipping. These were compared to four units of each material that had been stored at the reference temperature of –80 °C without dry ice incubation. The undiluted units of SM1 and SM3 were analysed by one-step RT-dPCR using the HIV-1 *gag* assay (Table B-2). For LTS testing of SM2, a volumetric dilution series was prepared for each of the eight units (four with and four without dry ice incubation) to a concentration of 5,000 /µL based on RT-dPCR analysis. Yeast tRNA (AM7119, ThermoFisher Scientific) was used as diluent at a concentration of 5 ng/µL. Long-term stability was assessed by one-step RT-dPCR using the HIV-1 *gag* assay (Table B-2). In addition, the undiluted units were analysed using the QuantiFluor RNA Kit as described for the short-term stability (STS) and homogeneity studies.

### Intermediate precision of the QuantiFluor® RNA system

The intermediate precision of the QuantiFluor® RNA System using the Promega Quantus™ Fluorometer was evaluated. A dilution series of the Qubit™ XR Assay RNA Standard (Invitrogen, Q33236) consisting of 4 concentrations (over a range of 0.4 ng/µL to 50 ng/µL) was constructed in RNA Storage Solution (Ambion, AM7001). Six replicates of each dilution were measured on three different days, RNA Storage Solution (RSS) controls were also included as blank samples. 5 µL of each sample was analysed using the High (1 ng/µL to 500 ng/µL) and Low (0.1 ng/µL to 10 ng/µL) standard calibration protocols, as described for Homogeneity, STS and LTS studies.

### Inter-platform analysis using the Stilla® Naica® system

Study Materials 1 and 3 were analysed using the HIV-1 *gag* assay (Table B-2) on the Naica® digital PCR system (Naica). One unit of each study material was analysed in quadruplicate. RNA templates were heat denatured at 65 °C for 5 min and immediately quenched on ice prior to addition into the reaction. 5.5 µL of template was added to a total prepared reaction volume of 27.5 µL containing qScriptTM XLT One-Step RT-qPCR ToughMix (1X final concentration; Quantabio), sterile nuclease-free water (Ambion), 0.1 µM fluorescein (VWR), and primers and probe at a concentration of 900 nM and 200 nM respectively. Fluorescein solution was prepared by weighing out fluorescein sodium salt (VWR) and solubilising in nuclease-free water (Ambion, USA) to a stock concentration of 200 µM. A working stock of 2 µM fluorescein was prepared in nuclease-free water for use as a reference dye. 25 µL of total reaction volume was applied to a Stilla Sapphire chip (version 2) which was loaded into the Naica Geode. A no template control (NTC) of nuclease-free water was also included in the experiment. Partitioning was achieved under 950 mbar of pressure at 40 °C for 12 min (7). Reverse transcription was performed at 50 °C for 10 min, followed by one cycle at 95 °C for 1 min. PCR conditions were 45 cycles of 95 °C for 30 ss and 58 °C for 30 ss. A final decompression step back to atmospheric pressure and room temperature was performed for 33 min. Chips were scanned using the Naica Prism and Crystal Reader software version 2.1.6. The following 133 parameters were applied; focus 0.9 mm, exposure time for blue channel 45 ms, green channel 250 ms, red channel 50 ms. Data were analysed using Crystal miner software version 2.1.6. A partition volume of 0.59 nL (Stilla) was used to calculate copy number concentration (Table B-8).

**Table B-8: Analysis of Study Materials using the Stilla® Naica® system**
Coordinators assigned values and uncertainties for Study Materials 1 and 3 can be found in Table 5 (main text).

| Study Material | Mean *gag* gene concentration (/µL) | SD (/µL) |
|---|---|---|
| SM1 | 957 | 33 |
| SM3 | 166 | 9 |

### Evaluating the impact of multiple thaw-freeze cycles on study material stability

The impact of repeated thawing from storage at -80 °C on material stability was evaluated for Study Materials 1 and 3. Two units of each material were thawed on each of three different days. A 25 µL aliquot was removed for analysis on each day and the units returned to storage at -80 °C. The aliquots of repeatedly thawed material were compared with units that had been stored at -80 °C, with a single thaw just prior to analysis. Aliquots of material that had been exposed to each condition were analysed by one-step RT-dPCR on the QX200 in triplicate, using the HIV-1 *gag* assay (Table B-2). Results are shown in Table B-9. Figure B-3A demonstrates Study Material 1 to be stable following multiple thaw cycles, with no significant effects observed for either unit by one-way ANOVA ($p > 0.05$). The same is observed for Study Material 3 unit 1, however Figure B-3B indicates a possible inverse relationship between RNA copy number concentration and timepoint for unit 2, which was shown to be statistically significant by one-way ANOVA ($p = 0.04$). The study was performed on a limited number of units. Although no significant inter-unit inhomogeneity was demonstrated (Table 3, main text), analysis of additional units would further clarify the impact of multiple thaw cycles on copy number concentration across the batch of prepared material.

Table B-9: Freeze-thaw stability analysis of Study Materials 1 and 3.

Timepoint refers to the number of additional thaw cycles that were encountered prior to analysis.

| **Study Material** | **Unit no.** | **Timepoint** | **Mean *gag* gene concentration (/µL)** | **SD (/µL)** |
|---|---|---|---|---|
| SM1 | 1 | 1 | 979 | 41 |
| | 1 | 2 | 992 | 4 |
| | 1 | 3 | 954 | 17 |
| | 2 | 1 | 1013 | 32 |
| | 2 | 2 | 972 | 35 |
| | 2 | 3 | 986 | 24 |
| | 3 | 0 | 1004 | 38 |
| | 4 | 0 | 968 | 19 |
| SM3 | 1 | 1 | 176 | 12 |
| | 1 | 2 | 179 | 6 |
| | 1 | 3 | 172 | 7 |
| | 2 | 1 | 198 | 19 |
| | 2 | 2 | 168 | 4 |
| | 2 | 3 | 165 | 10 |
| | 3 | 0 | 188 | 8 |
| | 4 | 0 | 189 | 5 |

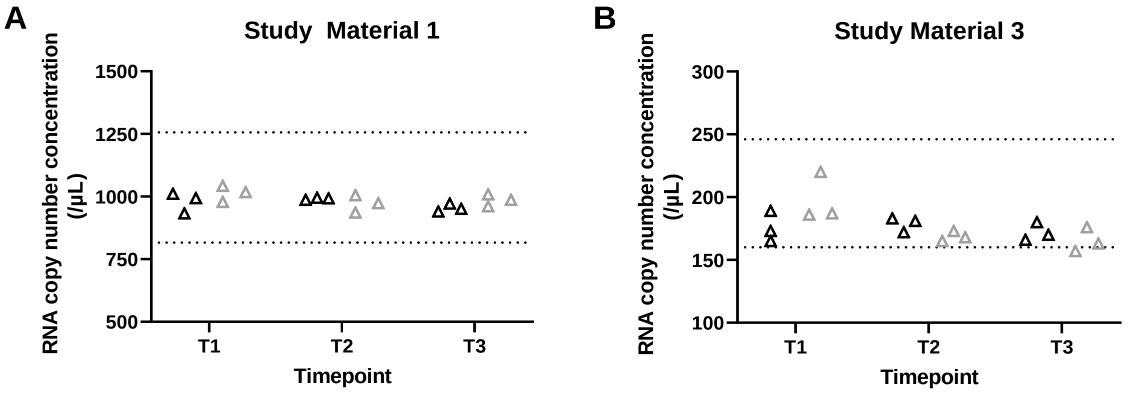


**Figure B-3: Stability of Study Material 1 (A) and Study Material 3 (B) after multiple thaw cycles.**

RNA copy number concentration is plotted for each measurement of two units analysed (black triangles; unit 1, grey triangles: unit 2). Dashed lines show upper and lower limits of coordinator's assigned values (Table 5, main text).

### REFERENCES


1. New England BioLabs. T7 RNA Polymerase Promoter Sequence 2021 [Available from: https://www.neb.sg/faqs/2015/01/30/what-is-the-promoter-sequence-of-t7-rna-polymerase.
2. Bosman KJ, Nijhuis M, van Ham PM, Wensing AM, Vervisch K, Vandekerckhove L, De Spiegelaere W. Comparison of digital PCR platforms and semi-nested qPCR as a tool to determine the size of the HIV reservoir. Scientific Reports. 2015;5:13811.
3. Busby E, Whale AS, Ferns RB, Grant PR, Morley G, Campbell J, Foy CA, Nastouli E, Huggett JF, Garson JA. Instability of 8E5 calibration standard revealed by digital PCR risks inaccurate quantification of HIV DNA in clinical samples by qPCR. Scientific Reports. 2017;7(1):1209.
4. Strain MC, Lada SM, Luong T, Rought SE, Gianella S, Terry VH, Spina CA, Woelk CH, Richman DD. Highly Precise Measurement of HIV DNA by Droplet Digital PCR. PloS one. 2013;8(4):e55943.
5. Devonshire AS, Honeyborne I, Gutteridge A, Whale AS, Nixon G, Wilson P, Jones G, McHugh TD, Foy CA, Huggett JF. Highly reproducible absolute quantification of Mycobacterium tuberculosis complex by digital PCR. Anal Chem. 2015;87(7):3706-13.
6. Whale AS, Bushell C, Grant PR, Cowen S, Guttierrez-Aguirre I, O'Sullivan DM, Zel J, Milavec M, Foy CA, Nastouli E, *et al.* Detection of rare drug resistance mutations by digital PCR in a human influenza A virus model system and clinical samples. J Clin Microbiol. 2016;54(2):392-400.
7. Madic J, Zocevic A, Senlis V, Fradet E, Andre B, Muller S, Dangla R, Droniou ME. Three-color crystal digital PCR. Biomol Detect Quantif. 2016;10:34-46.
