## Supplementary material for "CCQM-P199: Interlaboratory comparability study of HIV-1 RNA copy number quantification": CCQM P199 supplementary file Appendix C

#### APPENDIX C: Protocol

1. **CCQM P199 Study Protocol**
2. **CCQM P199 Protocol Appendix A**

### STUDY PROPOSAL AND PROTOCOL

### CCQM NAWG P199: HIV-1 RNA copy number quantification

#### 1. Aim

The study rationale is to support higher order measurement of RNA copy number concentration of a specific sequence present in the Human Immunodeficiency Virus-1 (HIV-1) viral genome. The specific aims are:

1. To measure the copy number concentration of the HIV-1 *gag* gene.
2. To evaluate variability in results reported between laboratories measuring <10,000 copies of *gag* (Study Materials 1 and 3)
3. To compare the methods used for Study Materials 1 and 3 with orthogonal methods (such as mass-spectrometry or flow cytometric single molecule counting) measuring the same template prepared at two concentrations.
4. To compare *gag* measurements from more simple and complex RNA templates (*in vitro* transcribed RNA and viral genomic RNA)
5. To provide evidence for Calibration and Measurement Claims by participating laboratories.

#### 2. Background

This study is proposed to apply the aims and approach of study "CCQM-P154 Absolute Quantification of DNA" to absolute quantification of RNA.

The aim of CCQM-P154 was to assess the quantification of low-level amounts of DNA in an absolute manner without the aid of calibration using enumeration-based techniques (eight laboratories performed dPCR with one laboratory using flow cytometric single molecule counting). The results reported for the low level 'Level II' material (consensus value 7970 copies/mg) were compared to values reported by laboratories using orthogonal methods (ID-MS, UV-CE) for the approximately 100,000 times-more concentrated 'Level I' material from which the Level II material was prepared. By calculating the gravimetric dilution factors between the Level I and II materials, a direct comparison of the newer methods was possible with the established SI traceable methods used to measure the Level I material (Yoo *et al*., 2016). The close agreement between the mean results of the four orthogonal approaches tested (CV 1.8%) strongly supported the accuracy of more recently developed enumeration-based techniques.

Previous CCQM pilot studies have tested NMI's capabilities to perform accurate measurements of RNA copy number concentration (P103 "Measurement of Multiplexed Biomarker Panel of RNA Transcripts") and RNA copy number ratio (P103.1: "Multiple cancer cell biomarker measurement"). reverse transcription-digital PCR (RT-dPCR) was used by sub-set of laboratories in these studies. Reported values for copy number concentration using RT-dPCR were within 1.5-fold (P103) and 1.4-fold (P103.1) of those assigned on the basis of molecule weight and UV spectroscopy. In CCQM P155 "Multiple cancer cell biomarker measurement", RT-dPCR was used to value assign the copy number concentration of three *in vitro* transcribed transcripts in the study calibration material which enable reporting of mRNA copy number concentration for the two cellular study materials.

Published studies have demonstrated the utility of 'absolute quantification' of mRNA and viral RNA in a number of applications using methods like RT-dPCR (Chen *et al*., 2013; Whale *et al.,* 2013;

White *et al.,* 2012). However error in reverse transcription, currently a necessary step for RNA measurement using molecular biology methodologies, has also been shown to significantly impact dPCR measurements of mRNA (Sanders *et al.,* 2013) and miRNA (Stein *et al.,* 2017).

HIV-1 was chosen as a model for CCQM P199 as it is major pathogen which is quantified clinically to guide treatment and monitor resistance, and is reported as an 'absolute' concentration (such as copies/mL plasma). This differs from other RNA targets measured clinically, which are reported as a copy number ratio, such as the transcript of the *BCR-ABL* fusion gene.

### 3. Study Materials

#### 3.1 Description

Three Study Materials have been prepared by The National Measurement Laboratory at LGC (NML at LGC) and NIBSC (Study Material 3). Sequence information is provided in separate document "CCQM NAWG P199: "HIV-1 RNA copy number quantification": Study Material Sequence information" (14 January 2019). It is intended that all study participants analyse Study Material 1, whereas analysis of Study Materials 2 and 3 is optional. Study participants will be provided with four units of each study material.

Study Material 1 contains *in vitro* transcribed HIV-1 (HXB2) RNA fragment at an approximate concentration ~$10^2$-$10^4$ copies/µl in ~5 ng/µL human Jurkat cell line total RNA (Ambion "FirstChoice® Human T-Cell Leukemia (Jurkat) Total RNA", P/N AM7858) in buffered solution (1 mM sodium citrate, pH 6.5 (RNA Storage Solution Thermo Fisher Scientific P/N AM7001)). A total of 300 units, each containing 100 µL, were prepared.

Study Material 2 contains *in vitro* transcribed HIV-1 (HXB2) RNA fragment at an approximate concentration of between $10^9$-$10^{10}$ copies/µl (~5-30 ng/µl) in RNA Storage Solution (as above) in a volume of 200 µl per unit. A total of 120 units were prepared. This material does not contain any other nucleic acids and is designed to be suitable for analysis by chemical analysis methods and single molecule flow cytometry.

Study Material 3 is composed of purified HIV-1 viral genomic RNA from viral stocks used to prepare the WHO 3rd and 4th International Standard (IS) for HIV-1 (Gall *et al*., 2014). The material contains a similar *gag* RNA copy number concentration to Study Material 1. Prior to RNA purification, the HIV viral stock was heat inactivated for 1 hour and then tested via tissue culture passage for one month with no viral growth detected. This is confirmed in Appendix A to the Protocol (letter from NIBSC dated 18 Jan. 2019). RNA was purified using the QIAamp® UltraSens® Virus (Qiagen) and diluted in ~5 ng/µl human Jurkat cell line total RNA in RNA Storage Solution (as above). Each unit of material contains 100 µL sample. A total of 300 units were prepared.

#### 3.2 Homogeneity

The homogeneity of all study materials was assessed by performing eight replicate measurements (sub-samplings) of 10 units. The homogeneity of Study Materials 1 and 3 was evaluated by RT-dPCR analysis (BioRad QX200). The homogeneity of Study Material 2 was evaluated by fluorimetric assay (QuantiFluor® RNA System using the Quantus fluorimeter (Promega)). Data was analysed using ANOVA (Materials 1 and 3) and linear mixed effects model with unit and replicate measurement as random effects. The magnitude and statistical significance of between unit variation (standard deviation $s_b$) are given in Table 1.

**Table 1: Results of homogeneity study**

| Study Material | Relative $s_b$ (%)* | Significance ($p$) |
|---|---|---|
| 1 | 2.1% | < 0.001 |
| 2 | 0.36% | *N.S.* |
| 3 | 1.5% | *N.S.* |

Key: *N.S.,* not significant ($p > 0.05$). *Rounded outwards to 2 s.f.

##### 3.3 Stability

A short term stability study was performed by incubation of study materials on dry ice or at raised ambient temperature (~ 27°C) for 1 and 7 days and compared to reference temperature (-80°C) ($n = 3$ units per condition). Stability for Study Materials 1 and 3 was assessed by RT-dPCR ($n = 3$) and for Study Material 2 by fluorometric assay (as Homogeneity study) and Agilent 2100 Bioanalyzer (fragment size). The effect of incubation time and temperature were evaluated using mixed effect models and no significant effect of incubation time was evident, therefore the effect of the two shipment temperatures was compared with the reference temperature using a mixed effect model with temperature as a fixed effect and sample (unit) as a random effect. The results of this analysis are shown in Table 2. Analysis of Study Material 2 using the Agilent Bioanalyzer showed a peak corresponding to the expected size of 2,266 bp was observed under all simulated shipment conditions.

**Table 2: Results of short term stability study**

| **Study Material** | **Effect of 27°C incubation (*vs.* reference -80°C)** Direction, magnitude (significance) | **Effect of dry ice incubation (*vs.* reference -80°C)** Direction, magnitude (significance) |
|---|---|---|
| 1 | ↓8.1% ($p = 0.003$) | ↓2.2% (*N.S.*) |
| 2 | ↓2.6% (*N.S.*) | ↓2.7% (*N.S.*) |
| 3 | ↓15% ($p < 0.001$) | ↓4.4% (*N.S.*) |

Key: *N.S.,* not significant ($p > 0.05$). *Rounded outwards to 2 s.f.

A long term stability study will be commenced prior to shipment of study materials, with analysis of four units of each material performed every three months until completion of the study, with the final analysis being performed after submission of results. Only units stored at the reference temperature of −80° C will be analysed as this is the storage temperature for RNA samples which is recommended to participants.

#### 4. Measurand

The RNA copy number concentration of the *gag* gene expressed in copies per µL (c/µL). This quantity shall be reported for Study Material 1. Analysis of Study Materials 2 and 3 is optional for participants.

#### 5. Shipping and storage of study materials

Study materials will be shipped to participants on dry ice. If the materials have thawed at the point of receipt please request a fresh shipment to be sent.

It is recommended to store the study materials at -80 °C on receipt until analysis. When study materials are to be analysed, they should be thawed, vortexed to briefly to homogenise the contents and kept on ice. It is recommended that study materials do not undergo more than three freeze-thaw cycles.

#### 6. Recommended probe types for use with One-Step RT-ddPCR Advanced Kit for Probes

In the case of adopting a one-step RT-dPCR approach using the QX100 or QX200, it is recommended that participants check the compatibility of fluorescent probe types with One-Step RT-ddPCR Advanced Kit for Probes (Bio-Rad P/N 1864021 and 1864022). Double-quenched probes (such as ZEN probes (IDT)) and Minor Groove Binding Non-Fluorescent Quencher (MGB-NFQ) probes (Thermo Fisher Scientific) have been shown to give better separation of negative and positive droplets and lower background of negative droplets compared to single-quenched probes (Maier *et al.,* 2019; Pinheiro de Oliveira *et al*., 2019; NML at LGC personal communication regarding BHQnova). The NML at LGC has also successfully used BHQnova (Biosearch Technologies) double-quenched probes in combination with the One-Step RT-ddPCR Advanced Kit for Probes.

#### 7. Schedule

| Date | Timeline | Actions (Participants) |
|---|---|---|
| April 2019 CCQM | Call for participation | |
| 17 May 2019 | Final protocol circulated | |
| 31 May 2019 | Registration for participation | Reply with Form 1 |
| 01- 21 June 2019 | Study Material distribution | Reply with Form 2 |
| 16 September 2019 | Submission of results | Reporting with Forms 3-4 |
| 02-04 October 2019 | Initial report at NAWG (Turin) | Presentation of results |

#### 8. Expected outcomes of the study

The study aims to support competencies in targeted measurement of RNA-based materials expressed as copy number concentration. As no primer or probe sequences are recommended, this extends this competency to the design/selection of molecular assays where they are used.

Analysis of Study Material 1 supports measurement claims for quantification of low concentration RNA samples (≤ $10^4$ copies/µl) of RNA templates in a complex matrix (human total RNA) in aqueous solution. Analysis of Study Material 2 supports claims for measurement of high concentration samples of pure RNA templates in aqueous solution. Analysis of Study Material 3 would extend the claim related to Study Material 1 to capability in analysis of viral genomic RNA templates where greater sequence heterogeneity increases the measurement challenge.

Due to the Study Materials being purified RNA samples, the study is not designed to assess the influence of matrix (inhibitors etc) or pre-analytical factors, such as extraction, on RNA measurements.

#### 9. References

Chen WW, Balaj L, Liau LM, Samuels ML, Kotsopoulos SK, Maguire CA, et al. (2013). BEAMing and Droplet Digital PCR Analysis of Mutant *IDH1* mRNA in Glioma Patient Serum and Cerebrospinal Fluid Extracellular Vesicles. Mol. Ther. Nucl. Acids 2:e109.

Devonshire AS, Sanders R, Whale AS, Nixon GJ, Cowen S, Ellison SL, *et al*. (2016). An international comparability study on quantification of mRNA gene expression ratios: CCQM-P103.1. Biomol. Detect. Quantif. 8:15-28.

Gall A, Morris C, Kellam P & Berry N. 2014. Complete Genome Sequence of the WHO International Standard for HIV-1 RNA Determined by Deep Sequencing. Genome Announcements, 2: e01254-13.

Maier J, Lange T, Cross M, Wildenberger K, Niederwieser D, Franke GN (2019). Optimized Digital Droplet PCR for *BCR-ABL*. J. Mol. Diagn. 21(1):27-37.

Pinheiro-de-Oliveira TF, Fonseca-Júnior AA, Camargos MF, Laguardia-Nascimento M, Giannattasio-Ferraz S, Cottorello ACP *et al*. (2019). Reverse transcriptase droplet digital PCR to identify the emerging vesicular virus *Senecavirus A* in biological samples. Transbound. Emerg. Dis. doi.10.1111/tbed.13168.

Sanders R, Mason DJ, Foy CA, Huggett JF. Evaluation of digital PCR for absolute RNA quantification. PLoS One 2013;8:e75296.

Stein EV, Duewer DL, Farkas N, Romsos EL, Wang L, Cole KD. Steps to achieve quantitative measurements of microRNA using two step droplet digital PCR. PLoS One 2017;12:e0188085.

Whale AS, Bushell C, Grant PR, Cowen S, Guttierrez-Aguirre I, O'Sullivan DM, et al. Detection of rare drug resistance mutations by digital PCR in a human influenza A virus model system and clinical samples. J. Clin. Microbiol. 2016;54:392-400.

White RA, 3rd, Quake SR, Curr K. Digital PCR provides absolute quantitation of viral load for an occult RNA virus. J. Virol. Methods 2012;179:45-50.

Yoo HB, Park SR, Dong L, Wang J, Sui Z, Pavsic J, et al. (2016). International Comparison of Enumeration-Based Quantification of DNA Copy-Concentration Using Flow Cytometric Counting and Digital Polymerase Chain Reaction. Anal. Chem. 88:12169-76.

Medicines & Healthcare products Regulatory Agency

NIBSC

18th January 2019

Dear Alison,

Please find enclosed 4 x 60µl HIV-1 RNA extracts. The HIV-1 virus was originally isolated in 1996 post-mortem from a patient that died from an AIDS-defining illness. The virus was subsequently passaged through human PBMC's to increase volume and titre and stored under vapour phase liquid nitrogen.

For the purposes of this study, the virus has been heat inactivated at 60$^0$C for 1 hr and efficacy of inactivation demonstrated by parallel culture of the inactivated material alongside non-inactivated material for a period of one month. No viral growth was detected in the inactivated material. Subsequently, this material has been extracted using the QIAGEN UltraSENS Virus Kit.

The virus has previously been used to produce the 3rd and 4th HIV-1 WHO International standards. The complete genome sequencing of the 4th HIV-1 International Standard has been confirmed by deep sequencing (Genome Announc. 2014 Feb 6;2(1)).

Best Regards,

Clare
