## Supplementary material for "CCQM-P199: Interlaboratory comparability study of HIV-1 RNA copy number quantification": CCQM P199 supplementary file Appendix D

### APPENDIX D: Registration Form

#### Form 1: Confirmation of Participation

| |
|---|
| Organisation name |
| Study Materials to be analysed by organisation (please tick): |
| Study Material 1 |
| Study Material 2 |
| Study Material 3 |
| Contact person and email address |
| Address for sample shipment |
| Contact telephone for sample shipment |
| Special considerations for import |
