## Supplementary material for "CCQM-P199: Interlaboratory comparability study of HIV-1 RNA copy number quantification": CCQM P199 supplementary file Appendix E

### APPENDIX E: Study Material Receipt Form

#### Form 2: Receipt of Study Materials

| | |
|---|---|
| Organisation name | |
| Contact person and email address | |
| Date and time of sample reception | |
| Dry ice present on receipt? (yes/no) | |
| Samples still frozen? (yes/no) | |
| Any sign of sample leakage (yes/no) | |
| Any mishaps during delivery? (yes/no) | If yes, please describe below: |
