## Supplementary material for "CCQM-P199: Interlaboratory comparability study of HIV-1 RNA copy number quantification": CCQM P199 supplementary file Appendix F

### APPENDIX F: Reporting Form

#### Form 3: Submission of Results

| |
|---|
| Organisation name |
| Contact person and email address |

| **MATERIAL** | **MEASURAND** (UNIT) |
|---|---|
| **STUDY MATERIAL 1** | **RNA copy number concentration (*gag* gene)** (copies/µL) |
| Value (*x*) | |
| Standard uncertainty (*u*) | |
| Coverage factor (*k*) | |
| Expanded uncertainty (*U*) | |
| Relative expanded uncertainty (Rel *U*) | |
| **STUDY MATERIAL 2** | **RNA copy number concentration (*gag* gene)** (copies/µL) |
| Value (*x*) | |
| Standard uncertainty (*u*) | |
| Coverage factor (*k*) | |
| Expanded uncertainty (*U*) | |
| Relative expanded uncertainty (Rel *U*) | |
| **STUDY MATERIAL 3** | **RNA copy number concentration (*gag* gene)** |

| | (copies/μL) |
|---|---|
| Value (*x*) | |
| Standard uncertainty (*u*) | |
| Coverage factor (*k*) | |
| Expanded uncertainty (*U*) | |
| Relative expanded uncertainty (Rel *U*) | |
