## Supplementary material for "CCQM-P199: Interlaboratory comparability study of HIV-1 RNA copy number quantification": CCQM P199 supplementary file Appendix G

#### APPENDIX G: Experimental details form

##### Form 4: Experimental details

**(1)** Experimental information**:** Please complete CCQM-P199 dMIQE form.

**(2)** Experimental Design: Please describe or show in diagrammatic form the experimental design which was applied.

**(3)** Measurement uncertainty. Please summarise the calculation approach used and list the factors which were included

##### Form 5: dMIQE checklist

See next page

### CCQM P199 Reply Form 5: dMIQE Information Table

| |
|---|
| Organisation name |
| Contact person and email address |

| ITEM TO CHECK | IMPORTANCE | Comments |
|---|---|---|
| **EXPERIMENTAL DESIGN** | | |
| Definition of experimental and control groups | **E** | *Describe use of positive and negative controls* |
| Number within each group | **E** | *Complete in Reply Form 4 (Experimental Design)* |
| Assay carried out by core lab or investigator's lab? | D | |
| Power analysis | D | N/A |
| **SAMPLE** | N/A | Purified RNA samples only |
| Description | **E** | N/A |
| Volume or mass of sample processed | **E** | N/A |
| Microdissection or macrodissection | **E** | N/A |
| Processing procedure | **E** | N/A |
| If frozen - how and how quickly? | **E** | N/A |
| If fixed - with what, how quickly? | **E** | N/A |
| Sample storage conditions and duration (especially for FFPE samples) | **E** | N/A |
| **NUCLEIC ACID EXTRACTION** | N/A | Purified RNA samples only |
| Quantification - instrument/method | **E** | N/A |
| Storage conditions: temperature, concentration, duration, buffer | **E** | N/A |
| DNA or RNA quantification | **E** | N/A |
| Quality/integrity-instrument/method; e.g. RIN/RQI and trace or 3’:5’ | **E** | N/A |
| Template structural information | **E** | N/A |
| Template modification (digestion, sonication, pre-amplification etc.) | **E** | |

| | | |
|---|---|---|
| Template treatment (initial heating or chemical denaturation) | **E** | |
| Inhibition dilution or spike; | **E** | N/A |
| DNA contamination assessment of RNA sample | **E** | N/A |
| Details of DNase treatment where performed | **E** | N/A |
| Manufacturer of reagents used and catalogue number | D | N/A |
| Storage of nucleic acid: temperature, concentration, duration, buffer | **E** | |
| **REVERSE TRANSCRIPTION** | | |
| One or two step protocol | **E** | |
| *RT priming method + concentration of oligos (for 2-step; if 1-step complete below)* | **E** | |
| *Amount of RNA used per reaction (for 2-step; if 1-step complete below)* | **E** | *(Volume of study material ($\mu l$))* |
| *Detailed reaction components and conditions (for 2-step; if 1-step complete below)* | **E** | |
| RT efficiency | D | |
| Estimated copies measured with and without addition of RT | D | *RT minus results* |
| *Manufacturer of reagents used and catalogue number (for 2-step; if 1-step complete below)* | D | |
| *Reaction volume (for 2-step; if 1-step complete below)* | D | |
| *Storage of cDNA: temperature, concentration, duration, buffer (for 2-step)* | D | |
| **dPCR TARGET INFORMATION** | | |
| Sequence accession number | **E** | |
| Location of amplicon | D | |
| Amplicon length | **E** | |
| In silico specificity screen (BLAST, etc) | **E** | |
| Pseudogenes, retropseudogenes or other homologs? | D | |
| Sequence alignment | D | |
| Secondary structure analysis of amplicon and GC content | D | |
| Location of each primer by exon or intron (if applicable) | E | N/A |
| Where appropriate, which splice variants are targeted? | **E** | N/A |
| **dPCR OLIGONUCLEOTIDES** | | |

| | | |
|---|---|---|
| Primer sequences | **E** | |
| RTPrimerDB Identification Number | D | N/A |
| Probe sequences | D | |
| Location and identity of any modifications | **E** | |
| Manufacturer of oligonucleotides | D | |
| Purification method | D | |
| **dPCR PROTOCOL** | | |
| Complete reaction conditions | **E** | |
| Reaction volume and amount of RNA/cDNA/DNA | **E** | |
| Primer, (probe), Mg++ and dNTP concentrations | **E** | *(if not proprietary)* |
| Polymerase identity and concentration | **E** | |
| Buffer/kit Catalogue No and manufacturer | **E** | |
| Exact chemical constitution of the buffer | D | *(if not proprietary)* |
| Additives (SYBR Green I, DMSO, etc.) | **E** | |
| Plates/tubes Catalogue No and manufacturer | D | |
| Complete thermal cycling parameters | **E** | |
| Reaction setup | D | *(Manual / automatic droplet generation)* |
| Gravimetric or volumetric dilutions (manual/robotic) | D | |
| Total PCR reaction volume prepared | D | |
| Partition number | **E** | |
| Individual partition volume | **E** | |
| Total volume of the partitions measured (effective reaction size) | **E** | |
| **Partition volume measurement uncertainty** | D | *(Standard uncertainty)* |
| Comprehensive details and appropriate use of controls | **E** | |
| Manufacturer of dPCR instrument | **E** | |
| **dPCR VALIDATION** | | |
| Optimisation data for the assay | D | *(Summary of optimisation experiments / representative plots)* |

| | | |
|---|---|---|
| Specificity (when measuring rare mutations, pathogen sequences etc.) | **E** | |
| Limit of detection of calibration control | D | |
| If multiplexing, comparison with singleplex assays | **E** | |
| **DATA ANALYSIS** | | |
| Average copies per partition (λ or equivalent ) | **E** | |
| dPCR analysis program (source, version) | **E** | |
| Outlier identification and disposition | **E** | |
| Results of NTCs (for participant analysis) | **E** | |
| Examples of positive(s) and negative experimental results as supplemental data | **E** | *Please provide amplification plots for (1) Positive control (2) Negative Control (3) Study Material (4) NTC* |
| Where appropriate, justification of number and choice of reference genes | **E** | N/A |
| Where appropriate, description of normalisation method | **E** | N/A |
| Number and concordance of biological replicates | D | N/A |
| Number and stage (RT or qPCR) of technical replicates | **E** | *Complete in Reply Form 4* |
| Repeatability (intra-assay variation) | **E** | |
| Reproducibility (inter-assay/user/lab etc. variation ) (***intermediate precision***) | D | |
| Experimental variance or confidence interval*** (***measurement uncertainty***) | **E** | *Complete in Reply Form 3* |
| Statistical methods used for analysis (***measurement uncertainty***) | **E** | *Complete in Reply Form 4* |
| Data submission using RDML | D | |
