## Supplementary material for "CCQM-P199: Interlaboratory comparability study of HIV-1 RNA copy number quantification": CCQM P199 supplementary file Appendix H

#### APPENDIX H: Summary of Participants' Analytical Information

The following Tables summarize the detailed information about the analytical procedures each participant provided in their "Analytical Information" worksheets. The presentation of the information in many entries has been consolidated and standardized.

The participant's measurement uncertainty statements are provided verbatim in Appendix I.

**Table H-1: Summary of RT-dPCR Analytical information for CCQM P199 Study Materials 1 and 3**

| Lab ID (Result)* | | One or two-step RT-PCR | RT efficiency correction (Y/N) | RT reagent (Manufacturer, Part Number) / procedure (two-step only) | RT-PCR reagent (one-step) or PCR reagent (two-step) | dPCR instrument** | Thermal Cycler (for two-step, state if same or different cyclers used RT and dPCR) |
|---|---|---|---|---|---|---|---|
| 1 | | one-step | N | N/A | One-step RT-ddPCR advanced kit for probes, Bio-Rad 186-4021 | QX200 (manual) | BioRad CFX96 Touch Deep Well |
| 2 | | one-step | Y | N/A | One-step RT-ddPCR advanced kit for probes, Bio-Rad 186-4021 | QX200 (manual) | Bio-Rad C1000 Touch |
| 3 (1) | | one-step | N | N/A | One-step RT-ddPCR advanced kit for probes, Bio-Rad 186-4021 | QX200 (manual) | Thermo Fisher Veriti 96-well |
| 3 (2) | | two-step | N | SuperScript III (Thermo Fisher 18080051). Gene-specific priming. | Bio-Rad ddPCR for probes Catalogue number186-3010 | QX200 (manual) | |
| 4 | | one-step | N | N/A | One-step RT-ddPCR advanced kit for probes, Bio-Rad 186-4022 | QX200 (manual) | Bio-Rad C1000 Touch |
| 5 | | one-step | N | N/A | One-step RT-ddPCR advanced kit for probes, Bio-Rad 186-4022 | QX100 (manual) | Bio-Rad C1000 |
| 6 | | one-step | (Y – supplementary results only) | N/A | One-step RT-ddPCR advanced kit for probes, Bio-Rad 186-4021 | QX200 (manual) | Thermo Fisher Veriti 96 well Thermal Cycler |
| 7 | | one-step | N | N/A | One-step RT-ddPCR advanced kit for probes, Bio-Rad 186-4021 | QX200 (AutoDG) | Bio-Rad T 100 Thermal cycler |
| 8 | | one-step | N | N/A | One-step RT-ddPCR advanced kit for probes, Bio-Rad 186-4022 | QX200 (AutoDG) | ProFlex (Thermo Fisher Scientific) |
| 9 | | one-step | N | N/A | One-step RT-ddPCR advanced kit for probes, Bio-Rad 186-4022 | QX100 (AutoDG) | Bio-Rad C1000 |
| 10 | | two-step | N | SuperScript IV First-Strand Synthesis System (Thermo Fisher Scientific, 18091050) | QuantStudio 3D Digital PCR mix v2, Thermo Fisher Scientific A26358 | QuantStudio 3D Digital PCR system | RT-step: TaKaRa PCR Thermal Cycler (TaKaRa) dPCR step: QuantStudio 3D Digital PCR System (Thermo Fisher Scientific) |

| Lab ID (Result)* | | One or two-step RT-PCR | RT efficiency correction (Y/N) | RT reagent (Manufacturer, Part Number) / procedure (two-step only) | RT-PCR reagent (one-step) or PCR reagent (two-step) | dPCR instrument** | Thermal Cycler (for two-step, state if same or different cyclers used RT and dPCR) |
|---|---|---|---|---|---|---|---|
| 11 | | one-step | N | N/A | One-step RT-ddPCR advanced kit for probes, Bio-Rad 186-4021 | QX200 (manual) | C1000 Touch Thermal Cycler, Bio-Rad 1851197 |
| 12 | | two-step | N | TaqMan RNA-to-CT 1-Step Kit, Thermo Fisher Scientific, 4392653.<br>RT Priming: Reverse oligo (500 nM[1]). | ddPCR Supermix for Probes (No dUTP), Bio-Rad, 186-3023 | QX200 (manual) | Veriti 96-Well Fast Thermal Cycler, ThermoFisher Sci. |
| 12 (S) | | two-step (as main result) | N | As main result. | QuantStudio 3D Digital PCR Master Mix, Thermo Fisher 10027614 | QuantStudio 3D Digital PCR system | Veriti 96-Well Fast Thermal Cycler, ThermoFisher Sci. |
| 13 (1) | | one-step | N | N/A | One-step RT-ddPCR advanced kit for probes, Bio-Rad 186-4022 | QX200, manual DG | N/A |
| 13 (2) | | two-step | N | RevertAid H Minus First Strand cDNA Synthesis Kit (ThermoScientific, #K1632). RT Priming: Reverse oligo 500 nM and random primer. | ddPCR Supermix for Probes, Bio-Rad 186-3026 | QX200, manual DG | CFX96 (same for one- and two-step) |

*For multiple results, result number (for multiple nominated results) or supplementary (S) stated in brackets. **State if manual or auto droplet generation for QX100/QX200.

[1] The molecular biology community express amount of substance concentration with units of molarity (M) (SI units: mmol/L).

**Table H-2: Summary of Analytical Techniques for CCQM P199 Study Material 2**

| Laboratory ID | Analytical Technique | Instrument | Additional information |
|---|---|---|---|
| 2 | RT-dPCR | Bio-Rad (QX200) | One SM2 unit was gravimetrically serially diluted up to ≈ 10 c/µL based on estimations given in study protocol in total RNA ≈ 5 ng/µL (extracted in-house from cultured Jurkat cell line) and then analyzed by RT-ddPCR. RT efficiency was estimated by measurement of SM2 by UV spectroscopy using a Shimadzu UV 2700 spectrophotometer equipped with a TCC 240 thermal control unit. One SM2 unit was gravimetrically serially diluted in water to ≈ 500 µL and then measured in the spectrophotometer. Gravimetric dilution of SM2 was performed at "room temperature" (20 °C - in equilibrium with the temperature in the balance room). Each diluted material was immediately placed on ice after mixing and weighing until further use. Spectrophotometer measurements were also taken with the diluted material equilibrated at "room temperature" in a thermostated cuvette holder set to 20 °C |
| 3 | Flow cytometry | Built in-house | Each SM2 unit was divided into three and hybridised with 70 probes (5'FAM labeled) interspersed within the target RNA. After hybridization the samples were gravimetrically diluted down to adequate concentration for direct counting. The sample was first added directly into hybridization buffer, which was then diluted using 1x TE (Tris EDTA) buffer at ambient temperature (≈ 23 °C to 25 °C). For direct counting, this was further diluted in the FCM running buffer (5 mM Tris-Cl, pH 9.5) (diluted approximately 1:300). |
| 6 | RT-dPCR | Bio-Rad (QX200) | SM2 unit was gravimetrically serially diluted in RSS at ambient temperature. The dilution factor was ≈ $1.8 \times 10^6$ to ≈ $2.7 \times 10^6$.<br>RT efficiency was estimated by measurement of in-house in vitro transcript *gag*-RNA (≈ 3 ng/µL) by nanodrop and RT-dPCR on four vials. |
| 8 | RT-dPCR | Bio-Rad (QX200) | SM2 was volumetrically diluted in water (DF = $10^6$) and then analyzed by RT-ddPCR |
| 9 | RT-dPCR | Bio-Rad (AutoDG & QX100) | For both RT-dPCR and HPLC-UV, the 4 x 200 µL vials of Study Material 2 were equilibrated at room temperature for 30 min, pooled into a 2 mL Axygen tube and homogenized by vortexing. Pooled SM2 was diluted 1:1 wt/wt in 1 mM sodium citrate pH 6.4.<br>For RT-dPCR, a further 4 dilution steps were performed with a proprietary buffer (pH 6.0) at ambient temperature (total dilution factor of $9.2 \times 10^5$) ($n$ = 3 dilutions) for analysis with Assay 1 and Assay 3 (Table H3). |

| Laboratory ID | Analytical Technique | Instrument | Additional information |
|---|---|---|---|
| 9 | HPLC-UV | Dionex (Thermofisher) Ultimate 3000RS Capillary HPLC-UV system | Calibration was performed using NMIJ CRM 6204-b RNA standards (1000A, 1000B) each gravimetrically diluted to ≈ 4 ng/µL.<br>**HPLC:**<br>Column: ProSwift RP-4H 200 µm x 25 cm monolith PS-DVB Serial number 215009<br>Mobile phases: A 200 mM TEAA pH 8.1, B 20 % $CH_3CN$ in A<br>Gradient: 30 % to 90 % B 4 min; 90 % B 6 min; 30 % B 6.5 min; 30 % B 13 min, at 5 uL/min flow<br>Injection volume: 0.5 µL<br>Column oven temp: 40 °C<br>**Detector:** UV 260 nm, Data collection rate: 2.5 Hz, Time constant: 0.6 s<br>The final analysis followed a standard-sample-standard bracketing format (1000A-SM2-1000A, 1000B-SM2-1000B), which was repeated 3 times. Five injections were completed for each standard and sample sequence. |
| 10 | Acid hydrolysis/isotope dilution mass spectrometry | LCMS-8030+ (Shimadzu) | Method described in Shibayama *et al.*, 2016 (1). For the quantification of RNA, optimization of the reaction conditions (especially the reaction time) was performed using NMIJ CRM 6204-b. |
| 11 | Flow cytometry | Home-built flow cytometer with single photon counting fluorescence detection | SM2 was gravimetrically diluted on ice in TE buffer to 4 different concentrations. The final dilution factors were in the range of 1:50 000 to 1:2 000 000. Quant-iT RiboGreen RNA reagent (Cat. no. R11491-Invitrogen) was used for RNA staining. Staining was carried out following the manufacturers protocol for the low-range assay. |

**Table H-3: PCR assay specifications CCQM-P199**

| **Laboratory ID** | **Study Material (specify or write All)** | **Assay name/ reference (if published) (**Duplex if performed) | **Oligonucleotide sequences (5' → 3')** | **Final concentration Forward/Reverse/Probe (nM) (two-step only: reverse primer concentration)** | **Amplicon size (bp)** | **Supplier and purification** |
|---|---|---|---|---|---|---|
| 1 | SM1 | P11-SM1<br>P11-SM1<br>S11-SM1 | Fw: CCCTTTAGAGACTATGTAGACC<br>Rv: GTCTTACAATCTGGGTTCGC<br>Pr: FAM-TTACCTCCTGTGAAGCTTGC-BHQ1Plus | 900/900/300 | 122 | Biosearch technologies. RPC for primers and RP HPLC/dual HPLC for probes |
| 1 | SM3 | P10-SM3<br>P-10-SM3<br>S10-SM3 | Fw: CCCTTTAGGGACTATGTAGAC<br>Rv: GTCTTACAATCTGGGTTTGC<br>Pr: FAM-TTACTTCCTGTGAAGCTTGC-BHQ1plus | 900/900/300 | 122 | Biosearch technologies. RPC for primers and RP HPLC/dual HPLC for probes |
| 2 | All | Fw1381-1400 K03455.1<br>Rv 1505-1484 K03455.1<br>probe 1456-1475 K03455.1 | Fw: CAAGCAGCCATGCAAATGTT<br>Rv: ATGTCACTTCCCCTTGGTTCT<br>Pr: FAM-CCTGGTGCAATAGGCCCTGC-BHQ1 | 900/900/250 | 124 | Exxtend Biotecnologia Ltda. Standard desalting for primers/ RP HPLC for probe |
| 3 (two-step) | All | AssayB_F:<br>AssayB_R:<br>AssayB_P: | Fw: CAGGGGCAAATGGTACATCA<br>Rv: GGGGTGGCTCCTTCTGATAA<br>Pr: FAM/ZEN-AAGAGAAGGCTTTCAGCCCAGAAGT-IBFQ | (two-step primer concentration: 2 µM) 1000/1000/250 | 128 | Integrated DNA technologies (IDT) |
| 3 (one-step) | All | AssayC_F:<br>AssayC_R:<br>AssayC_P: | Fw: CAGAACATCCAGGGGCAAAT<br>Rv: CTGGGCTGAAAGCCTTCTCT<br>Pr: FAM/ZEN- GGTACATCAGGCCATATCACCTAGAAC-IBFQ | 1000/1000/250 | 94 | Integrated DNA technologies (IDT) |

| **Laboratory ID** | **Study Material (specify or write All)** | **Assay name/ reference (if published)** (Duplex if performed) | **Oligonucleotide sequences (5' → 3')** | **Final concentration Forward/Reverse/Probe (nM) (two-step only: reverse primer concentration)** | **Amplicon size (bp)** | **Supplier and purification** |
|---|---|---|---|---|---|---|
| 4 | All | *gag*, Bosman *et al*. 2015 (2) | Fw: TGGGTAAAAGTAGTAGAAGAGAAGGCTTT<br>Rv: CCCCCCACTGTGTTTAGCAT<br>Pr: FAM-TCAGCATTATCAGAAGGAG-MGBEQ | 900/900/200 | 116 | Biosearch technologies. Standard desalting for primers and dual HPLC for probes |
| 5 | All | *gag* | Fw: AGTRGGGGGACAYCARGCAGCHATGCARAT<br>Rv: TACTAGTAGTTCCTGCTATRTCACTTCC<br>Pr: FAM-AT CAA TGA R/ZEN/G ARG CTG CAG AAT GGG A-3IABkFQ | 900/900/250 | 155 | Integrated DNA technologies (IDT). Standard desalting for primers and HPLC for probes |
| 6 | All | *gag Assay 1* (***Error! Reference source not found.***) | Fw: CATGTTTTCAGCATTATCAGAAGGA<br>Rv: TGCTTGATGTCCCCCCACT<br>Pr: FAM- CCACCCCACAAGATTTAAACACCATGCTAA - TAMRA | 600/600/100 | 79 | Sangon biotech (Shanghai, China). HPLC |
| 7 | All | *gag* | Fw: AATTACCCTATAGTGCAGAACATCCA<br>Rv: TCTCTTCTACTACTTTTACCCATGCATT<br>Pr: FAM-CAAATGGTACATCAGGCC- MGB-NFQ | 900/900/250 | 94 | Applied Biosystems. Standard desalting for primers and HPLC for probes |
| 8 | SM1 & 2 | *gag*-2 | Fw: TCCAAAATGCGAACCCAGAT<br>Rv: TCCTCCTACTCCCTGACATG<br>Pr: FAM-AGCATTGGGACCAGCGGCTACACTAGAAGA-MGB-NFQ | 250/250/250 | 98 | Primers: Eurofins/Probe: Applied Biosystems |
| 8 | SM1 & 2 | *gag*-3 | Fw: TGGGATAGAGTGCATCCAGT<br>Rv: GTCACTTCCCCTTGGTTCTC<br>Pr: FAM- GCATGCAGGGCCTATTGCACCAGGC-MGB-NFQ | 250/250/250 | 72 | Primers: Eurofins/Probe: Applied Biosystems |

| **Laboratory ID** | **Study Material (specify or write All)** | **Assay name/ reference (if published)** (Duplex if performed) | **Oligonucleotide sequences (5' → 3')** | **Final concentration Forward/Reverse/Probe (nM) (two-step only: reverse primer concentration)** | **Amplicon size (bp)** | **Supplier and purification** |
|---|---|---|---|---|---|---|
| 8 | SM1 & 2 | *gag*-4 | Fw: ATGAGGAAGCTGCAGAATGG<br>Rv: CTTGGTTCTCTCATCTGGCC<br>Pr: FAM- CCAGTGCATGCAGGGCCTATTGCACC-MGB-NFQ | 250/250/250 | 79 | Primers: Eurofins/Probe: Applied Biosystems |
| 8 | SM1 & 2 | *gag*-5 | Fw: CAATTGTGGCAAAGAAGGGC<br>Rv: TTGGTGTCCTTCCTTTCCAC<br>Pr: FAM- ACACAGCCAGAAATTGCAGGGCCCCT-MGB-NFQ | 250/250/250 | 88 | Primers: Eurofins/Probe: Applied Biosystems |
| 8 | SM3 | A | Fw: ATGAGGAAGCTGCAGAATGG<br>Rv: GTCACTTCCCCTTGGTTCTC<br>Pr: FAM- CAGGCAGGGCCTATTGCACCAGGC-MGB-NFQ | 250/250/250 | 89 | Primers: Eurofins/Probe: Applied Biosystems |
| 8 | SM3 | B | Fw: GGTCCAAAATGCAAACCCAGAT<br>Rv: TCCCCCCACTCCCTGACAT<br>Pr: FAM- AGCATTGGGACCAGCAGCTACACTAGAAGA-MGB-NFQ | 250/250/250 | 100 | Primers: IDT/Probe: Applied Biosystems |
| 8 | SM3 | C | Fw: CAATTGTGGCAAAGAAGGGC<br>Rv: TTGGTGTCCTTCCTCTCCAC<br>Pr: FAM- ACATAGCCAGAAATTGCAGGGCCCCT-MGB-NFQ | 250/250/250 | 88 | Forward primer: Eurofins<br>Reverse primer: IDT<br>Probe: Applied Biosystems |
| 9 | SM1, SM2 and SM3 | Assay 1 | Fw: GGTGCGAGAGCGTCAGTATT<br>Rv: AACATATAGTATGGGCAAGCAGGG<br>Pr: FAM-TCGGTTAAGGCCAGGGGGAAA-BHQ1 | 900/900/250 | 115 | Sigma Aldrich. HPLC |
| 9 | SM1 and SM2 only | Assay 3 | Fw: CAAGGGAAGGCCAGGGAATTT<br>Rv: CCACCAGAAGAGAGCTTCAGG<br>Pr: FAM-AGCAGACCAGAGCCAACAGCC-BHQ1 | 900/900/250 | 70 | Sigma Aldrich HPLC |

| **Laboratory ID** | **Study Material (specify or write All)** | **Assay name/ reference (if published)** (Duplex if performed) | **Oligonucleotide sequences (5' → 3')** | **Final concentration Forward/Reverse/Probe (nM) (two-step only: reverse primer concentration)** | **Amplicon size (bp)** | **Supplier and purification** |
|---|---|---|---|---|---|---|
| 10 | SM1 | *gag* | Fw: ACAGCATGTCAGGGAGTAGGA<br>Rv: TGCCTCTCTGCATCATTATGGT<br>Pr: FAM-CCGGCCATAAGGCAA-MGB-NFQ | 900/900/250<br>(two-step: primer concentration: 500 nM) | 104 | Thermo Fisher Scientific<br>HPLC |
| 10 | SM3 | *gag* | Fw: CACCAGAAGAGAGCTTCAGGTT<br>Rv: GGGTCGTTGCCAAAGAGTGA<br>Pr: FAM-AGAAGATAGACAAGGAACTGTAT-MGB-NFQ | 900/900/250<br>(two-step: primer concentration: 500 nM) | 119 | Thermo Fisher Scientific<br>HPLC |
| 11 (nominated result) | SM1 & 3 | *gag*; Kondo *et al.* 2009 (3) | Fw: AGTRGGGGGACAYCARGCAGCHATGCARAT<br>Rv: TACTAGTAGTTCCTGCTATRTCACTTCC<br>Pr: FAM-ATCAATGAR/ZEN/GARGCTGCAGA ATGGGA-3IABkFQ | 900/900/250 | 154 | Integrated DNA Technologies, Inc. (IDT), HPLC |
| 11 (S1) | SM1 & 3 | *gag*, Bosman *et al.* 2015 (2) | Fw: TGGGTAAAAGTAGTAGAAGAGAAGGCTTT<br>Rv: CCCCCCACTGTGTTTAGCAT<br>Pr: FAM-TCAGCATTA/ZEN/TCAGAAGGAG-3IABkFQ | 900/900/250 | 116 | Integrated DNA Technologies, Inc. (IDT), HPLC |
| 11 (S2) | SM3 | Duplex<br>*gag*; Kondo *et al.* 2009 (3)<br>*pol*; Lim *et al.* 2016 (4) | *gag* Fw: AGTRGGGGGACAYCARGCAGCHATGCARAT<br>*gag* Rv: TACTAGTAGTTCCTGCTATRTCACTTCC<br>*gag* Pr: FAM-ATCAATGAR/ZEN/GARGCTGCAGA ATGGGA-3IABkFQ<br>*pol* Fw: TTTGGAAAGGACCAGC<br>*pol* Rv: CTGCCATCTGTTTTCCATA<br>*pol* Pr: HEX-TGGAAAGGT/ZEN/GAAGGGGCAGT-3IABkFQ | 900/900/250 | 154<br>182 | Integrated DNA Technologies, Inc. (IDT), HPLC |
| 12 | All | *gag* | Fw: GGGAGAATTAGATCGATGGGAAA<br>Rv: ACAGGCCAGGATTAACTGC<br>Pr: FAM-AATCGTTCT/ZEN/AGCTCCCTGCTTGCC-3IBFQ | 900/900/250<br>(two-step primer concentration: 500 nM) | 122 | Integrated DNA Technologies, Inc. (IDT)<br>HPLC |

| **Laboratory ID** | **Study Material (specify or write All)** | **Assay name/ reference (if published)** (Duplex if performed) | **Oligonucleotide sequences (5’ → 3’)** | **Final concentration Forward/Reverse/Probe (nM) (two-step only: reverse primer concentration)** | **Amplicon size (bp)** | **Supplier and purification** |
|---|---|---|---|---|---|---|
| 13 | SM1 | *gag* | Fw: GAGCTAGAACGATTCGCAGTTA<br>Rv: CTGTCTGAAGGGATGGTTGTA<br>Pr: FAM-CTGGCCTGTTAGAAACATCAGAAGGCT-RTQ1 | 500/500/100<br>(two-step primer concentration: 500 nM) | 94 | Syntol (Moscow, Russia)<br>PAAG+HPLC |

**Table H-4: Technique/method-specific parameters for CCQM P199 RT-dPCR**

| Lab ID | Partition volume (nL) | Partition volume standard uncertainty (nL) (if considered[†]) | Partition volume basis* | Mean accepted partition (droplet) number per reaction | SD partition number | Software version |
|---|---|---|---|---|---|---|
| 1 | 0.769 | 0.0105 | 1 (Measurement of one-step RT-dPCR mastermix, Lot No 64146105 | 17058 | 1247 | Bio-Rad QuantaSoft software v1.7 |
| 2 | 0.786 | 0.023 | 2 Ref: (5) | Study Material 1: 11940<br>Study Material 2: 13635<br>Study Material 3: 13809 | Study Material 1: 1096<br>Study Material 2: 2195<br>Study Material 3: 2071 | Bio-Rad QuantaSoft Analysis Pro v1.0596 |
| 3 (one-step) | 0.85 | 5.5 % | 3 | 11657 per reaction<br>21134 merged value used for analysis | 4023 per reaction<br>3980 merged value used for analysis | Bio-Rad QuantaSoft v1.7.4.0917 |
| 3 (two-step) | 0.85 | 5.5 % | 3 | 10806 per reaction<br>21543 merged value used for analysis | 1542 per reaction<br>3938 merged value used for analysis | Bio-Rad QuantaSoft v1.7.4.0917 |
| 4 | 0.776 | 0.0403 (5.2 %) | 2 (6-8) + 4 (reagent comparison with Supermix w/o dUTP) | 15636 | 1445 | Bio-Rad QuantaSoft v1.7.4.0917 |
| 5 | 0.85 | 2.9 % | 2 | 14307 | 1978 | Bio-Rad QuantaSoft v1.7.4 |
| 6 | 0.71 | 0.4 % | 1 (measurement of RT-dPCR mastermix based on 812 droplets' image) | 15033 | 1684 | Bio-Rad QuantaSoft v1.7.4 |
| 7 | 0.85 | 0.01050 (1.2 %) | 3 | 15774 | 1048 | Bio-Rad QuantaSoft v.1.7.4.0917 |

| **Lab ID** | **Partition volume (nL)** | **Partition volume standard uncertainty (nL) (if considered†)** | **Partition volume basis*** | **Mean accepted partition (droplet) number per reaction** | **SD partition number** | **Software version** |
| --- | --- | --- | --- | --- | --- | --- |
| 8 | 0.7472 | 0.0130830 | 1 + 4 (reagent comparison with Supermix w/ and w/o dUTP) | Study Material 1: 12846<br>Study Material 2: 11999<br>Study Material 3: 12282 | 1730<br>1566<br>2571 | Bio-Rad QuantaSoft v1.7.4.0917 |
| 9 | 0.735 | 0.012 | 1 | Study Material 1: 12335<br>Study Material 2: 12544<br>Study Material 3: 12433 | Study Material 1: 1941<br>Study Material 2: 1228<br>Study Material 3: 1492 | Bio-Rad QuantaSoft v1.7.4<br>In-House analysis v3.0 (Excel 2013) |
| 10 | 0.753 | 0.00478 | 1 | Study Material 1: 17317<br>Study Material 3: 16510 | Study material 1: 885<br>Study material 3: 1263 | QuantStudio 3D Analysis Suite Cloud Software |
| 11 | 0.880 | 0.072 | 1 (light microscopy) (volume uncertainty indicates standard deviation of distribution of single droplet size) | Singleplex (Study Material 1 and 3): 13006<br>Duplex (Study Material 3): 14419 | Singleplex: 1448<br>Duplex: 1579 | Bio-Rad QuantaSoft version 1.7.4.0917 |
| 12 (RT-ddPCR) | 0.834 | NA | 2 | Study material 1: 12802 | 3163 | QuantaSoft Analysis Pro, v1.0.596 |
| 12 (RT-cdPCR) | 0.755 | 3 % (%CV manufacturer) | 3 (chip version 2) | Study material 1: 17106 | 382 | QS3D cloud, version: 3.1.5-PRC-build4 |
| 13 | 0.79 | 3.5 % | 1 | one-step: 12577<br>two-step: 15623 | one-step:1550<br>two-step: 1886 | Bio-Rad QuantaSoft version 1.7.4.0917 |

† Uncertainty values given as reported by participants
‡ NA – not applicable

*Basis for partition volume value: 1 = in-house direct measurement (microscopy); 2 = Literature; 3 = Manufacturer's value; 4 = other

**Table H-5: Reverse transcription and PCR thermal cycling parameters for CCQM P199 RT-dPCR**
**Table H5.A: Reverse transcription parameters (Two-step RT-dPCR only)**

| Laboratory ID | RT temperature (°C) | RT time (min) | RT inactivation temperature (°C) | RT inactivation time (min) | Final incubation Temperature (°C) (time) |
|---|---|---|---|---|---|
| 3 (two-step) | 50 | 50 | 85 | 5 | N/A |
| 10 | 55 | 10 | 80 | 10 | 10 (hold) |
| 12 (two-step) | 37 | 60 | 95 | 5 | N/A |
| 13 (two-step) | 45 | 60 | N/A | N/A | N/A |

**Table H5.B: Reverse transcription and PCR parameters (One-step RT-dPCR and Two-step RT-dPCR (dPCR only))**

| Laboratory ID | RT temp (°C) | RT time (min) | PCR intial step temp | PCR intial step time (min) | PCR cycling temp 1 (°C) | PCR cycling time 1 (s) | PCR cycling temp 2 (°C) | PCR cycling time 2 (s) | Cycle number | PCR final incubation temperature (time) | Ramp rate (ddPCR only) |
|---|---|---|---|---|---|---|---|---|---|---|---|
| 1 | 50 | 60 | 95 | 10 | 95 | 30 | 60 | 60 | 40 | 98 | 1 ºC/s |
| 2 | 45 | 60 | 95 | 10 | 95 | 30 | 55 | 60 | 40 | 98°C (10 min)<br>37°C (10 min) | |
| 3 (one-step) | 42 | 60 | 95 | 10 | 95 | 30 | 60 | 150 | 70 | 98°C (10 min) | 1.5 ºC/s |
| 3 (two-step) | N/A | N/A | 95 | 10 | 94 | 30 | 58 | 60 | 50 | 98 | 2 ºC/s |
| 4 | 47.5 | 60 | 95 | 10 | 94 | 30 | 58 | 60 | 40 | 90 | 2 ºC/s |
| 5 | 50 | 60 | 95 | 10 | 95 | 30 | 55 | 60 | 45 | 98 | |
| 6 | 45 | 10 | 95 | 10 | 95 | 15 | 57 | 60 | 40 | 98 | |
| 7 | 45 | 60 | 95 | 10 | 94 | 30 | 60 | 60 | 40 | 98 | 2 ºC/s |
| 8 *gag*-2, 3, A & C | 50 | 60 | 95 | 10 | 95 | 30 | 57 | 60 | 60 | 98 | |
| 8 *gag*-4 | 50 | 60 | 95 | 10 | 95 | 30 | 58 | 60 | 60 | 98 | |
| 8 *gag*-5 | 50 | 60 | 95 | 10 | 95 | 30 | 56 | 60 | 60 | 98 | |
| 8 B | 50 | 60 | 95 | 10 | 95 | 30 | 55 | 60 | 60 | 98 | |
| 9 | 45 | 60 | 95 | 10 | 95 | 30 | 60 | 60 | 40 | 98 | 2.5 °C/s |
| 10 | N/A | N/A | 96 | 10 | 56 | 120 | 98 | 30 | 50 | 56°C (10 min)<br>10°C (hold) | N/A |

| Laboratory ID | RT temp (°C) | RT time (min) | PCR intial step temp | PCR intial step time (min) | PCR cycling temp 1 (°C) | PCR cycling time 1 (s) | PCR cycling temp 2 (°C) | PCR cycling time 2 (s) | Cycle number | PCR final incubation temperature (time) | Ramp rate (ddPCR only) |
|---|---|---|---|---|---|---|---|---|---|---|---|
| 11 *gag* assays (Kondo & Bosman) | 50 | 60 | 95 | 10 | 95 | 30 | 58 | 60 | 45 | 98 | 2 ºC/s |
| 11 duplex assay (*gag*+*pol*) | 50 | 60 | 95 | 10 | 95 | 30 | 55 | 60 | 45 | 98 | 2 ºC/s |
| 12 RT-ddPCR | N/A | N/A | 95 | 10 | 94 | 30 | 61 | 60 | 40 | 98°C (10 min)<br>4°C (hold) | 2 °C/s |
| 12 RT-cdPCR | N/A | N/A | 95 | 10 | 94 | 30 | 61 | 60 | 39 | 10°C (hold) | N/A |
| 13 (one-step) | 50 | 60 | 95 | 10 | 95 | 30 | 57 | 60 | 54 | 98 (10 min) | 2 ºC/s |
| 13 (two-step) | N/A | N/A | 95 | 10 | 95 | 30 | 57.6 | 60 | 45 | 98°C (10 min)<br>4°C (hold) | 2 ºC/s |

**Table H-6: Additional Comments for CCQM-P199**

| Lab ID | Additional Comments |
|---|---|
| 1 | A 74 nucleotide RNA internal quality control was designed with a sequence based on the K03455.1 gp120 reference genome and was synthesised by Integrated DNA technologies (IDT). Based on the molecular weight and the amount reported in the specification sheet for the RNA fragment, a stock solution was prepared gravimetrically. A reference value with uncertainty was calculated for this control (≈ 24359 /µL ± 495 /µL). The internal amplification control was validated by ddPCR in simplex and duplex form with SM1 and SM3 target sequences. From these results a value for RT efficiency was calculated.<br>IQC: $7.35 \times 10^{14}$ copies ± $1.47 \times 10^{13}$ copies; the assigned value was calculated as the average of reported values in the specification sheet, measured by optical density at 260 nm with UV spectroscopy in nmol and ug units, once they were converted in copies using Avogadro constant.<br><br>The uncertainty of IQC stock was calculated as a type B uncertainty, from the concentration values in copies (calculated from nmol and ug reported values), assuming a rectangular distribution.<br><br>IQC was reconstituted in 300 µL of TE buffer (1X, pH 8.0) to produce the IQC stock, and then gravimetrically diluted to a final concentration of 24359 copies/µL ± 495 copies/µL (int_Q), where the uncertainty was calculated from the dilution factor model, the IQC stock uncertainty and the balance certificate, according to GUM.<br><br>RT efficiency (RT eff) was calculated as the ratio between the measured Int_Q copy number concentration in the master mix and the expected value according to the gravimetric dilution (in the master mix).<br><br>RT eff = experimental Int Q value / expected Int_Q value<br><br>RT efficiency uncertainty was calculated according to previous equation, and a repeatability component; combined following GUM. |
| 2 | SM2 RT efficiency correction factor was calculated as the ratio *copy number concentration UV / copy number concentration dPCR*. Its uncertainty was estimated by combining both UV and dPCR copy number concentration uncertainties. SM2 RNA copy number concentration was calculated through information available in "CCQM P199 HIV-1 target sequence information v1.0 date 14 january 2019". Target sequence extinction coefficient was estimated via nearest-neighbor approach (Nucleic Acids Res. 2004 Jan 13;32(1):e13; Nucleic Acids Res. 2008 Jul 1; 36) assuming 100 % purity. Target sequence molecular weight was estimated according Nucleic Acids Res. 2008 Jul 1; 36 (assuming 100 % purity); SM2 was gravimetrically diluted with water, mixed thoroughly, and UV absorption was determined on a quartz cuvette. RNA copy number concentration was deduced from UV absorption @ 260 nm (10 mm optical path, 20 °C); SM2 RNA copy number concentration uncertainty was estimated from spectrophotometer repeatability, optical path length uncertainty, and molar extinction coefficient uncertainty (set as 2.5 % epsilon). Uncertainty contribution due to gravimetric dilution was shown to be negligible. SM2 was gravimetrically diluted and assayed via dPCR as described for SM1 and SM3. Template gravimetric dilution was computed on uncertainty budget.<br>Shimadzu UV2700 spectrophotometer coupled with TCC-240 Thermoelectrically Temperature Controlled Cell Holder was used in all deteminations.<br><br>Assigned value and uncertainty of SM2: 6.68E+09 cp/µL ± 1.40E+09 cp/µL (U, k=2)<br>Average value and uncertainty of RT efficiency: Value 1.11; U = 7.1 % (0.0715) (k=2)<br>SM2 RT efficiency correction factor and it relative % u was adopted to all study materials<br>Thermo Fisher LTR assay used for preliminary performance comparison. |
| 4 | Prior to analysis by RT-dPCR, all nucleic acid templates (SM1, SM2, SM3) were heat denatured for 5 minutes at 65 °C, then quenched on ice prior to loading for at least 1 min. |
| 6 | RT efficiency was measured based on UV measurement with a NIMC control material ((8.05±0.37) E+9) and the result with correction of RT was not the nominated result.<br>The average value for RT efficiency is 0.72 and its uncertainty is 6.2 %. |

| | |
|---|---|
| 8 | Data reported were the combination of data obtained with 4 different *gag* assays (Materials 1 & 2) or 3 different assays (Material 3) (using the NIST Consensus builder (https://consensus.nist.gov) and the DerSimonian-Laird method of analysis) |
| 9 | All four vials of Study Material 1 and 3 were thawed on ice for 30 min then pooled into an Axygen 1.5 mL tube (MCT-150-L-C) (total sample volume 400 µL), heated to 60 °C for 10 min, and homogenized by vortexing. No heat treatment of Study Material 2 was performed. |

**Table H-7:** **Study Material 2 impurity analysis (UPLC methodology) (Laboratory 10)**

| UPLC parameter | Information |
|---|---|
| Column | Acquity UPLC Protein BEH (Ethylene Bridged Hybrid) SEC (size exclusion chromatography) column (Waters) |
| Elution buffer | 0.1 M Tris-HCl (pH 8.1) |
| Flow rate / temperature | 0.4 ml/min (25 °C) |
| Injection volume | 5 µL |
| Detection | 260 nm (PDA) |

### REFERENCES

1. Shibayama S, Fujii S-i, Inagaki K, Yamazaki T, Takatsu A. Formic acid hydrolysis/liquid chromatography isotope dilution mass spectrometry: An accurate method for large DNA quantification. Journal of Chromatography A. 2016;1468:109-15.
2. Bosman KJ, Nijhuis M, van Ham PM, Wensing AM, Vervisch K, Vandekerckhove L, De Spiegelaere W. Comparison of digital PCR platforms and semi-nested qPCR as a tool to determine the size of the HIV reservoir. Scientific Reports. 2015;5:13811.
3. Kondo M, Sudo K, Tanaka R, Sano T, Sagara H, Iwamuro S, Takebe Y, Imai M, Kato S. Quantitation of HIV-1 group M proviral DNA using TaqMan MGB real-time PCR. Journal of Virological Methods. 2009;157.
4. Lim K, Park M, Lee MH, Woo HJ, Kim J-B. Development and Assessment of New RT-qPCR Assay for Detection of HIV-1 Subtypes. Biomedical Science Letters. 2016;22(3):83-97.
5. Emslie KR, JL HM, Griffiths K, Forbes-Smith M, Pinheiro LB, Burke DG. Droplet Volume Variability and Impact on Digital PCR Copy Number Concentration Measurements. Anal Chem. 2019;91(6):4124-31.
6. Dagata JA, Farkas N, Kramar JA. Method for Measuring the Volume of Nominally 100 µm Diameter Spherical Water-in-Oil Emulsion Droplets. NIST Special Publication 260-184. 2016.
7. Košir AB, Divieto C, Pavšič J, Pavarelli S, Dobnik D, Dreo T, Bellotti R, Sassi MP, Žel J. Droplet volume variability as a critical factor for accuracy of absolute quantification using droplet digital PCR. Anal Bioanal Chem. 2017;409(28):6689-97.
8. Pinheiro LB, O'Brien H, Druce J, Do H, Kay P, Daniels M, You J, Burke D, Griffiths K, Emslie KR. Interlaboratory Reproducibility of Droplet Digital Polymerase Chain Reaction Using a New DNA Reference Material Format. Anal Chem. 2017;89(21):11243-51.
