## Supplementary material for "CCQM-P199: Interlaboratory comparability study of HIV-1 RNA copy number quantification": CCQM P199 supplementary file Appendix I

### APPENDIX I: Summary of Participants’ Uncertainty Estimation Approaches

#### Laboratory 1

SM1

<table>
<tr><th>Measurement uncertainty factor</th><th>Relative standard uncertainty</th><th>Degrees of freedom</th><th>% of combined uncertainty</th></tr>
<tr><td>Lambda (λ)</td><td>4.25 %</td><td rowspan="6">The reported value was calculated from the mean of three measurements in three bottles in three replicas in three different days (one day per bottle), then the degrees of freedom used to calculate the t factor for the coverage factor was 9.</td><td>42 %</td></tr>
<tr><td>Droplet volume</td><td>4.0 %</td><td>4 %</td></tr>
<tr><td>Dilution</td><td>0.03 %</td><td>0.0028 %</td></tr>
<tr><td>RT efficiency</td><td>4.11 %</td><td>39 %</td></tr>
<tr><td>Precision</td><td>2.57 %</td><td>15 %</td></tr>
<tr><td>Combined standard uncertainty</td><td></td><td>100 %</td></tr>
<tr><td>Model</td><td>$RNA\frac{copies}{\mu L} = \frac{\lambda * P}{v * D * E}$</td><td>Where:<br>$\lambda$ Copies/partition<br>$v$ Partition volume<br>$D$ gravimetric dilution<br>$P$ Precision<br>$E$ RT efficiency</td><td></td></tr>
</table>

SM3

<table>
<tr><th>Measurement uncertainty factor</th><th>Relative standard uncertainty</th><th>Degrees of freedom</th><th>% of combined uncertainty</th></tr>
<tr><td>Lambda (λ)</td><td>11.18 %</td><td rowspan="6">The reported value was calculated from the mean of three measurements in three bottles in three replicas in three different days (one day per bottle), then the degrees of freedom used to calculate the t factor for the coverage factor was 9.</td><td>77 %</td></tr>
<tr><td>Droplet volume</td><td>1.36 %</td><td>1 %</td></tr>
<tr><td>Dilution</td><td>0.36 %</td><td>0 %</td></tr>
<tr><td>RT efficiency</td><td>4.24 %</td><td>11 %</td></tr>
<tr><td>Precision</td><td>4.20 %</td><td>11 %</td></tr>
<tr><td>Combined standard uncertainty</td><td></td><td>100 %</td></tr>
</table>

### Laboratory 2

SM1

| **Measurement uncertainty factor** | **Standard uncertainty (copies/μL)** | **Degrees of freedom** | **% of combined uncertainty** |
|---|---|---|---|
| Precision (repeatability) | 6.303216072 | infinite | 66.9 |
| λ | 3.97516E-05 | infinite | 0.0 |
| Droplet volume | 9.12095E-12 | infinite | 0.0 |
| Gravimetric dilutions | 3.77829E-08 | infinite | 0.0 |
| Density , template RNA | 2.72926E-06 | infinite | 0.0 |
| Density , PCR mix | 4.434303 | infinite | 33.1 |
| Combined standard uncertainty | 7.706722521 | infinite | 100 % |

SM2

| **Measurement uncertainty factor** | **Standard uncertainty (copies/μL)** | **Degrees of freedom** | **% of combined uncertainty** |
|---|---|---|---|
| *Precision (repeatability)* | 30.377886 | infinite | 36.0 |
| λ | 4.688E-06 | infinite | 0.0 |
| *Droplet volume* | 3.803E-08 | infinite | 0.0 |
| Gravimetric dilutions | 1.4026629 | infinite | 0.0 |
| Density, template RNA | 2618175.6 | infinite | 0.0 |
| Density, PCR mix | 419024321 | infinite | 0.0 |
| Correction factor, UV | 7.8253 % | infinite | 64.0 |
| Combined standard uncertainty | 6.99E+08 | infinite | 100 % |

SM3

| **Measurement uncertainty factor** | **Standard uncertainty (copies/μL)** | **Degrees of freedom** | **% of combined uncertainty** |
|---|---|---|---|
| *Precision (repeatability)* | 2.605414 | infinite | 82.4 |
| λ | 0.008303 | infinite | 0.0 |
| *Droplet volume* | 2.16E-12 | infinite | 0.0 |
| Gravimetric dilutions | 1.35E-08 | infinite | 0.0 |
| Density, template RNA | 6.45E-07 | infinite | 0.0 |
| Density, PCR mix | 1.204826 | infinite | 17.6 |

| | | | |
|---|---|---|---|
| Combined standard uncertainty | 2.870516 | infinite | 100 % |

### Laboratory 3

SM1, SM2, SM3

| **Measurement uncertainty factor** | **Relative standard uncertainty** | **Degrees of freedom** | **% of combined uncertainty** |
|---|---|---|---|
| *Precision (repeatability)* | SM1(one-step) 1.2 %<br>SM1(two-step) 2.9 %<br>SM3(one-step) 2.3 %<br>SM3(two-step) 3.0 % | | Type A<br>SM1(one-step) 3.3 %<br>SM1(two-step) 8.6 %<br>SM3(one-step) 3.1 %<br>SM3(two-step) 8.7 % |
| *intermediate precision (between assays)* | SM1(one-step) 3.2 %<br>SM1(two-step) 8.1 %<br>SM3(one-step) 2.0 %<br>SM3(two-step) 8.1 % | | |
| *partition volume* | 5.5 % | | Type B<br>SM1(one-step) 5.9 %<br>SM1(two-step) 5.9 %<br>SM3(one-step) 3.1 %<br>SM3(two-step) 5.7 % |
| *manual thresholding* | 0.7 % | | |
| sample homogeneity | SM1: 2.1 %<br>SM3: 1.5 % | | |
| Combined standard uncertainty | | | (A with B)<br>SM1(one-step) 6.8 %<br>SM1(two-step) 10.4 %<br>SM3(one-step) 6.5 %<br>SM3(two-step) 10.4 % |

### Laboratory 4

### Study Material 1

| **Measurement uncertainty factor** | **Relative standard uncertainty** | **Degrees of freedom*** | **% of combined uncertainty** |
|---|---|---|---|
| *Precision* | 2.45 % | 2 | 9.2 % |
| *Bias (Assay)* | 5.4 % | 2 | 44.6 % |

| | | | |
|---|---|---|---|
| *Uncertainty in Bias (Assay)* | 1.76 % | 2 | 4.7 % |
| *Partition volume (Supermix without dUTP)* | 5.1 % | 3 | 0.9 % |
| *Partition volume (Supermix/RTddPCR ratio)* | 0.78 % | 1 | 9.2 % |
| Combined standard uncertainty (rel) | 8.09 % | 5 | 100 % |

### Study Material 3

| **Measurement uncertainty factor** | **Relative standard uncertainty** | **Degrees of freedom*** | **% of combined uncertainty** |
|---|---|---|---|
| *Precision* | 3.27 % | 4 | 21.2 % |
| *Bias (Assay)* | 3.6 %* | 2 | 25.6 % |
| *Partition volume (Supermix without dUTP)* | 5.1 % | 3 | 51.5 % |
| *Partition volume (Supermix/RTddPCR ratio)* | 0.78 % | 1 | 1.2 % |
| Combined standard uncertainty (rel) | 7.11 % | 6 | 100 % |

**There was a calculation error in the measurement uncertainty calculation for Study Material 3. The uncertainty related to Assay bias was intended to be the same as for Study Material 1.*

### Laboratory 5

| **Measurement uncertainty factor** | **Relative standard uncertainty** | **Relativestandard uncertainty** | **Degrees of freedom*** | **Degrees of freedom*** | **% of combined uncertainty** | **% of combined uncertainty** |
|---|---|---|---|---|---|---|
| | SM1 | SM3 | SM1 | SM3 | SM1 | SM3 |
| *Precision (repeatability)* | 1.6 % | 2.7 % | 14 | 14 | 22 | 27 |
| *Droplet volume* | 2.9 % | 2.9 % | | | 30 | 29 |
| *Pipeting* | 3 % | 3 % | | | 31 | 29 |
| *Homogeneity* | 2.1 % | 1.5 % | | | 17 | 15 |
| Combined standard uncertainty | | | | | 100 % | 100 % |

### Laboratory 6

### Study Material 1

| **Measurement uncertainty factor** | **Relative standard uncertainty (%)** | **Degrees of freedom*** | **% of combined uncertainty** |
|---|---|---|---|
| $u_{\text{precision}}$ | 1.2 | 3 | 3.50 |
| $u_{dilution\ factor}$ | 1.9 | 1 | 8.54 |
| $u_{partition\ volume}$ | 5.3 | 63 | 66.07 |
| $u_{\text{stability}}$ | 2.2 | / | 11.45 |
| $u_{\text{homogeniety}}$ | 2.1 | / | 10.44 |
| Combined standard uncertainty | 6.5 | / | 100 |

### Study Material 2

| **Measurement uncertainty factor** | **Relative standard uncertainty (%)** | **Degrees of freedom*** | **% of combined uncertainty** |
|---|---|---|---|
| $u_{\text{precision}}$ | 2.2 | 12 | 10.43 |
| $u_{dilution\ factor}$ | 2.2 | 1 | 10.79 |
| $u_{partition\ volume}$ | 5.3 | 63 | 62.24 |
| $u_{\text{stability}}$ | 2.7 | / | 16.25 |
| $u_{\text{homogeniety}}$ | 0.4 | / | 0.29 |
| Combined standard uncertainty | 6.7 | | 100 |

### Study Material 3

| Measurement uncertainty factor | Relative standard uncertainty (%) | Degrees of freedom* | % of combined uncertainty |
|---|---|---|---|
| $u_{\text{precision}}$ | 5.1 | 3 | 29.95 |
| $u_{\textit{dilution factor}}$ | 3.4 | 1 | 13.26 |
| $u_{\textit{partition volume}}$ | 5.3 | 63 | 32.02 |
| $u_{\text{stability}}$ | 4.4 | / | 22.20 |
| $u_{\text{homogeniety}}$ | 1.5 | / | 2.58 |
| Combined standard uncertainty | 9.3 | / | 100 |

### Laboratory 7

| Measurement uncertainty factor | Relative standard uncertainty | Degrees of freedom* | % of combined uncertainty |
|---|---|---|---|
| *SM1* | | | |
| Method precision | 0.0421 | 3 | 72.5 |
| Droplet volume | 0.0124 | ∞ | 21.3 |
| Volume (Volumetric) | 0.0036 | ∞ | 6.2 |
| Combined standard uncertainty | | | 100 % |
| *SM3* | | | |
| Method precision | 0.0658 | 3 | 80.5 |
| Droplet volume | 0.0124 | ∞ | 15.1 |
| Volume (Volumetric) | 0.0036 | ∞ | 4.4 |
| Combined standard uncertainty | | | 100 % |

### Laboratory 8

Part A: Uncertainties for each assay were calculated using the NIST uncertainty machine (uncertainty.nist.gov)

| Measurement uncertainty factor (SM1) | Mean (copies/ul) [for assay figure] | Standard uncertainty (u) [for assay figure] | Relative standard uncertainty (%) | uncertainty contribution lambda (%) | uncertainty contribution partiton volume (%) | Degrees of freedom* | % of combined uncertainty |
|---|---|---|---|---|---|---|---|
| gag-2 standard uncertainty | 766.5 | 15.3 | 2.0 % | 22.82 | 77.18 | N/A | N/A |
| gag-3 standard uncertainty | 847.6 | 11.9 | 1.4 % | 93.58 | 6.42 | N/A | N/A |
| gag-4 standard uncertainty | 868.5 | 10.5 | 1.2 % | 91.21 | 8.79 | N/A | N/A |
| gag-5 standard uncertainty | 866.9 | 13.4 | 1.5 % | 94.68 | 5.32 | N/A | N/A |

| Measurement uncertainty factor (SM2) | Mean (copies/ul) [for assay figure] | Standard uncertainty (u) [for assay figure] | Relative standard uncertainty (%) | uncertainty contribution lambda (%) | uncertainty contribution partiton volume (%) | Degrees of freedom* | % of combined uncertainty |
|---|---|---|---|---|---|---|---|
| gag-2 standard uncertainty | 4.48E+09 | 1.00E+08 | 2.2 % | 38.66 | 61.34 | N/A | N/A |
| gag-3 standard uncertainty | 4.67E+09 | 2.56E+08 | 5.5 % | 89.79 | 10.21 | N/A | N/A |
| gag-4 standard uncertainty | 4.66E+09 | 2.73E+08 | 5.9 % | 91.05 | 8.95 | N/A | N/A |
| gag-5 standard uncertainty | 4.82E+09 | 3.25E+08 | 6.7 % | 93.27 | 6.73 | N/A | N/A |

| Measurement uncertainty factor (SM3) | Mean (copies/ul) [for assay figure] | Standard uncertainty (u) [for assay figure] | Relative standard uncertainty (%) | uncertainty contribution lambda (%) | uncertainty contribution partiton volume (%) | Degrees of freedom* | % of combined uncertainty |
|---|---|---|---|---|---|---|---|
| gag-A standard uncertainty | 181.2 | 4.77 | 2.6 % | 55.77 | 44.23 | N/A | N/A |
| gag-B standard uncertainty | 169.3 | 3.88 | 2.3 % | 41.56 | 58.44 | N/A | N/A |

| | | | | | | | |
|---|---|---|---|---|---|---|---|
| gag-C standard uncertainty | 185.5 | 4.68 | 2.5 % | 51.75 | 48.25 | N/A | N/A |

Part B: Results and standard uncertainties for each assay were combined using the NIST consensus builder (consensus.nist.gov) using DerSimonian Laird with a coverage probability of 0.99 and a random number generator seed = 5.

### Laboratory 9

SM1

| **Measurement uncertainty factor** | **Relative standard uncertainty** | **Degrees of freedom** | **% of combined uncertainty** |
|---|---|---|---|
| Droplet volume (Type B) | 1.24 % | 59 | 5.1 % |
| Between Assays | 3.74 % | 3 | 46.4 % |
| Between subsamples | 1.65 % | 3 | 9.0 % |
| Within subsamples | 3.45 % | 13 | 39.4 % |
| Dilutions | 0.14 % | 105 | 0.07 % |
| Combined standard uncertainty | 5.5 % | 11.5 | 100 % |

SM2 dPCR

| **Measurement uncertainty factor** | **Relative standard uncertainty** | **Degrees of freedom** | **% of combined uncertainty** |
|---|---|---|---|
| Droplet volume (Type B) | 1.24 % | 59 | 9.9 % |
| Between Assays | 3.02 % | 3 | 58.6 % |
| Between independent runs | 1.52 % | 3 | 14.9 % |
| Within independent runs | 1.60 % | 13 | 16.5 % |
| Dilutions | 0.13 % | 222 | 0.11 % |
| Combined standard uncertainty | 3.9 % | 8.0 | 100 % |

SM2 HPLC

| Measurement uncertainty factor | Relative standard uncertainty | Degrees of freedom | % of combined uncertainty |
|---|---|---|---|
| Precision | 3.83 % | 5 | 22.7 % |
| Calibration reference material | 4.27 % | 100 | 28.2 % |
| Chromatographic resolution | 2.89 % | 60 | 12.9 % |
| Stability of reference material | 4.83 % | 20 | 36.0 % |
| Dilutions | 0.14 % | 100 | 0.03 % |
| Combined standard uncertainty | 8.0 % | 55.8 | 100 % |

SM3

| Measurement uncertainty factor | Relative standard uncertainty | Degrees of freedom | % of combined uncertainty |
|---|---|---|---|
| Droplet volume (Type B) | 1.24 % | 59 | 2.5 % |
| Between independent runs | 3.28 % | 5 | 17.8 % |
| Within independent runs | 5.86 % | 17 | 56.6 % |
| Dilutions | 0.10 % | 74 | 0.02 % |
| Between Assays | 3.74 % | 3 | 23.1 % |
| Combined standard uncertainty | 7.8 % | 23.3 | 100 % |

### Laboratory 10

SM1

| Measurement uncertainty factor | Relative standard uncertainty (%) | Degrees of freedom | % of combined uncertainty |
|---|---|---|---|
| Between preparation of dPCR reaction solution | 2.90 | 2 | 55.6 % |
| Between reverse transcription | 0.69 | 6 | 13.2 % |
| Between chips | 0.95 | 18 | 18.2 % |
| Preparation of dPCR reaction solution (balance) | 0.02 | - | 0.3 % |
| Preparation reverse transcription reaction solution (balance) | 0.03 | - | 0.5 % |
| Partition volume | 0.635 | - | 12.2 % |
| Combined standard uncertainty | 3.19 | | 100 % |

SM2

| Measurement uncertainty factor | Relative standard uncertainty (%) | Degrees of freedom | % of combined uncertainty |
|---|---|---|---|
| IDMS quantification for AMP | 0.49 | - | 7.4 % |
| IDMS quantification for CMP | 1.40 | - | 21.1 % |
| IDMS quantification for GMP | 0.39 | - | 5.9 % |
| IDMS quantification for UMP | 0.53 | - | 8.0 % |
| Between NMPs | 3.51 | - | 53.0 % |
| Density of SM2 | 0.31 | - | 4.7 % |
| Combined standard uncertatinty | 3.88 | | 100 % |

SM3

| Measurement uncertainty factor | Relative standard uncertainty (%) | Degrees of freedom | % of combined uncertainty |
|---|---|---|---|
| Between preparation of dPCR reaction solution | 3.39 | 2 | 38.9 % |
| Between reverse transcription | 2.27 | 6 | 26.1 % |
| Between chips | 2.37 | 18 | 27.2 % |
| Preparation of dPCR reaction solution (balance) | 0.02 | - | 0.2 % |
| Preparation reverse transcription reaction solution (balance) | 0.03 | - | 0.3 % |
| Partition volume | 0.64 | - | 7.3 % |
| Combined standard uncertainty | 4.76 | | 100 % |

### Laboratory 11

SM1

| Measurement uncertainty factor | (Relative) standard uncertainty | Degrees of freedom | % of combined uncertainty |
|---|---|---|---|
| Pipetting error | 0.013 | 2 | |
| Droplet volume | 0.082 | 104 | |

| | | | |
|---|---|---|---|
| Reproducibility | 0.104 | 21 | |
| Combined standard uncertainty | 0.133 | | 100 % |

SM3

| Measurement uncertainty factor | (Relative) standard uncertainty | Degrees of freedom | % of combined uncertainty |
|---|---|---|---|
| Pipetting error | 0.013 | 2 | |
| Droplet volume | 0.082 | 104 | |
| Reproducibility | 0.112 | 23 | |
| Combined standard uncertainty | 0.139 | | 100 % |

### Laboratory 12

SM1 ddPCR – QX200

| Measurement uncertainty factor | Standard uncertainty (copies/μL) | Degrees of freedom | % of combined uncertainty |
|---|---|---|---|
| Repeatability | 98 | ∞ | 36 |
| Intermediate Precision | 116 | ∞ | 43 |
| Poisson uncertainty | 58 | ∞ | 21 |
| Combined standard uncertainty | 163 | ∞ | 100 % |

SM1 cdPCR – QS3D

| Measurement uncertainty factor | Standard uncertainty (copies/μL) | Degrees of freedom | % of combined uncertainty |
|---|---|---|---|
| Repeatability | 89 | ∞ | 26 |
| Intermediate Precision | 92 | ∞ | 27 |
| Poisson uncertainty | 102 | ∞ | 30 |
| Partition volume uncertainty | 59 | ∞ | 17 |
| Combined standard uncertainty | 174 | ∞ | 100 % |

### Laboratory 13

One-step RT-dPCR (SM1)

| Measurement uncertainty factor | Relative standard uncertainty | Degrees of freedom | % of combined uncertainty |
|---|---|---|---|
| Repeatability (intra-assay) | 1.0 % | | 5.7 |
| Intermediate precision (inter-assay) | 1.5 % | | 12.7 |
| Pipette measurements | 1.5 % | | 12.7 |
| Partition volume | 3.5 % | | 69.3 |
| Combined standard uncertainty | 4.2 % | | 100 % |

Two-step RT-dPCR (SM1)

| Measurement uncertainty factor | Relative standard uncertainty | Degrees of freedom | % of combined uncertainty |
|---|---|---|---|
| Repeatability (intra-assay) | 1.7 % | | 12.5 |
| Intermediate precision (inter-assay) | 2.4 % | | 25.0 |
| Pipette measurements | 1.5 % | | 9.8 |
| Partition volume | 3.5 % | | 53.2 |
| Combined standard uncertainty | 4.8 % | | 100 % |

**Table I-1: Summary of measurement uncertainty sources considered for measurement of Study Materials 1 & 3 and Study Material 2 (RT-dPCR)**

| Type | A | A/B* | A | A/B | A/B | A/B | B | B | B | B | B | B | B | B | B |
|---|---|---|---|---|---|---|---|---|---|---|---|---|---|---|---|
| **Factor**<br>**Institute** | **RT-d/qPCR Method repeatbility** | **RT-d/qPCR intermediate precision** | **Between vial** | **Assay** | **Threshold setting** | **Sample dilution (volumetric)** | **Sample dilution (gravimetric)** | **Reaction preparation (volumetric)** | **Reaction preparation (gravimetric)** | **Homogeneity** | **Partition volume** | **RT efficiency** | **Poisson error** | **Material integrity** | **Nanodrop calibration** |
| 1 | Y | Y | Y | N/A | N/A | N/A | N/A | N/A | Y | N/A | Y | Y | Y | N/A | N/A |
| 2 | Y | Y | N | N | Y | N/A | N/A | N/A | Y | N | Y | Y | N | N | N/A |
| 2 (SM2) | Y | Y | N | N | Y | N/A | Y | N/A | Y | N | Y | Y | N | N | N/A |
| 3 | Y | Y | | Y | Y | | | | | Y | Y | | | | |
| 4 | Y | Y | Y | Y | | | | | | | Y | | | | |
| 5 | Y | Y | N | N | N | N | N | N | Y | Y | Y | N | N | N | N |
| 6 | Y | Y | Y | N | N | N | Y | Y | Y | Y | Y | N | N | N | N |
| 6 (SM2) | Y | Y | Y | N | N | N | Y | Y | Y | Y | Y | N | N | N | N |
| 7 | Y | Y | Y | N | N | Y | N | Y | | Y | Y | N | N | N | N |
| 8 | Y | Y | Y | Y | N | N | N | N | N | N | Y | N | N | N | N/A |
| 8 (SM2) | Y | Y | Y | Y | N | N | N | N | N | N | Y | N | N | N | N/A |
| 9 | Y | Y | N | Y | N | N/A | Y | Y | Y | N | Y | N | N | N | N/A |
| 9 (SM2) | Y | Y | N | Y | N | N/A | Y | Y | Y | N | Y | N | N | N | N/A |
| 10 | Y | Y | | | | | Y | | Y | | Y | | | | |
| 11 | Y | N | Y | N | Y | N | N | Y | N | N | Y | N | N/A | N | N |
| 12 ddPCR | Y | Y | | | | | | | | | | | Y | | |
| 12 cdPCR | Y | Y | | | | | | | | | Y | | Y | | |

| Type | A | A/B* | A | A/B | A/B | A/B | B | B | B | B | B | B | B | B | B |
|---|---|---|---|---|---|---|---|---|---|---|---|---|---|---|---|
| **Factor**<br><br>**Institute** | **RT-d/qPCR Method repeatability** | **RT-d/qPCR intermediate precision** | **Between vial** | **Assay** | **Threshold setting** | **Sample dilution (volumetric)** | **Sample dilution (gravimetric)** | **Reaction preparation (volumetric)** | **Reaction preparation (gravimetric)** | **Homogeneity** | **Partition volume** | **RT efficiency** | **Poisson error** | **Material integrity** | **Nanodrop calibration** |
| 13 | Y | Y | Y | N | N | N | N | Y | N | Y | Y | N | N | N | N |

**KEY**: *RT-dPCR intermediate precision: If not stated in brackets, Type A approach. Y: Yes; N: No; N/A: Not applicable.

**Table I-2 <u>Summary of Participants' Uncertainty Estimation for SM2 (HPLC, LC-MS and single molecule flow cytometry)</u>**

| **Institute** | **Factors included in uncertainty calculations** |
|---|---|
| 3 | Statistical variations (day-to-day, vial-by-vial, repeatability)<br>homogeneity and stability |
| 9 | See Appendix I part 1. |
| 10 | IDMS – each NMPs quantification including between STD, between acid hydrolysis, between measurement, preparation of sample and standard, and qNMR quantification of NMPs concentration.<br>Between NMPs<br>Density of sample |
| 11 | Uncertainties of mass and density in gravimetric dilution (combined uncertainties 0.26% to 0.28%)<br>Calculation of dilution:<br>$\Phi = 1/\left(1+\frac{m_1\,\rho_{\mathrm{storage}}}{m_{\mathrm{sample}}\,\rho_{\mathrm{storage}}}\right)\times 1/\left(1+\frac{m_2\,\rho_{\mathrm{TE}}}{m_{\mathrm{step\,1}}\,\rho_{\mathrm{storage}}}\right)\times 1/\left(1+\frac{m_3\,\rho_{\mathrm{TE}}+m_{\mathrm{stain}}\,\rho_{\mathrm{TE}}}{m_{\mathrm{step\,2}}\,\rho_{\mathrm{TE}}}\right)$<br>$\Phi$ ... volume fraction<br>$m_1$ ... mass of dillutent in first dilution step (storage buffer)<br>$\rho_{\mathrm{storage}}$ ... density of storage buffer<br>$m_{\mathrm{sample}}$ ... sample mass in first dilution step<br>$m_2$ ... mass of dillutent in second dilution step (TE buffer)<br>$\rho_{\mathrm{TE}}$ ... density of TE buffer<br>$m_{\mathrm{step\,1}}$ ... mass of first dilution used in second dilution step<br>$m_3$ ... mass of dilutent in third dilution step (TE buffer)<br>$m_{\mathrm{stain}}$ ... mass of staining solution added to third dilution step<br>$m_{\mathrm{step\,2}}$ ... mass of second dilution used in third dilution step<br><br>Uncertainty of sample volume (0.3 %); Uncertainty due to counting statistics ($\sqrt{N_i}$; 0.37% to 0.81%) |

| Institute | Factors included in uncertainty calculations |
| --- | --- |
| 3 | Statistical variations (day-to-day, vial-by-vial, repeatability)<br>homogeneity and stability |
| | Calculation of concentration measured for each dilution:<br>$$C_i = N_i \frac{1}{1-\frac{\tau_{D,i}}{t_i}} \div V_i - N_{\text{blank}} \frac{1}{1-\frac{\tau_{D,\text{blank}}}{t_{\text{blank}}}} \div V_{\text{blank}}$$<br>$N$ … measured counts<br>$\tau_D$ … integreated dead time<br>$t$ … measurement duration<br>$V$ … sample volume<br><br>The measured counts are corrected for coincidence losses using the integrated (measured) deadtime. Additionally, for each measurement counts of a blank measurement are subtracted.<br>Finally the sample concentration is calculated as weighted mean of the concentrations measured for each dilution and the uncertainty is calculated as the standard deviation of the measured concentrations:<br>$$\bar{C} = \sum_i \frac{C_i}{u(C_i)^2} \Big/ \sum_i \frac{1}{u(C_i)^2}$$ |
