## Supplementary material for "CCQM-P199: Interlaboratory comparability study of HIV-1 RNA copy number quantification": CCQM P199 supplementary file Appendix J

### APPENDIX J: Participants' Original Results as Reported

This section details any changes to the original submitted values (LGC internal ref: v1.0 dated 01 October 2019 and v2.0 dated 17 December 2019 (including Laboratory 3's Study Material 2 result)). Tables 12 to 14 of the main report correspond to the final dataset (LGC internal ref: v2.2). For clarity, only laboratories/results which were amended in the course of reporting and checking are detailed below.

**Table J-1: Study Material 1 Amendments to reported results**

| Lab ID (result) | Results reported date | Data version (NML) | *x* | *u* | *k* | *U* | *Rel U* |
|---|---|---|---|---|---|---|---|
| 5 | 8/23/2019 | 1.0 | 656.04 | *23.44* | 2 | 64.89 | 9.89% |
| 5 | 1/13/2020 | 2.0 | 656.04 | *32.44* | 2 | 64.89 | 9.89% |
| 12 | 9/19/2019 | 1.0 | 1851 | *182* | 2 | *364* | *19.7%* |
| 12 | 1/17/2020 | 2.0 | 1851 | *163* | 2 | *325* | *17.6%* |

*Red italic* indicates the change(s) to reported values or uncertainties.

**Table J-2: Study Material 2 Amendments to reported results**

| Lab ID | Results reported date | Data version (NML) | *x* | *u* | *k* | *U* | *Rel U* |
|---|---|---|---|---|---|---|---|
| 2 | 9/23/2019 | 1.0 | 6.68E+09 | *6.990E+08* | 2 | 1.40E+09 | 20.9% |
| 2* | 1/27/2020 | 2.0 | 6.68E+09 | *0.699E+09* | 2 | 1.40E+09 | 20.9% |
| 9 | 9/27/2019 | 2.0 | *6.00E+09* | *4.8E+08* | 2.00 | *9.7E+08* | 16% |
| 9 | 3/21/2021 | 2.2 | $6.33 \times 10^9$ | $0.51 \times 10^9$ | 2.00 | $1.02 \times 10^9$ | 16% |

*Number of significant figures in uncertainty (*u*) only. *Red italic* indicates the change(s) to reported values or uncertainties.
