## Supplementary material for "CCQM-P199: Interlaboratory comparability study of HIV-1 RNA copy number quantification": CCQM P199 supplementary file Appendix K

### APPENDIX K: Supplementary and follow-up results

This section details additional results which were shared or relate to experiments performed following discussion of the study results at NAWG for troubleshooting purposes.

#### K.1 Additional data: Study Material 2 purity analysis

Additional data informing the fragment size profile of Study Material 2 was shared by Laboratory 9 (Figure K.1) and by Laboratory 3 (Figure K.2) from their analysis / evaluation of the material during their study participation.

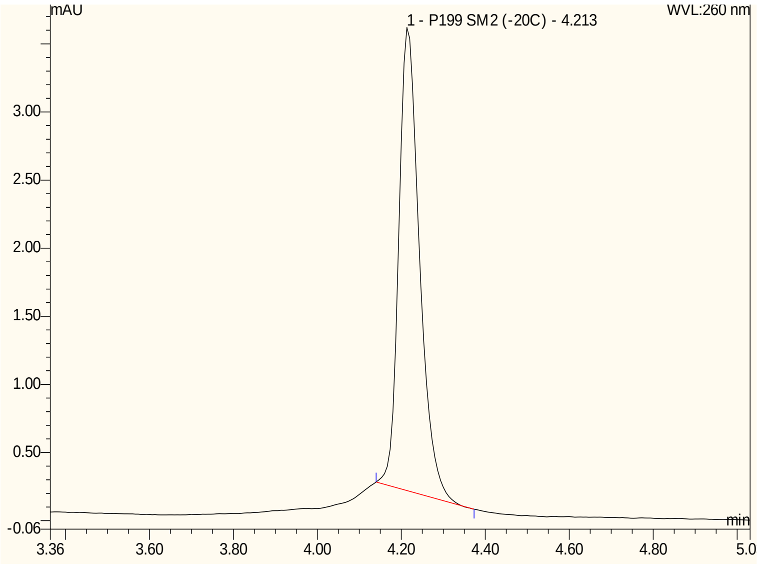


**Figure K-1: HPLC Chromatogram (Laboratory 9) participant analysis of Study Material 2. Y-axis shows relative intensity of UV signal (260 nm; milli-Absorbance Units (mAU)); x-axis shows time (min).**

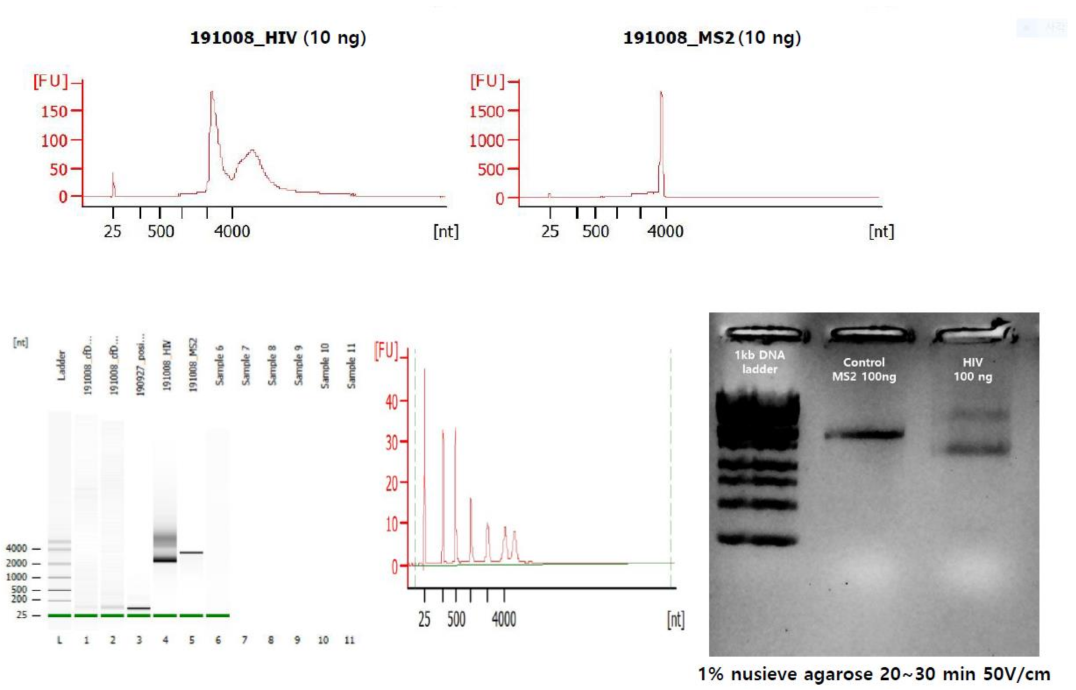


**Figure K-2: Capillary electrophoresis and gel electrophoresis (Laboratory 3) participant analysis of Study Material 2. MS2 RNA was used as control.**

Figure K.3 shows follow-up analysis of Study Material 2 performed by Laboratory 10 using UPLC in combination with S1 nuclease, which is specific for single strand nucleic acids, to confirm that size-based impurities identified by UPLC consisted of RNA (rather than DNA).

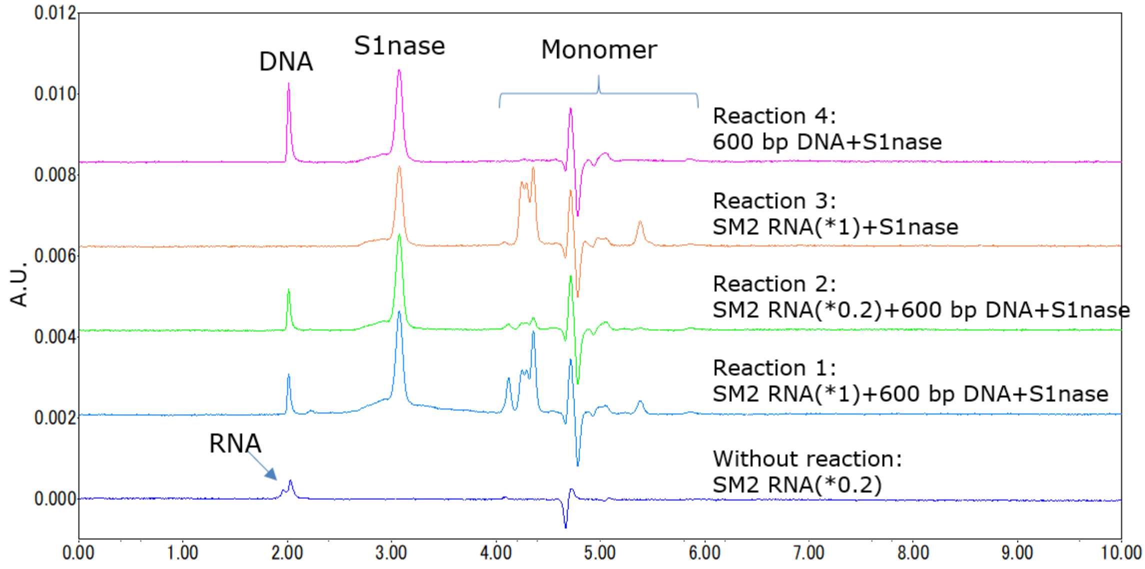


**Figure K-3: Results of S1 nuclease-digestion of Study Material 2 (Laboratory 10).**
Study Material 2 (SM2) was incubated with a double-stranded DNA control (600 bp DNA (NMIJ CRM)) with S1 nuclease (Takara bio Japan) at 37 °C for 2 h. Digested samples were analysed by UPLC (as detailed in Appendix H, Table H-7). RNA was degraded and monomer peaks were observed in Reaction 1 to 3. DNA was not degraded

and DNA peak was observed in Reaction 1, 2 and 4. These results lead to the conclusion that the main peak and impurity peak in Study Material 2 were derived from RNA.

### K.2. Follow-up results

Additional results for Study Material 1 were generated after discussion of the participant datasets by Laboratory 12 as part of troubleshooting the outlying results during the study.

**Table K-1: Follow-up results (Study Material 1)**

| **Lab ID** | Laboratory 12 |
|---|---|
| **MATERIAL** | **MEASURAND** (UNIT) |
| **STUDY MATERIAL 1** | **RNA copy number concentration (*gag* gene)** (copies/µL) |
| Value (*x*) | 945 |
| Standard uncertainty (*u*) | 143 |
| Coverage factor (*k*) | 2 |
| Expanded uncertainty (*U*) | 286 |
| Relative expanded uncertainty (Rel *U*) | 30.2 % |

Date of results: May 08, 2020. Analysis was performed with High capacity RNA to cDNA kit (Thermo Scientific (P/N 4387406)) and Laboratory 12's previously applied dPCR assay (Appendix H). Two independent experiments were performed across two days with different units of material (RT $n$ = 3, dPCR $n$ = 2 per cDNA).
