## Supplementary material for "CCQM-P199: Interlaboratory comparability study of HIV-1 RNA copy number quantification": CCQM P199 supplementary file Appendix L

### APPENDIX L: Additional statistical analysis

This section details results of statistical analysis.

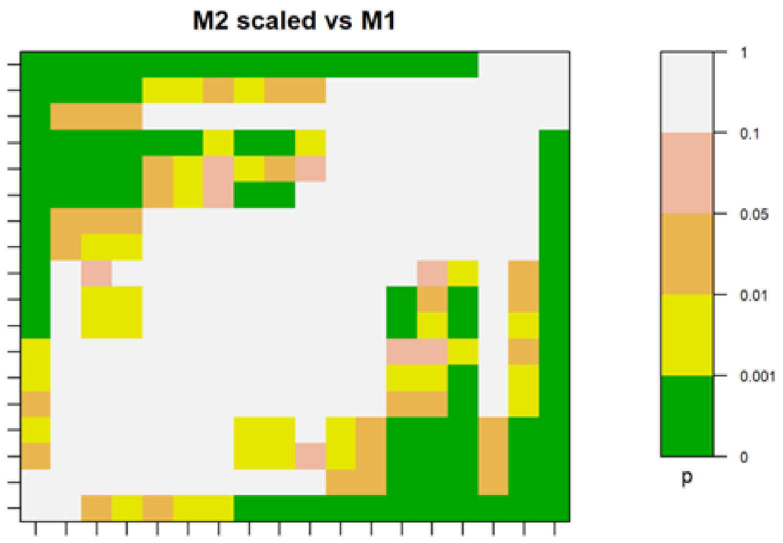


**Figure L-1: Pairwise comparison of laboratories: results of consistency analysis of M2 scaled to M1 using the associated gravimetric dilution factor**

A previous iteration of the gravimetric uncertainty value (rel u = 0.86 %) was used to calculate the M2 (scaled) uncertainties.
